# Gut microbiota within-host evolution enforces colonization resistance against enteric infection

**DOI:** 10.64898/2026.03.27.714693

**Authors:** Marta Salvado Silva, Simon Woelfel, Claudia Eberl, Denise Medeiros Selegato, Abilash Durai Raj, Philipp C. Münch, Birte K. Jung, Hélène Omer, Michael Hellwig, Lisa Osbelt, Bidong Nguyen, Silvia Bolsega, Susanne Wudy, Debora Garzetti, Diana Ring, Monica S. Matchado, Marla Gaissmaier, Alexandra von Strempel, Saib Hussain, Lea Fuchs, Marijana Basic, Christina Ludwig, Jürgen Lassak, Emma Slack, Till Strowig, Alice C. McHardy, Wolf-Dietrich Hardt, Michael Zimmermann, Dirk Haller, Bärbel Stecher

## Abstract

Limited resource availability in the gut promotes competitive interactions between bacteria, which drive adaptive within-host evolution (*1–3*). While adaptive evolution of bacterial communities has been increasingly studied in the recent years (*4–7*), its functional implications for host physiology remain unknown. Here, we show that within-host evolution of the human commensal *Enterococcus faecalis* boosts colonization resistance to enteric *Salmonella enterica* serovar Typhimurium (*S*. Typhimurium) infection. During gut colonization, *E. faecalis* evolves the ability to metabolize fructoselysine, an abundant Amadori rearrangement product generated by thermal food processing. The depletion of this diet-derived nutrient prevents *S*. Typhimurium colonization by restricting an essential resource. This protective mechanism was conserved across independent mouse colonies and arises via diverse evolutionary trajectories, including nucleotide polymorphisms, gene amplifications, and a horizontal gene transfer event. Additionally, analysis of *E. faecalis* isolates from human infants revealed that adaptation to fructoselysine availability occurs in a diet-dependent manner. Isolates from infants fed with fructoselysine-rich formula were able to utilize fructoselysine, whereas those from infants fed with fructoselysine-poor breast milk were not. Conclusively, our results identify an inherent microbiome-driven self-healing mechanism, wherein bacterial evolution restores colonization resistance against enteric pathogens through evolved nutrient depletion. Understanding these evolutionary dynamics will inform microbiome-targeted approaches to prevent and treat infectious diseases by harnessing adaptive bacterial metabolism.

## Main

Microbial colonization of the human intestine begins at birth and follows complex eco-evolutionary principles (*8–10*). Competition for available resources plays a key role in the microbial assembly process (*11–13*). While the gut microbiota exhibits remarkable strain-level stability over time (*14*), individual bacterial lineages can undergo significant evolutionary changes (*14–16*). Natural selection acting on *de novo* mutations drives strain diversification, leading to lineage coexistence or competitive exclusion based on environmental pressures (*17*). The microbial genes under selection suggest that diet, drugs, immune defenses, and other microbes are major factors driving adaptive microbial evolution in the gut (*17–19*). While the fitness-promoting effects of these mutations on bacteria are largely well understood, the functional consequences for the host remain unknown.

A key function of the gut microbiome is colonization resistance, primarily achieved through nutrient blocking, where enteric pathogens are excluded by competing for shared nutrient sources with resident bacteria (*20, 21*). Within-host evolution fine-tunes bacterial metabolism to optimize nutrient acquisition and depletion(*22*). Theoretical models predict that evolution-mediated priority effects could strengthen colonization resistance against pathogens (*23*). Based on these assumptions, we hypothesized that microbiota-driven within-host evolution enhances colonization resistance by increasing the consumption of limiting nutrients, thereby restricting pathogen access to essential resources (**Fig. 1A**).

**Fig. 1:**
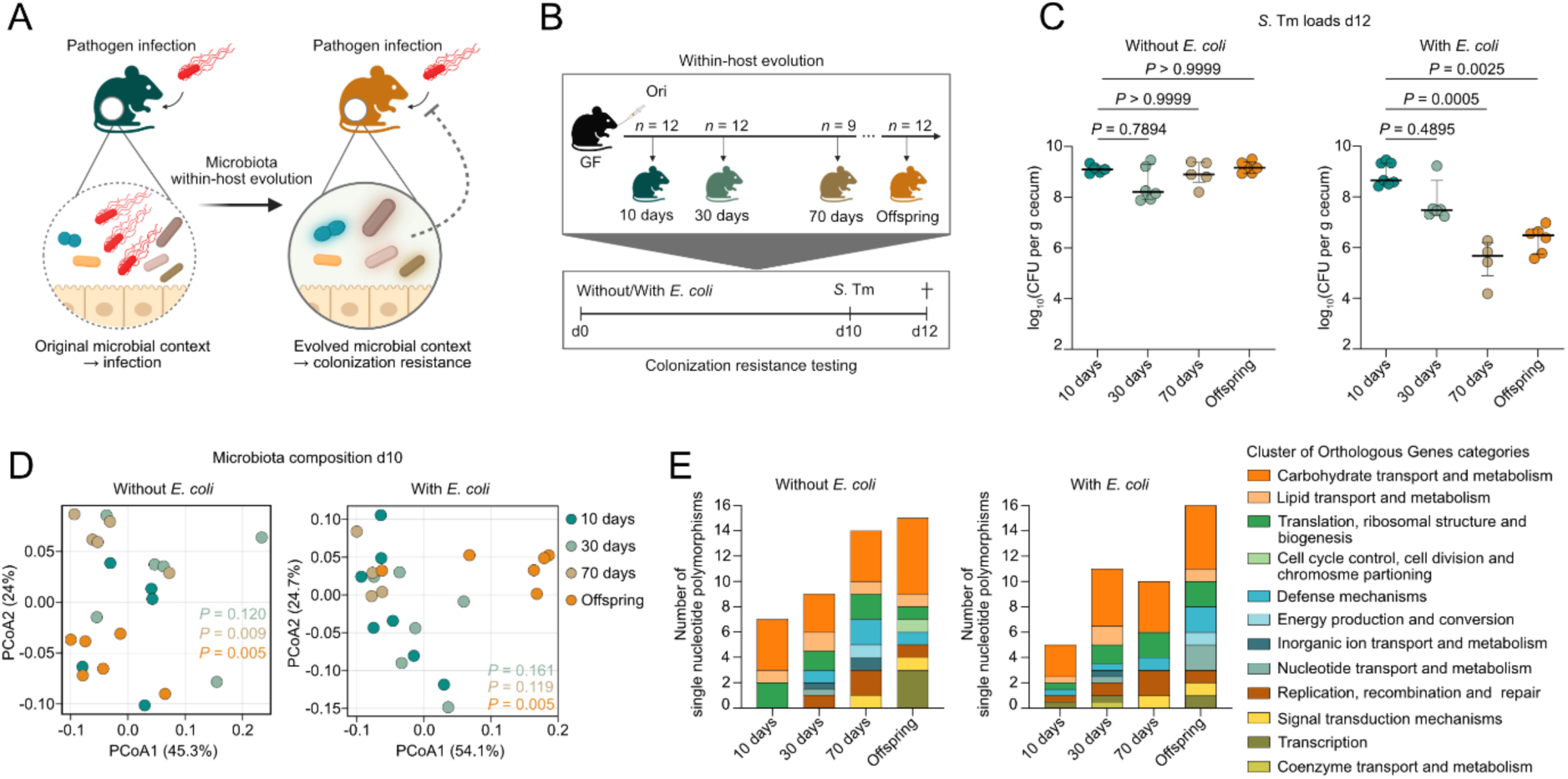
Resistance against *S.* Typhimurium colonization is an adaptable trait of a microbial community. **(A)** Working hypothesis. Microbiota within-host evolution establishes a protective microbial context and leads to colonization resistance against enteric pathogens. **(B)** Longitudinal assessment of colonization resistance against *S*. Typhimurium (*S*. Tm) in mice colonized with the OMM^12^ microbial community. Germ-free mice were colonized with the original OMM^12^ community(*24*), freshly assembled from cryo-stocks. Mice were maintained for 10, 30, or 70 days and then inoculated with *E. coli* or PBS, followed by infection with *S*. Tm 10 days later, to assess *E. coli*-mediated colonization resistance. Offspring of the original OMM^12^-inoculated mice were likewise treated with *E. coli* or PBS and infected with *S.* Tm. All mice were sacrificed 2 days after *S.* Tm infection (d12). **(C)** Cecum *S*. Tm loads on day two post-infection (d12). Data represent medians with interquartile ranges, and statistical analyses were performed using Dunn’s post-hoc tests. The detection limit is 10 CFU per g. **(D)** Fecal microbiota composition on the day of *S*. Tm infection (d10). Principal coordinate analyses were based on the Bray-Curtis dissimilarity distance matrices of relative OMM^12^ abundance profiles at different colonization timepoints. Statistical analyses were performed using PERMANOVA with Bonferroni correction, comparing the 30 days, 70 days, and offspring timepoints to the 10 days timepoint, respectively. **(E)** The number of coding sequence single nucleotide polymorphisms in OMM^12^ microbiota per Clusters of Orthologous Genes (COG) category in feces metagenomes on the day of *S*. Tm infection (d10). One or two mice were used per timepoint.

### Protection against *S*. Typhimurium is an adaptable trait of a microbial gut community

To test this hypothesis, we utilized the well-established OMM^12^ model of intestinal colonization resistance, which is based on a twelve-member synthetic bacterial community (Oligo-mouse-microbiota 12; OMM^12^) (*24*) that stably colonizes the gut of germ-free mice (*25*). In this model, *E. coli* Mt1B1 provides colonization resistance against *S.* Typhimurium in a context-dependent manner by depleting the sugar alcohol galactitol, which would otherwise fuel pathogen growth (*20*). Importantly, the OMM^12^ further depletes additional nutrients that are not utilized by *E. coli* but would fuel *S*. Typhimurium growth if not depleted.

We inoculated germ-free mice with original OMM^12^ cultures (*26*), maintained them for 10, 30, or 70 days, and subsequently colonized them with *E. coli* Mt1B1 before orally challenging them with *S.* Typhimurium (**Fig. 1B**). Stably colonized mice born from OMM^12^-colonized parents (offspring) served as controls. Quantifying *S*. Typhimurium cecum loads following pathogen challenge revealed that *E. coli* only provided colonization resistance in mice colonized with OMM^12^ for over 70 days (**Fig. 1C**) but not in those with shorter-term colonization, despite similar overall OMM^12^ community composition (**Fig. 1D; Fig. S1A**). No overt symptoms of gut inflammation were observed (**Fig. S1B**) and experiments in *Rag2*^-/-^mice, which lack mature T- and B-lymphocytes, excluded a significant role for adaptive immunity in the underlying resistance mechanism (**Fig. S1C-F**). Accumulation of single nucleotide polymorphisms in various OMM^12^ members indicated rapid evolutionary adaptation over the course of 70 days (**Fig. 1E**), including a notable enrichment of single nucleotide polymorphisms in genes associated with carbohydrate transport and metabolism. Based on these findings, we concluded that within-host microbiota evolution occurred during the investigated time frame and that bacterial carbon metabolism is a target of selection.

### Transplantation of an evolved microbiota establishes a protective microbial context against *S.* Typhimurium

To establish causality between microbiota evolution and colonization resistance, we inoculated germ-free mice with fresh OMM^12^ cultures (*24*) or cecum content from stably colonized OMM^12^ mouse lines, which harbor an evolved microbiota (Evo HAN, Evo MUC2, Evo MUC; **Fig. 2A-B**, **Fig. S2A**). We then analyzed colonization resistance against *S.* Typhimurium. Mice colonized with evolved microbiota showed resistance to *S.* Typhimurium colonization, as indicated by reduced cecum *S.* Typhimurium loads (**Fig. 2C**, **Fig. S2B**). On the day of *S.* Typhimurium challenge, we identified distinct metabolite profiles in cecal washes from mice colonized with original OMM^12^ or evolved Evo MUC2. These changes in intestinal nutrient availability suggest alterations in microbiota metabolic activities during within-host evolution (**Fig. 2D**). Metabolome changes were accompanied by shifts in microbiota composition, including increased *E. faecalis* absolute abundance in mice colonized with evolved OMM^12^ (**Fig. 2E**, **Fig. S2C-D**). These results demonstrate that colonization resistance is mediated by microbiota within-host evolution, which drives changes in intestinal metabolite profiles and results in increased *E. faecalis* loads.

**Fig. 2:**
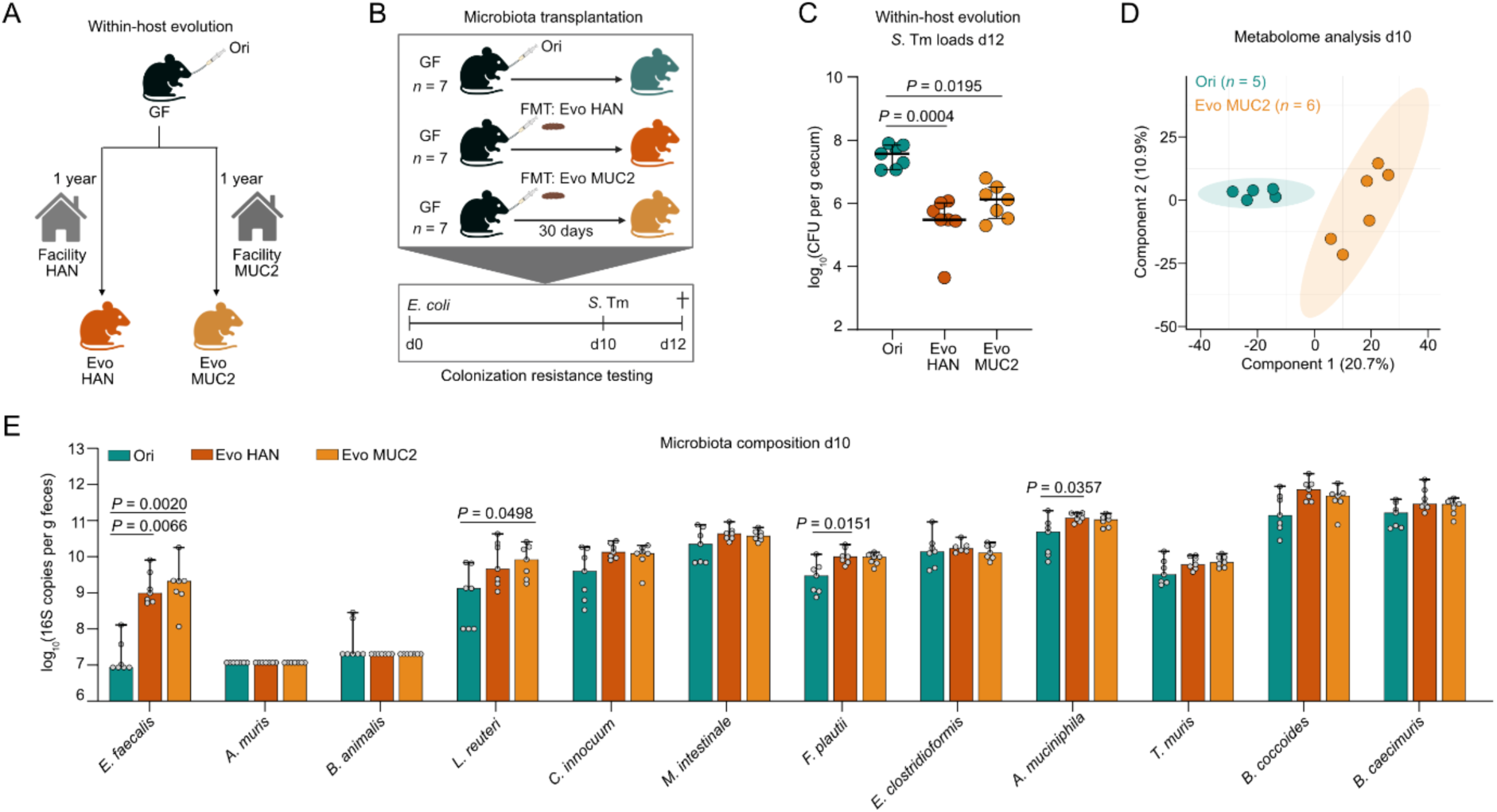
Transplantation of evolved OMM^12^ microbiota is sufficient to mediate a protective microbial context against *S.* Typhimurium. **(A)** Microbiota within-host evolution scheme. Germ-free mice were colonized with an OMM^12^ community freshly assembled from single species stocks. Mice were housed in two different animal facilities, resulting in two distinct mouse colonies (Evo HAN, Evo MUC2) harboring separately evolved OMM^12^ communities. **(B)** Assessment of colonization resistance against *S*. Typhimurium (*S*. Tm) in mice colonized with the original OMM^12^ community(*24*) or transplanted with evolved OMM^12^ communities (Evo HAN, Evo MUC2). Colonization resistance was assessed as described in Fig. 1B, but all mice were inoculated with *E. coli*. **(C)** Cecum *S*. Tm loads on day 2 post-infection (d12). Data represent medians with interquartile ranges, and statistical analysis was performed using a Dunn’s post-hoc test. The detection limit is 10 CFU per g. **(D)** Principal component analyses of cecum metabolomes from mice colonized as described in b with either the original(*26*) or evolved (Evo HAN, Evo MUC2) OMM^12^ communities. Mice (*n* = 5 per group) were sacrificed on day 10 post-*E. coli* colonization or on day two post-*S*. Tm infection (d12). **(E)** Fecal microbiota composition on the day of *S.* Tm infection (d10) in the mice shown in (B) and (C). Absolute abundances were determined via qPCR, quantifying species-specific 16S rRNA gene copies per g feces. Data represent medians with 95% confidence intervals. Statistical analysis was performed using a Dunn’s post-hoc test, comparing the original with each evolved OMM^12^ community. Only *P*-values below 0.05 are shown.

### Transplantation of evolved *E. faecalis* strains establishes a protective microbial context against *S.* Typhimurium

*E. faecalis* is a prevalent human gut commensal known for its high genome plasticity and plays a critical role in ecosystem structuring (*27*) and modulation of colonization resistance (*28, 20*). Therefore, we reasoned that increased *E. faecalis* loads in our mouse model result from genetic alterations during within-host evolution and could underlie protection against *S*. Typhimurium. To delineate the *E. faecalis* mutations associated with colonization resistance, we isolated and sequenced evolved *E. faecalis* from the cecum of stably colonized OMM^12^ mice housed in different animal facilities (Evo HAN, Evo MUC2; *n* = 11 isolates per facility; **Fig. 3A**). By mapping sequence reads to the ancestral *E. faecalis* KB1 reference genome, we identified several unique single nucleotide polymorphisms across the evolved isolates. Phylogenetic analysis of evolved *E. faecalis* isolates revealed distinct clusters for each mouse line and varying single nucleotide polymorphism counts, which were linked to mutation rate polymorphisms in evolved isolates (**Fig. 3B**, **Fig. S3A-B**). Notably, mutation rate polymorphisms in the gut microbiome are thought to arise from strong selection prompted by competitive interactions among microbiota (*29*). *E. faecalis* from the Evo MUC2 mouse colony contained an integrative conjugative element (ICE) (*2*), which was horizontally acquired from another OMM^12^ member – *Blautia coccoides* YL58 (**Fig. S3C)**. ICEs are mobile genetic elements commonly found in the human gut microbiota, playing a crucial role in bacterial adaptive evolution by enabling the spread of beneficial traits between different bacterial species (*30*). The ICE in *E. faecalis* isolates is 156 kb long and encodes 262 genes involved in conjugation, defense mechanisms, metabolism, and other functions (**Fig. S3D**).

**Fig. 3:**
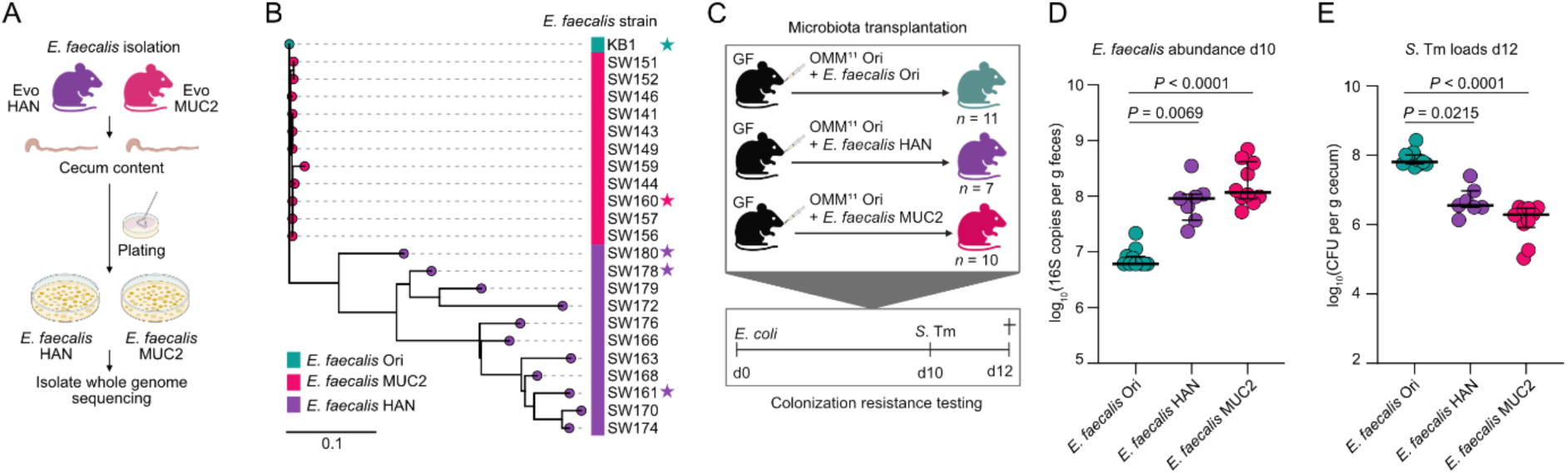
Transplantation of evolved *E. faecalis* is sufficient to mediate a protective microbial context against *S.* Typhimurium. **(A)** *E. faecalis* isolation scheme. Evolved *E. faecalis* were isolated from the cecal contents of long-term colonized OMM^12^ mice (Evo HAN, Evo MUC2) from two different animal facilities. The isolates were subjected to short-read whole genome sequencing (*n* = 1 mouse per facility). (**B)** Single nucleotide polymorphism-based phylogenetic tree of evolved *E. faecalis* isolates (*n* = 11 isolates per mouse) and original *E. faecalis* KB1. The tree was created using the Random Accelerated Maximum Likelihood method, and branch lengths represent the substitutions per variable site (*n* = 861). A polymorphism allele frequency cut-off of 0.8 was applied. Stars highlight the strains used in (C). (**C)** Assessment of colonization resistance against *S*. Typhimurium (*S*. Tm) by original or evolved *E. faecalis*. Germ-free mice were colonized with the original OMM^11^-*E. faecalis* community, supplemented with either original *E. faecalis* KB1 (*E. faecalis* Ori) or evolved *E. faecalis* isolates. The evolved isolates SW161, SW178, and SW180, or SW160 represent *E. faecalis* HAN and *E. faecalis* MUC2, respectively. Colonization resistance was assessed as described in Fig. 2B. (**D)** *E. faecalis* abundance on the day of *S.* Tm infection (d10). Absolute abundances were determined via qPCR, quantifying species-specific 16S rRNA gene copies per g feces. The abundances of remaining OMM^12^-species are provided in Fig. S3. (**E)** Cecum *S*. Tm loads on day 2 post-infection (d12). Data represent medians with interquartile ranges. The detection limit is 10 CFU per g. (**D** to **E)** Statistical analysis was performed using a Dunn’s post-hoc test.

To test whether *E. faecalis* within-host evolution mediates protection against *S*. Typhimurium, we assembled original OMM^11^ communities lacking *E. faecalis* and supplemented them with original *E. faecalis* KB1 or with evolved *E. faecalis* isolates (**Fig. 3C**). Isolates were selected from two independent mouse lines and either represent *E. faecalis* phylogenetic clusters (SW161, SW178, SW180: Evo HAN) or harbor the identified ICE (SW160: Evo MUC2). We inoculated germ-free mice with the assembled communities and assessed colonization resistance against *S*. Typhimurium. Consistent with our previous experiments, we found an increased absolute abundance of *E. faecalis* in mice colonized with evolved isolates on the day of *S*. Typhimurium infection (**Fig. 3D**, **Fig. S3E**). Strikingly, mice colonized with evolved *E. faecalis* isolates from either mouse line, Evo HAN or Evo MUC2, were protected from *S*. Typhimurium colonization (**Fig. 3E**), suggesting that *E. faecalis* within-host evolution results in increased loads and mediates colonization resistance against *S.* Typhimurium.

We then isolated additional OMM^12^ species from the same animals to elucidate whether this role was unique to *E. faecalis* or could also be mediated by other evolved OMM^12^ species (**Fig. S4A-B).** We assembled two semi-evolved communities in which selected species were represented by either the original strain or evolved isolates (**Fig. S4C**). Semi-evolved communities included evolved isolates from five OMM^12^ species, plus either original *E. faecalis* KB1 or the evolved isolate SW160. When analyzing colonization resistance against *S*. Typhimurium in mice colonized with these communities, we found that only the semi-evolved community with evolved *E. faecalis* SW160 – not the equivalent containing original *E. faecalis* KB1 – provided protection against pathogen colonization (**Fig. S4D-F**). Thus, colonization resistance against *S*. Typhimurium relies on *E. faecalis* within-host evolution.

### Evolved *E. faecalis* isolates consume fructoselysine and provide colonization resistance by niche preemption

Microbial members can provide colonization resistance by producing inhibitory substances or by competing for resources (*31*). We found no evidence of *S.* Typhimurium growth inhibition by *E. faecalis* SW160 (**Fig. S5A**). However, in contrast to original *E. faecalis* KB1, evolved *E. faecalis* SW160 could grow in cecal content medium obtained from OMM^11^ mice, lacking *E. faecalis* (**Fig. 4A**). This suggests that within-host evolution facilitates *E. faecalis* growth on nutrient sources that may underly competitive interactions with *S*. Typhimurium in the gut. Comparative proteome analysis of *E. faecalis* KB1 and SW160 revealed increased production of enzymes encoded in the fructoselysine/glucoselysine utilization operon (**Fig. 4B-D**, **Fig. S5B-E).** In line with this finding and in contrast to original *E. faecalis* KB1, *E. faecalis* SW160 and several other evolved *E. faecalis* isolates from Evo HAN and Evo MUC2 mice could grow in medium containing fructoselysine as sole carbon source (**Fig. 4D**, **Fig. S5F-G**). Fructoselysine is an Amadori rearrangement product that is generated by thermal food processing and is highly abundant in human diet (*32, 33*). It is worth noting that this operon is not encoded on the ICE transferred from *B. coccoides* YL58 but is part of the *E. faecalis* genome. Given that fructoselysine is a diet-derived nutrient potentially supporting *S.* Typhimurium colonization in the gut, we hypothesized that *E. faecalis* SW160 may limit *S.* Typhimurium colonization by depleting this compound. To test this, we analyzed the metabolomes of cecal washes from OMM^12^ mice harboring either the original *E. faecalis* KB1 or evolved *E. faecalis* SW160, before and after *S.* Typhimurium infection (**Fig. 4E-H**, **Fig. S5H-I**). We found that cecal fructoselysine concentrations were reduced in mice colonized with *E. faecalis* SW160 on the day of *S*. Typhimurium infection, suggesting that this evolved strain depleted intestinal fructoselysine (**Fig. 4G**). Following *S.* Typhimurium infection, all groups showed reduced fructoselysine levels, indicating that *S.* Typhimurium also consumed fructoselysine in the gut (**Fig. 4H**). Similarly, we observed higher fructoselysine levels in the cecal metabolomes of mice colonized with original OMM^12^ versus evolved OMM^12^ (Evo MUC2) communities (**Fig. S5J**). This suggests that our reductionist approach—simplifying complex evolved communities by combining original and evolved strains—is effective in determining the contributions of individual species to the observed colonization resistance phenotype. To test if fructoselysine depletion by evolved *E. faecalis* increases colonization resistance against *S*. Typhimurium, we performed infection experiments using isogenic *E. faecalis* SW160 and *S.* Typhimurium mutants deficient in fructoselysine utilization. These experiments verified that *S.* Typhimurium colonization of the mouse gut relies on fructoselysine utilization and that evolved *E. faecalis* blocks *S.* Typhimurium infection by preempting the fructoselysine niche (**Fig. 4I, Fig. S5K-M**).

**Fig. 4:**
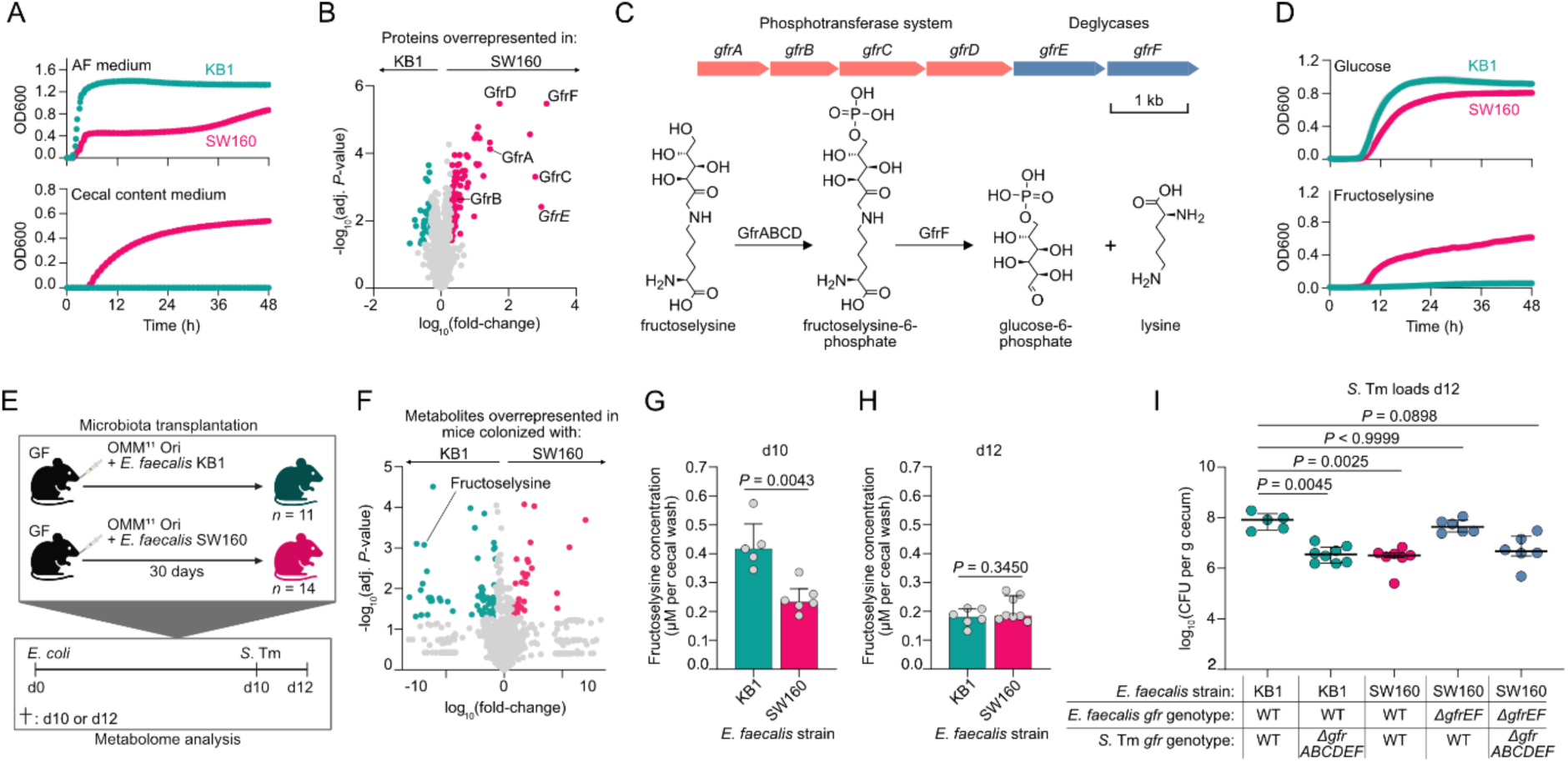
Evolved *E. faecalis* isolates consume fructoselysine and provide colonization resistance by niche preemption. **(A)** Growth curves of original *E. faecalis* KB1 and evolved *E. faecalis* SW160 in AF or cecal content liquid medium. Cecal content was obtained from a mouse long-term colonized with the OMM^11^-*E. faecalis* community. **(B)** Volcano plot showing proteins that are overrepresented in *E. faecalis* KB1 (turquoise) or SW160 (magenta). The bacteria were cultivated in AF liquid medium and their proteomes analyzed during the late stationary growth phase. Significance cut-offs were log_10_(2) and -log_10_(0.05) for fold-changes and Benjamini-Hochberg adjusted t-test *P*-values, respectively. Proteins encoded by the fructoselysine operon are highlighted. **(C)** Scheme of the *E. faecalis* fructoselysine utilization operon. *gfrABCD* and *gfrEF* encode components of the phosphotransferase system EIIA/EIIB/EIIC/EIID and deglycases, respectively. The metabolic pathway illustrates the biodegradation of fructoselysine to glucose-6-phosphate and lysine. **(D)** Results corresponding to b but using minimal medium supplemented with glucose or fructoselysine as the sole carbon source. **(E)** Experimental scheme for mouse intestinal metabolome analysis. Germ-free mice were colonized with the original OMM^11^-*E. faecalis* community and either the original *E. faecalis* KB1 or the evolved *E. faecalis* SW160. After inoculation with *E. coli*, mice were infected with *S*. Typhimurium (*S*. Tm) 10 days later. Metabolomes of cecal washes were analyzed on the day of *S.* Tm infection (d10; KB1: *n* = 5; SW160: *n* = 6) or two days post-infection (d12; KB1: *n* = 6; SW160: *n* = 8). **(F)** Volcano plot showing metabolites that are overrepresented in mice colonized with *E. faecalis* KB1 (turquoise) or SW160 (magenta) on the day of *S*. Tm infection (d10). The FDR-adjusted *P*-value significance cut-off was -log_10_(2). Additional data are available in Fig. S5. (**G** to **H)** Fructoselysine abundance in cecal washes on the day of *S*. Tm infection (d10; G) and on day two post-infection (d12; H). Data represent medians with interquartile ranges, and statistical analyses were performed using Mann-Whitney *U* tests. **(I)** Cecum *S*. Tm loads on day two post-infection (d12) in germ-free mice colonized with the original OMM^11^-*E. faecalis* community and either original *E. faecalis* KB1 (turquoise), evolved *E. faecalis* SW160 (magenta), or a fructoselysine utilization-deficient *E. faecalis* SW160 mutant (*ΔgfrEF;* blue). Mice were inoculated with *E. coli*, infected with either wild-type *S*. Tm (WT) or a fructoselysine utilization-deficient *S*. Tm mutant (*ΔgfrABCDEF*) 10 days later, and euthanized on day two post-infection (d12; similar to Fig. 2B). Data represent medians with Interquartile ranges, and statistical analysis was performed using a Dunn’s post-hoc test. The detection limit is 10 CFU per g. Cecum fructoselysine concentrations can be found in Fig. S5.

To evaluate the conservation of this mechanism, we isolated *E. faecalis* from three additional stably colonized OMM^12^ mouse lines housed in different animal facilities (Evo BS, Evo ZUC, Evo MUC). Notably, the Evo MUC mouse line was used for the longitudinal evolution experiments displayed in **Fig. 1** (**Fig. S6A**). We identified single nucleotide polymorphisms in the fructoselysine utilization operon in isolates from two of these mouse lines, located in genes encoding ABC phoshotransferase system components (*gfrA*, *gfrC*) or the fructoselysine-degylcase (**Fig. S6C**). Similarly, we found polymorphisms in *gfrA* and the operon promoter region in the genome of *E. faecalis* SW178, which was isolated from the Evo HAN mouse line, utilizes fructoselysine, and confers colonization resistance (**Fig. 3D**, **Fig. S5G**, **Fig. S6C**). Interestingly, we observed a 10- to 20-fold copy number increase of the fructoselysine utilization operon in all *E. faecalis* isolates from the Evo MUC mouse line (**Fig. S6D**). Such gene amplifications enable bacteria to rapidly adapt to a changing environment by increasing gene expression and are a key evolutionary process under selective pressure (*34*). Given that colonization resistance in this mouse line increased with the duration of microbiota colonization, we aimed to evaluate whether the onset of colonization resistance coincided with the emergence of the fructoselysine operon amplification (**Fig. 1**). Therefore, we screened for amplification in metagenomes from these animals sampled on the day of *S*. Typhimurium infection by mapping metagenome reads to the *E. faecalis* KB1 reference genome. The operon coverage fold-change relative to the reference genome was elevated in metagenomes from the 30 days, 70 days, and offspring groups, correlating with the level of colonization resistance against *S*. Typhimurium (**Fig. S6E**). Collectively, our findings suggest that increased fructoselysine consumption, which enforces colonization resistance against *S*. Typhimurium, is mediated by the acquisition of single nucleotide polymorphisms in genes of the fructoselysine utilization operon or by amplification of this operon in some *E. faecalis* isolates. Similar to *E. faecalis* isolates from Evo HAN and Evo MUC2 mouse lines, isolates from Evo BS, Evo ZUC, and Evo MUC2 mice exhibited enhanced levels of fructoselysine utilization enzymes and, to some extent, significant growth in OMM^11^ cecal content medium, as well as in medium containing fructoselysine as the sole carbon source (**Fig. S6F-J**). In conclusion, we isolated *E. faecalis* from five independent OMM^12^ mouse lines housed in four different animal facilities and identified convergent evolution towards increased fructoselysine utilization via diverse evolutionary trajectories.

### *E. faecalis* isolates from human infants show diet-dependent variations in fructoselysine utilization

In the human gut microbiome, the capacity for fructoselysine utilization is prevalent (*35, 36*). However, not all strains harboring the relevant genes are able to utilize fructoselysine, indicating that gene expression is tightly regulated (*35*). Fructoselysine is abundant in the human diet, particularly in ultra-processed foods, including infant formula (*37, 38*). Consequently, formula-fed infants exhibit higher fructoselysine levels in their stool than breast-fed infants (*39*). Given the high prevalence and abundance of *E. faecalis* in the infant gut, we reasoned that formula- and breast-fed infants provide a suitable cohort to investigate *E. faecalis* adaptive evolution in fructoselysine-rich and -poor gut environments.

We isolated *E. faecalis* from three-month-old infants who were fed either formula or breast milk (**Fig. 5A**). Stool samples were collected from a previously published randomized controlled trial(*39*) (**Fig. S7A**). In this study cohort, formula-fed infants exhibited higher microbiome richness than breast-fed infants, while *E. faecalis* abundance was comparable in both groups (**Fig. S7B-D**). Importantly, fructoselysine concentrations were higher in formula-fed infants compared to breast-fed infants (*39*). Since *E. faecalis* gut colonization typically begins at birth, these isolates were exposed to either a fructoselysine-low (breast milk) or fructoselysine-high (formula) environment for three months – a comparable timeframe during which we detected *E. faecalis* adaptive evolution towards fructoselysine utilization in our animal model. Remarkably, we found that only isolates from formula-fed infants could grow in a medium containing fructoselysine as the sole carbon source (**Fig. 5B-C**, **Fig. S7F**), which was not observed in isolates from breast-fed infants. Notably, *E. faecalis* isolates from both sources encoded the fructoselysine operon (**Fig. S7E**). Whole genome long-read sequencing and *de novo* assembly of selected isolates from formula-fed infants revealed that these isolates are phylogenetically distinct, suggesting that the ability to consume fructoselysine can arise independently in different genomic backgrounds (**Fig. 5B-C**). Our results demonstrate that human enterococci possess the capacity to utilize fructoselysine in a diet-dependent manner, suggesting that within-host evolution towards fructoselysine consumption also occurs in the human gut.

**Fig. 5:**
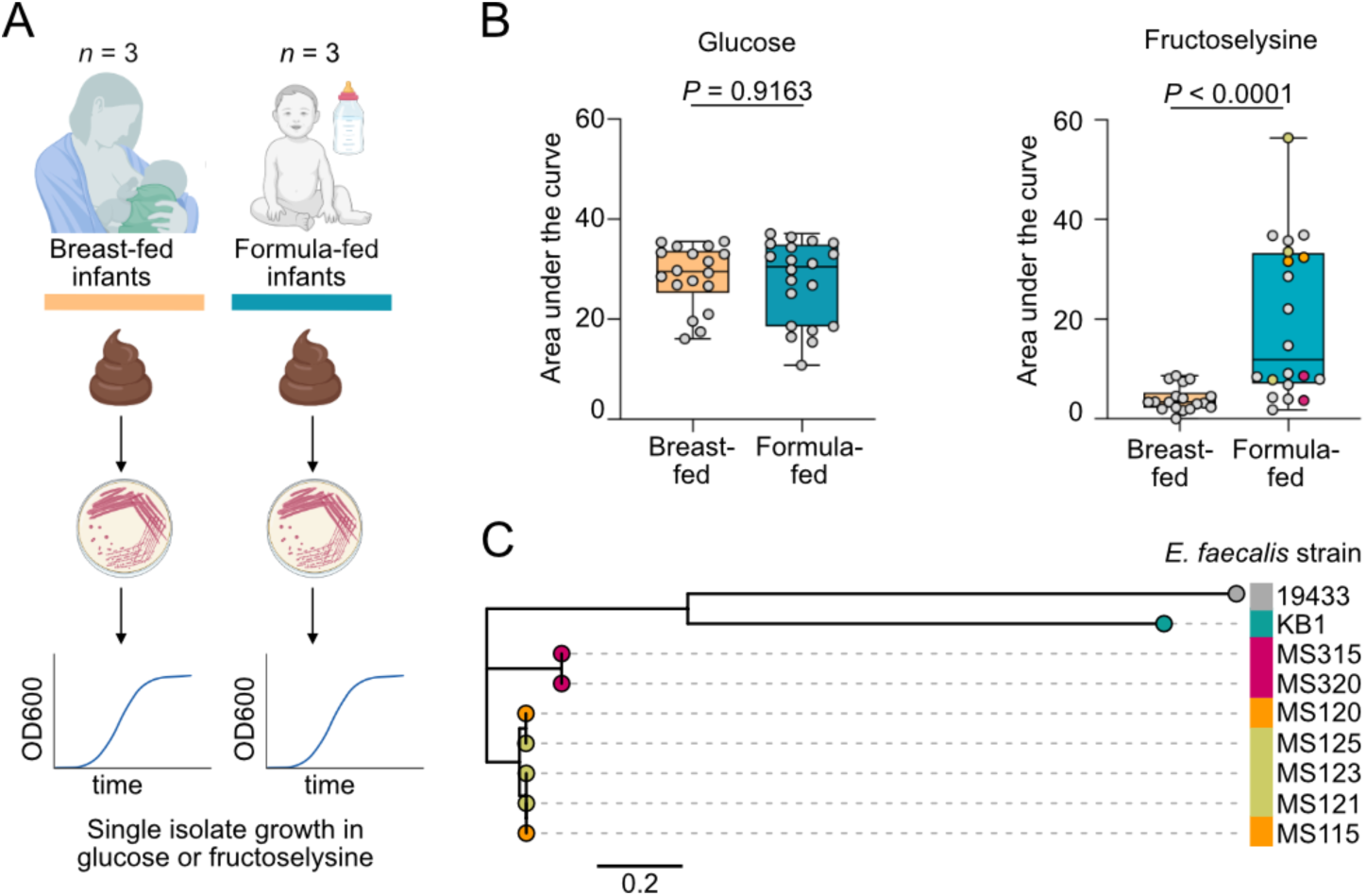
Increased prevalence of fructoselysine-proficient *E. faecalis* in formula-fed neonates. **(A)** *E. faecalis* isolation scheme. *E. faecalis* was isolated from the stool of three-month-old breast- or formula-fed infants. Isolate growth was assessed in minimal medium with glucose or fructoselysine as the sole carbon source. *n* = 6-7 isolates per infant. **(B)** Box plots showing the area under the curve based on the growth of *E. faecalis* isolates in minimal medium with glucose or fructoselysine as the sole carbon source (breast-fed: *n* = 18; formula-fed: *n* = 20). Colored data points correspond to whole genome-sequenced isolates in c. Statistical analyses were performed using Mann-Whitney *U* tests. **(C)** Whole genome-based phylogenetic tree of select *E. faecalis* isolates (*n* = 7) from formula-fed infants. *E. faecalis* KB1 and ATCC19433 were included as phylogenetic reference points. The tree was created using the Random Accelerated Maximum Likelihood method, and branch lengths represent the number of changes along that branch divided by the number of sites (*65*).

## Discussion

Recent advances in strain-level genomics highlight the dynamic evolution of human gut bacteria (*6*). However, the drivers and functional consequences of microbiota within-host evolution remain poorly understood. Here, we demonstrate that *E. faecalis*, a prevalent mammalian gut commensal, rapidly acquires the ability to metabolize fructoselysine, a diet-derived nutrient, within three months in mice. Fructoselysine-adapted strains hinder enteric pathogen colonization by restricting access to essential carbohydrates. Similarly, the presence of fructoselysine-consuming *E. faecalis* in human neonates fed fructoselysine-rich formula suggests adaptation in both murine and human hosts. These findings support the concept of a microbiome self-repair mechanism, driven by evolutionary processes, that closes nutrient niches and restores colonization resistance against invading pathogens.

Resource competition drives gut bacterial evolution (*17*), with diet shaping mutation selection (*5*). Adaptive mutations in *E. coli* genes linked to carbohydrate metabolism – including galactitol (*20*), sorbitol (*22*), gluconate, N-acetylglucosamine (*40*), raffinose (*41*), and lactose (*42*) – are well-documented, with the selection also acting on the fructoselysine operon (*43*). These carbohydrates support pathogen colonization (*31*), suggesting that the concept of an evolvable niche competitor extends to other gut commensals.

We characterized multiple evolved *E. faecalis* strains from independent OMM^12^ mouse colonies and show that they exhibited increased levels of fructoselysine pathway enzymes and enhanced growth on fructoselysine. Genomic analyses revealed diverse mutations, including fructoselysine operon amplification, promoter mutations, and ICE acquisition, indicating distinct genetic mechanisms that lead to increased expression of the fructoselysine operon. *E. faecalis* that evolved in this manner appear to experience a trade-off, as fructoselysine-adapted strains show reduced growth on glucose. Similar metabolic trade-offs have been observed in *E. coli*, where raffinose/lactulose adaptation correlates with a diminished capacity to utilize allose (*41*), and similarly, gluconate adaptation impairs N-acetylglucosamine metabolism (*40*). These findings suggest that fructoselysine utilization incurs a fitness cost, with strong negative selection against its metabolism in glucose-rich environments. Supporting this notion, *E. faecalis* isolates from breast-fed infants, who have low fructoselysine exposure, do not metabolize this compound.

The prevalence of fructoselysine-degrading *E. faecalis* in formula-fed neonates aligns with our findings in the murine model and suggests that nutrient depletion is also an adaptive mechanism of the human microbiota. The neonatal microbiome, characterized by its unique metabolic and immune environment, exhibits elevated evolutionary rates (*44*). Prior studies show enhanced microbiota-mediated fructoselysine degradation in formula-fed compared to breast-fed neonates; however, whether this arises from microbial evolution or selection for preexisting fructoselysine-degrading strains remains unresolved (*36*).

Our findings experimentally support a “niche-filling model” as a driver of rapid bacterial evolution, where mutations allow species to exploit underutilized niches and preempt resources, thus blocking later colonizers (*23*). We demonstrate that *E. faecalis* adaptation to fructoselysine confers resistance to *S.* Typhimurium infection. Our findings underline the significant role of microbiota within-host evolution in microbiota-pathogen interactions and suggest that leveraging natural microbial adaptation could inform strategies aimed at enhancing microbiome-mediated pathogen resistance, with implications for infection prevention and human health.

## Declaration of generative AI and AI-assisted technologies in the writing process

ChatGPT, vs. 3.5, a language model developed by OpenAI in San Francisco, CA, U.S.A., was used for language editing and proofreading.

## Acknowledgements

The authors thank members of the Stecher Lab for excellent discussions and comments, and Prof. Curtis Huttenhower for providing expertise on metagenome analysis. D.M.S and M.Z thank the European Molecular Biology Laboratory (EMBL) for support. The authors thank Franziska Hackbarth for technical assistance and excellent maintenance of mass spectrometers at the BayBioMS, as well as Chen Meng for development of the data visualization platform omicsViewer(*64*). M.S.S. acknowledges support from the Doctoral Program “Infection Research on Human Pathogens@MvPI”.

## Funding

B.S. received funding by the European Research Council (ERC) under the European Union’s Horizon 2020 research and innovation program (grant agreement 865615), the German Research Foundation (DFG, German Research Foundation, SFB 1371, project number 395357507), and the German Center for Infection Research (DZIF). The Exploris 480 mass spectrometer was funded in part by the German Research Foundation (INST 95/1435-1 FUGG). The Infantibio-II study was initiated and financed by Töpfer GmbH (Dietmannsried, Germany). Besides manufacturing the study formula, they had no role in the conduct and management of the study or the analysis and interpretation of data or the creation of this manuscript. D.H. received funding from the German Research Foundation (DFG, Deutsche Forschungsgemeinschaft) SFB 1371 (no. 395357507). D.H. also received funding in the frame of the Joint Programming Initiative of the European Union (project name EcoBiotic) and the German Ministry of Education and Research (BMBF; FKZ 01EA2207).

## Authors contributions statement

Conceptualization: M.S.S., S.W., E.S., M.Z., D.H., B.S.

Methodology: M.S.S., S.W., D.M.S., C.L., A.C.M., M.Z., D.H., B.S.

Investigation: M.S.S., S.W., C.E., D.M.S., B.K.J., M.G., D.R., S.Wu., A.V.S., S.H., L.F., C.L.

Visualization: M.S.S., S.W., M.S.M.

Funding acquisition: D.H., B.S.

Project administration: B.S.

Supervision: T.S., A.C.M., W.D.H., M.Z., B.S.

Resources: H.O., M.H., J.L., L.O., B.N., S.B., M.B., T.S., W.D.H.

Data curation: M.S.S., S.W., D.M.S., A.D.R., P.C.M., D.G., M.S.M.

Writing – original draft: M.S.S., S.W., B.S.

Writing – review & editing: All authors

## Competing interests statement

The authors declare that no competing interests exist.

## Materials & Correspondence

Correspondence and requests for materials should be addressed to B.S.

## Supplementary materials

### Materials and Methods

#### Bacterial strains, culturing, and handling

All bacterial strains used in this study are listed in **Table S1**. OMM^12^ bacteria were routinely cultivated anaerobically (3% H_2_, 10% CO_2_, rest N_2_) at 37 °C in 10 ml pre-reduced AF liquid medium (*27*) for 24 to 48 h unless specified otherwise. *E. coli* and *S. Typhimurium* were routinely cultivated aerobically for 20 h at 37 °C and agitation in 10 ml LB liquid medium – *E. coli*: 1% (w/v) tryptone, 0.5% (w/v) yeast extract, 85.6 mM NaCl; *S.* Tm: 1% (w/v) tryptone, 0.5% (w/v) yeast extract, 300 mM NaCl. Cryo-stocks of bacterial monocultures or assembled OMM^12^ consortia were stored in the respective liquid medium with 10% (v/v) glycerol at -80 °C.

#### Preparation of cecum content medium

Cecum content medium was prepared as previously described with minor modifications (*45*). All steps were performed anaerobically. Briefly, cecum content was harvested from two 8- to 10-weeks-old gnotobiotic mice stably colonized with a modified OMM consortium lacking *E. faecalis* KB1 (OMM^11^-*E. faecalis*). Cecum content was resuspended in pre-reduced PBS at a 1:5 (w/v) ratio, incubated at 37 °C for 24 h, and pelleted via centrifugation at 10,000 x g for 3 min. The supernatant was subsequently filtered through 0.45 µm and 0.22 µm pore filters (Sarstedt, Germany) and used immediately for growth assays.

#### Quantification of bacterial growth

*E. coli* and *S. Typhimurium* growth was quantified in M9 liquid medium (400 mM Na_2_HPO_4,_ 200 mM KH_2_PO_4_, 90 mM NaCl, 37 mM NH_4_Cl, 1 mM MgSO_4_, 100 µM CaCl_2_, 37.6 µM thiamine, 3.2 mM histidine, 0.07 µM vitamin B12, 0.58 µM p-aminobenzoic acid, 0.08 µM D(+)-biotin, 1.62 µM nicotinic acid, 0.2 µM calcium pantothenate, 1.45 µM pyridoxine hydrochloride, and 0.54 µM thiamine-HCl x 2 H_2_O) supplemented with 5 g/L glucose or fructoselysine (see section “Synthesis and isolation of N-ε-(1-deoxy-D-fructosyl)-L-lysine (fructoselysine)”). Overnight cultures were washed 3 times with PBS and resuspended in M9 liquid medium at an OD_600_ of 0.04. Cultures (100 µl per well) were incubated aerobically at 37 °C in 96-well round bottom plates (Thermo Fisher Scientific, U.S.A.), and OD_600_ was recorded over a period of 12 to 24 h using a microplate reader (BioTek, U.S.A.).

*E. faecalis* growth was quantified in modified M9 liquid medium (MM9; 40 mM Na_2_HPO_4_, 20 mM KH_2_PO_4_, 9 mM NaCl, 37 mM NH_4_Cl, 2 mM MgSO_4_, 100 µM CaCl_2_, 2x (v/v) MEM vitamin solution, 1x (v/v) MEM amino acid solution, 2x (v/v) MEM non-essential amino acid solution) supplemented with 5 g/L glucose or fructoselysine, and in cecum content medium. Overnight cultures were washed 3 times with PBS and resuspended in MM9 or cecum content medium at an OD_600_ of 0.04 or 0.01, respectively. Cultures were incubated anaerobically at 37 °C in a 96-well round bottom plate using 100 µl or 200 µl per well for growth in MM9 or cecum content medium, respectively. OD_600_ was recorded over a period of 48 to 72 h, and the area under the curve was calculated using GraphPad Prism 10.1.2 (GraphPad Software, U.S.A.).

#### Synthesis and isolation of N-ε-(1-deoxy-D-fructosyl)-L-lysine (fructoselysine)

Fructoselysine was synthesized as described previously (*46*). 2.5 mmol boc-lysine and 15 mmol glucose were heated in 105 ml methanol for 4 h under reflux. Following cooling, methanol was evaporated *in vacuo* and the residue was taken up in 150 ml water. 5 ml glacial acetic acid was added, and the solution was applied to a 2.5 cm x 20 cm column filled with the strongly acidic cation exchange resin LEWATIT MonoPlus S 108 (Kurt Obermeier GmbH, Germany) equilibrated with 250 mL 6 M HCl and 250 ml water. Residual sugar and uncharged by-products were removed with 300 ml double-distilled water and the crude product was eluted with 300 ml pyridinium pH 6.0 acetate buffer. The solution was evaporated to dryness, the residue was taken up in 30 ml 0.1 N pH 3.0 pyridinium formate buffer and the pH of the solution was adjusted to 3.0 with formic acid. The solution was applied to a 1.5 cm x 50 cm column filled with the strongly acidic cation exchange resin DOWEX 50 WX-8 equilibrated with 250 ml 6 M HCl, 250 ml water, 250 ml 2 M pyridine, 250 ml water, and 250 ml of 0.1 N pH 3.0 pyridinium formate buffer. The column was rinsed with 10 ml 0.1 N pH 3.0 pyridinium formate buffer, and fructoselysine was eluted with 600 ml 0.4 N pH 4.35 pyridinium formate buffer. The eluate was collected in 6 ml fractions via fraction collector (Bio-Rad Laboratories, Germany). Amino compounds were detected in 1 µl spots in each fraction on a thin layer chromatography plate by spraying with a mixture of ethanol, acetic acid, and ninhydrin at a 100/3/0.1 ratio (w/v/v) or with 1% (w/v) triphenyl tetrazolium chloride in 1 M NaOH, followed by heating at 80 °C in a drying chamber. Fructoselysine eluted in the 100-160 ml eluate fraction. Fractions with a positive reaction in both elution buffers were pooled and repeatedly evaporated to dryness under reduced pressure after addition of water. The residue was taken up in 2 ml methanol and precipitated in 25 ml ice-cold butanone. The precipitate was filtered and stored in a desiccator overnight. The white residue was then collected and stored at -18 °C.

#### Assembly of OMM^12^ communities

To prepare OMM^12^ consortia for inoculation of mice, individual bacteria were cultivated in monoculture anaerobically for 24 h at 37 °C in 10 mL AF medium. Monocultures of ancestral strains or evolved isolates were mixed at equal ratio based on the OD_600_ of cultures and stored at -80 °C with 10% (v/v) glycerol as specified. All strains are listed in the **Table S1**.

#### Animal breeding and maintenance

Germ-free and OMM^12^- colonized C57BL/6J mice were bred and maintained in flexible film isolators (North Kent Plastic Limited, U.K.) at germ-free mouse facilities at the Max-von-Pettenkofer Institute (Facility MUC; Munich, Germany). Stable gnotobiotic OMM^12^ mouse colonies were also bred at Hannover Medical School (Facility HAN; Hannover, Germany), Helmholtz Center for Infection Research (Facility BS; Braunschweig, Germany), or ETH Zurich (Facility ZUC; Zurich, Switzerland), under controlled environmental conditions and strict 12-hour or 10/14-hour (Facility HAN) light/dark cycles. At all animal facilities mice were bred according to local animal welfare law in agreement with EU-directive 2010/63. HZI Brunswick: license #32.5/325.1.53/56.1. MHH: mouse breeding and experiments were approved by the local institutional Animal Care and Research Advisory Committee and permitted by the Lower Saxony State Office for Consumer Protection and Food Safety - LAVES (reference numbers: 42500/1H, and 2023/242). ETH Zurich: All animal experiments were reviewed and approved by Tierversuchskommission, Kantonales Veterinäramt Zürich under license ZH109/2022, complying with the cantonal and Swiss legislation. Mice received autoclaved double-distilled water and sterilized pelleted mouse diet ad libitum. Details on diets and sterilization methods used in the different facilities are listed in **Table S4**. Sentinel animal tests were performed routinely to monitor and verify the colonization status of germ-free and gnotobiotic animals. Mice were euthanized by cervical dislocation.

#### Mouse experiments

All animal experiments were reviewed and approved by the local authorities (Regierung von Oberbayern, under the ethical approval number ROB-55.2-2532.Vet_02-20-118). Experiments were performed at the Max von Pettenkofer Institute (MUC), other facilities maintained stably colonized OMM^12^ colonies and sacrificed mice for sampling. Germ-free C57BL/6J female and male 6- to 20-week-old mice were randomly assigned to experimental groups. Experiments were performed aseptically using the IsoCage P system (Tecniplast, Germany) or Han-gnotocages (ZOONLAB, Germany), allowing 2 to 5 mice per cage. Mice were colonized with bacterial OMM^12^ consortia or cecum content from OMM^12^ - colonized mice from MUC or HAN via oral and rectal inoculation as described previously (*20*). Cecum content for FMT experiments was collected as described in section “Re-isolation of evolved OMM^12^ bacteria from mouse cecum content”. Colonization with *E. coli* and infection with *S. Typhimurium* strains was performed as described previously (*20*).

#### Genomic DNA extraction

Genomic bacterial DNA was extracted from weighed mouse intestinal content and bacterial pellets as described previously (*24*). Briefly, genomic DNA was isolated using a phenol-chloroform-based extraction method and column-purified using the NucleoSpin gDNA clean-up kit (Macherey-Nagel, Germany) according to the manufacturer’s instructions. For whole genome sequencing, extracted DNA was RNA-depleted via RNaseA (Thermo Fisher Scientific, U.S.A.) treatment according to the manufacturer’s instructions between extraction and column-purification. Extracted DNA was routinely stored at -20 °C.

Genomic DNA of bacterial isolates from infant stool was isolated from bacterial overnight cultures using the Qiagen Genomic DNA Buffer Set and Genomic Tip 100/G (Qiagen, Germany) according to the manufacturer’s instructions.

#### Analysis of microbial community composition

The absolute abundance of OMM^12^ bacteria in feces from OMM^12^- colonized gnotobiotic mice was determined via 16S rRNA gene quantitative PCR, using species-specific 16S rRNA primers and hydrolysis probes as described previously (*24*). Briefly, species-specific 16S rRNA genes were amplified using FastStart Essential DNA Probes Master (Roche, Switzerland) and a LightCycler 96 system (Roche) according to the manufacturer’s instructions. Sample signals were normalized to the signal measured with linearized plasmids, containing species-specific 16S rRNA gene sequences to achieve absolute quantification of 16S rRNA gene copy numbers.

#### Proteome analysis

Proteomes of evolved *E. faecalis* isolates and the ancestor strain KB1 were compared using mass spectrometry analysis. To this end, *E. faecalis* KB1 and evolved isolates were inoculated into liquid APF medium (18.5 g/L BHI glucose-free, 15 g/L trypticase soy broth glucose-free, 5 g/L yeast extract, 14.35mM K_2_HPO_4_, 1 mg/L hemin, 2.5 g/L sugar mix (1:1 arabinose, fucose, lyxose, rhamnose, xylose), 2 g/L inulin, 2 g/L xylan, 0.025 g/L mucin, 0.5 mg/L menadione, 3% heat-inactivated fetal calf serum, 2.85mM cysteine-HCl, 3.77 mM Na_2_CO_3_) at a starting OD_600_ of 0.04, and cultivated anaerobically at 37 °C until the required growth phase was reached(*27*). *E. faecalis* KB1 and SW160 were assessed in exponential (T1), early stationary (T2), and late stationary (T3) growth phases, and *E. faecalis* KB1, SW160, SW178, SW269, SW293, and SW328 were assessed in late stationary growth phase in two separate experiments. Cultures were pelleted via centrifugation at 13,000 x g and 4 °C for 5 min, washed twice with PBS, pelleted again, and pellets were stored at -80 °C.

Bacteria were lysed using the SPEED protocol(*47*) with slight adaptations(*48*). Shortly, 100 µl of 100% TFA was added to the cell pellet, heated for 5 min at 55 °C, and neutralized with 900 µl 2 M Tris. Protein concentrations were determined using Bradford reagent (Biorad Laboratories, U.S.A.) and samples were stored at -80 °C. Three biological replicates were prepared for all samples. 50 µg total protein amount per sample was reduced and alkylated (9 mM tris(2-carboxyethyl)phosphine (TCEP) and 40 mM chloroacetamide (CAA) for 5 min at 95 °C). Samples were diluted with deionized water to a final concentration of 1 M Tris and 5% TFA. Trypsin (Roche, Switzerland) was added in a trypsin:protein ratio of 1:50. Samples were incubated overnight at 37 °C and 400 rpm, acidified and desalted with self-packed desalting tips (Empore C18 3M; CDS Analytical LLC, U.S.A).

Proteomic measurements were performed on a Vanquish^TM^ Neo UHPLC (microflow configuration; Thermo Fisher Scientific, U.S.A.) coupled to an Orbitrap Exploris^TM^ 480 mass spectrometer (Thermo Fisher Scientific, U.S.A.). Peptides were applied onto a commercially available Acclaim PepMap 100 C18 column (2 μm particle size, 1 mm ID × 150 mm, 100 Å pore size; Thermo Fisher Scientific, U.S.A.) and separated on a stepped gradient from 3% to 31% solvent B (0.1% FA, 3% DMSO in ACN) in solvent A (0.1% FA, 3% DMSO in HPLC grade water) over 60 min. A flow rate of 50 μl/min was applied. The mass spectrometer was operated in DDA and positive ionization mode. MS1 full scans (360 – 1300 m/z) were acquired with a resolution of 60,000, a normalized automatic gain control target value of 100%, and a maximum injection time of 50 ms. Peptide precursor selection for fragmentation was carried out using a cycle time of 1.2 s. Only precursors with charge states from two to six were selected, and dynamic exclusion of 30 s was enabled. Peptide fragmentation was performed using higher energy collision-induced dissociation and a normalized collision energy of 28%. The precursor isolation window width of the quadrupole was set to 1.1 m/z. MS2 spectra were acquired with a resolution of 15,000, a fixed first mass of 100 m/z, a normalized automatic gain control target value of 100%, and a maximum injection time of 40 ms.

For proteomic analysis, MaxQuant v1.6.3.4 with its built-in search engine Andromeda(*49*),(*50*) was used. MS2 spectra were searched against a protein database generated for the evolved *E. faecalis* strain SW160 supplemented with common contaminants (built-in option in MaxQuant). The protein annotation is available at https://github.com/MonicaSteffi/OMM12. Trypsin/P was specified as the proteolytic enzyme. Precursor tolerance was set to 4.5 ppm, and fragment ion tolerance to 20 ppm. The minimal peptide length was defined as seven amino acids. The “match-between-run” function was disabled. Carbamidomethylated cysteine was set as a fixed modification, and methionine oxidation and N-terminal protein acetylation were defined as variable modifications. Results were adjusted to a 1% false discovery rate on peptide spectrum match (PSM) level and protein level, employing a target-decoy approach using reversed protein sequences. Label-Free Quantification (LFQ)(*49*) was used for protein quantification with at least 2 peptides per protein identified. Only proteins identified in at least two out of three biological replicates were considered. Missing values were imputed as half the lowest detected LFQ intensity per protein, unless the imputed value exceeded the sample’s 20^th^ percentile, in which case the 20^th^ percentile value was used.

#### Isolation of evolved OMM^12^ isolates from mouse cecum content

Bacteria were isolated from the cecum content of gnotobiotic mice that had been colonized with the OMM^12^ consortium for at least one year at the time of isolation. Mice were sacrificed and isolation performed under anaerobic conditions. Cecum content was harvested and filtered through a 70 µm cell strainer (Corning, U.S.A.) before adding 10 to 20 ml PBS containing 10% glycerol. Content was kept at -80 °C for long-term storage. To re-isolate bacteria, a 10-fold dilution series of cecum content was prepared in PBS, and dilutions were spread on AF agar plates, which were incubated at 37 °C for 48 to 72 h. Colonies were resuspended in 200 µl AF medium and re-streaked on AF agar plates which were incubated at 37 °C for 24 h. Single colonies were cultured in AF medium for 24 h at 37 °C and stored at a final glycerol concentration of 10% at -80 °C. *Bacteroides caecimuris*, *Enterocloster clostridioformis*, and *Blautia coccoides* were identified based on colony morphology, *E. faecalis* and *Clostridium innocuum* were isolated as described previously (*51*). *Muribaculum intestinale* was isolated using magnetic-activated cell sorting as described previously (*52*). Anti-*M. intestinale* antibodies were produced by vaccinating mice with acetic acid-inactivated *M. intestinale* culture. Bacterial isolates were identified via Gram-staining and Sanger sequencing (Eurofins Genomics Germany GmbH, Germany) of 16S rRNA amplicons (**Table S2**).

#### Quantification of mutation rate polymorphisms in evolved *E. faecalis* isolates

The mutation rates of *E. faecalis* KB1 and evolved *E. faecalis* isolates were assessed using the Luria-Delbrück fluctuation assay (*53*). Briefly, a single bacterial colony was dissolved in PBS, the OD_600_ was adjusted to 0.1, and the solution was diluted 1:100 with Brain-heart infusion (BHI) (Thermo Scientific™ Oxoid™, Germany) liquid medium. 250 µl were transferred to a 96-well round bottom plate and incubated for 20 h at 37 °C. A 10-fold dilution series of the culture was prepared and 100 µl of each dilution were spread on BHI agar plates with or without 50 µg/ml rifampicin, respectively, and plates were incubated at 37 °C for 20 h. Biological triplicate measurements were performed for each *E. faecalis* isolate. Mutation rates were calculated using the webSalvador 0.1 tool (*54*) with the Lea-Coulson method where estimated Nt is the mean number of colonies on unselective agar plates, mutant count is the number of colonies on selective medium, both per 250 µl, and ε is the culture proportion spread on agar plates (ε = 0.4).

#### Whole genome sequencing of bacterial isolates

Genomic DNA of evolved bacterial isolates from OMM^12^ - colonized gnotobiotic mice was obtained as described in “Genomic DNA extraction”. Whole-genome sequencing was performed by Eurofins Genomics Germany GmbH using standard genomic library preparation, including unique dual indexing, and Illumina (U.S.A.) paired-end short-read sequencing technology with a read length of 2 x 250 bp and a sequencing depth of five million read pairs per sample.

Genomic DNA of bacterial isolates from infant stool was sequenced by Novogene Europe (U.K.) using PacBio Sequel II sequencing technology in continuous long-read mode. The average read length was between 16-23k for all samples. Genome *de novo* assembly was performed using Canu v2.2 (*55*) and assemblies were subsequently circularized using Circlator v1.5.5 (*56*).

#### Single nucleotide polymorphism and structural variation calling

Genome sequencing reads of evolved bacterial isolates were analyzed using the Geneious Prime software package version 2025.0.2 (Biomatter Ltd., New Zealand) as described before (*57*). Reads were imported as paired reads and mapped to the reference genome of the respective unevolved ancestor strains (GenBank accession numbers: CP065312.1, CP065314.1, CP065316.1, CP065317.1, CP065319.1, CP065320.1) using the built-in Geneious mapper with medium sensitivity and up to five iterations. Single nucleotide polymorphisms were defined as reference genome deviations with minimum total coverage depth of 100, minimum variant frequency of 80%, maximum variant *P*-value of 10^-6^, and minimum strand-bias *P*-value of 10^-5^ when exceeding 65% bias. Copy number variations based on amplification or deletion of genomic content were identified using mean read coverage analysis, dividing the mean coverage of a region by the mean coverage of the remaining genome (*57*). Horizontal gene transfer events were detected by successively mapping sequencing reads unable to map to the species reference genome to the reference genomes of other OMM^12^ strains.

#### Phylogenetic clustering of bacterial isolates

Phylogenetic trees based on single nucleotide polymorphisms of bacterial isolates from OMM^12^ - colonized gnotobiotic mice were created using the Geneious Prime software package version 2025.0.2. A multiple alignment of bacterial isolate genomes was performed and sites with identical bases were masked. Phylogenetic trees were created with the in-built Randomized Axelerated Maximum Likelihood (RAxML, version 8.2.11) method, using the GTR GAMMA nucleotide model and the rapid hill-climbing algorithm. The number of starting trees or bootstrap replicates and the parsimony random seed were set to 1, respectively. Phylogenetic trees based on genomes of *E. faecalis* infant stool isolates and reference genomes (GCF_000392875.1; CP065317.1) were created with kSNP4.1, using the maximum likelihood method and a k-mer size of 17, determined through Kchooser4 to optimize single nucleotide polymorphism detection accuracy. Phylogenetic trees were visualized using iTOL version 7(*58*).

#### Generation of *E. faecalis* mutants

Fructoselysine operon deglycase genes (GenBank locus ID: I5Q89_01965 and 5Q89_01960) were deleted in evolved *E. faecalis* SW160 via homologous recombination using the plasmid pLT06 (*59*). All primers and plasmids are listed in **Table S2**. Briefly, homologous regions upstream and downstream of deglycase genes were amplified from *E. faecalis* SW160 genomic DNA via PCR and assembled with an erythromycin resistance gene(*60*) and *Eco*RI- and *Bam*HI-restricted plasmid pLT06 using the NEBuilder HiFi DNA Assembly Master Mix. Enzymes and assembly reagents were obtained from New England Biolabs (U.S.A.) and used according to the manufacturer’s instructions. 10-beta electrocompetent *E. coli* were (New England Biolabs) transformed with the assembled plasmid at 37 °C and plasmid DNA was isolated using the QIAprep spin mini kit (Qiagen, Germany), and plasmid insert sequences were confirmed via Sanger sequencing (Eurofins Genomics Germany GmbH, Germany).

*E. faecalis* SW160 were treated with lysozyme (Carl Roth, Germany) and mutanolysin (Merk, Germany) to generate electrocompetent cells. Briefly, a bacterial culture with an OD_600_ of 0.5 to 1.0 was incubated on ice for 20 min and bacteria were pelleted via centrifugation at 4,000 x g at 4 °C for 20 min. The pellet was washed with pre-chilled 10% (v/v) glycerol, divided in two tubes, resuspended in 500 µl lysozyme solution (10 mM Tris pH 8.0, 20 g/L sucrose, 10 mM EDTA, 50 mM NaCl, 25 µl/ml lysozyme, 2 µg/ml mutanolysin), and incubated at 37 °C for 20 min. Bacteria were pelleted via centrifugation at 13,000 x g at 4 °C for 1 min, washed three times with pre-chilled electroporation buffer (171.15 g/L sucrose, 10% (v/v) glycerol), and resuspended in 250 µl electroporation buffer.

65 µl electrocompetent *E. faecalis* SW160 were transformed with the newly generated plasmid (pLT06 derived) via electroporation (1.8 kV, time constant: 4.0 to 4.7 ms), recovered in BHI liquid medium supplemented with 30 g/L sucrose for 2 h at 37 °C, and spread on BHI agar plates supplemented with 20 µg/ml chloramphenicol and 0.04 g/L X-Gal (company). Plates were subsequently incubated for 48 h at 30 °C. Blue colonies were inoculated in BHI liquid medium supplemented with 20 µg/ml chloramphenicol and incubated overnight at 44 °C. A culture 10-fold dilution series was prepared, plated on BHI agar plates supplemented with 20 µg/ml chloramphenicol and 0.04 g/L X-Gal, and plates were incubated overnight at 44 °C. Blue colonies were inoculated into BHI liquid medium and incubated at 30 °C. The culture was propagated until white colonies, which were chloramphenicol-sensitive and erythromycin-resistant, appeared after plating on BHI agar supplemented with 0.04 g/L X-Gal. The plates were then incubated overnight at 44 °C. Deletion of fructoselysine degylases was confirmed via PCR.

#### *S. Typhimurium* fructoselysine operon deletion

A *S. Typhimurium*^avir^ mutant strain with a deletion of the fructoselysine operon (*gfrABCDEF::cat*) was generated by phage P22-mediated transduction. The *S.* Typhimurium Z5537 donor strain was constructed using the λ Red recombination system (*61*). The entire *gfr* operon was replaced by homologous recombination with the chloramphenicol resistance cassette from plasmid pKD3. Strain *S. Typhimurium* Δ*gfrABCDEF* was created by phage P22 transduction of the *gfrABCDEF*::*cat* allele into the recipient strain *S. Typhimurium* M2702 (*62*).

#### *E. faecalis* isolation from infant stool

*E. faecalis* was isolated from stool samples of formula- or breast-fed infants of the Infantibio study. Stool samples were collected within a previous study registered at the German Clinical Trials Registry (‘‘Deutsches Register Klinischer Studien’’) with the trial number DRKS00012313, and sample details were published previously (*39*). Briefly, samples were frozen at -80 °C in glycerol no later than 24 h after collection. *E. faecalis* was isolated by selective plating on CHROMagar^TM^ Orientation plates (CHROMagar, France). Stool was spread on CHROMagar^TM^ Orientation plates and incubated overnight at 37 °C. Turquoise blue colonies were re-streaked on *Enterococcus* Differential Agar (2.5 g/L yeast extract, 5 g/L tryptone from casein, 1 g/L glucose, 330 mg/L thallium(I) acetate, 10 ml/l 2,3,5-Triphenyl-tetrazoliumchlorid), and pink colonies grown after overnight incubation at 37 °C were used to inoculate BHI liquid medium. Cultures were incubated overnight at 37 °C and stored at -80 °C in 10% (v/v) glycerol. Isolates were identified via sequencing of the 16S rRNA gene amplicons (Eurofins Genomics Germany GmbH, Germany) using primers fD1, fD2 and rP1 (**Table S2**)

#### Untargeted and targeted metabolome analysis

50 mg of mouse cecum content was mixed with 250 µl ultra-pure DNase- and RNase-free water (Invitrogen AG, U.S.A.) and pelleted via centrifugation at 10,000 g for 4 min at 4 °C. 200 µl supernatant (=cecal water) were frozen on dry ice and stored at -80 °C. Metabolites were extracted by adding 500 µl water and bead-beating supernatants with 200 µl 0.1 mm-diameter zirconia beads, followed by centrifugation at 15,000 x g at 4 °C for 10 min. Supernatant was transferred into a 96-well microtiter plate using 20 µl per well and mixed with 50 µl acetonitrile, 50 µl methanol, and an internal standard mixture. Samples were incubated at -20 °C for 1 h and precipitated proteins were pelleted via centrifugation at 4,500 x g and 4 °C for 15 min. 50 µl supernatant was transferred to a 96-well microtiter plate and mixed with 50 µl water, followed by centrifugation at 4,500 x g and 4 °C for 15 min. Sulfamethoxazole, caffeine, ipriflavone, and warfarin were used as internal standards at a final concentration of 80 nM each. Liquid chromatography-mass spec analysis was performed as previously described(*63*). Chromatographic separation was performed by normal-phased chromatography using InfinityLab Poroshell 120 HILIC-Z, 2.1 x 150 mm, 2.7-micron pore size columns (Agilent Technologies, U.S.A.), a 1200 Infinity UHPLC system (Agilent), and mobile phases A (water, 5 mM ammonium formate, 0.1% (v/v) formic acid) and B (acetonitrile, 5 mM ammonium formate, 0.1% (v/v) formic acid). The column compartment was kept at 25 °C. 5 µl sample were injected at 98% B and 0.250 ml/min flow, followed by 98% B until minute 3, a gradient to 70% B until minute 11, a gradient to 60% B until minute 12, a gradient to 5% B to minute 16, and remaining at 5% B until minute 18 before re-equilibration to 98% until minute 20. The qTOF (Agilent) was operated in positive scanning mode (50-1500 *m/z*) with the following source parameters: VCap = 3000 V; nozzle voltage = 0 V; gas temp = 225 °C; drying gas flow = 11 L/min; nebulizer = 40 psig; sheath gas temp = 225 °C; sheath gas flow = 10 L/min; fragmentor = 300 V; and Octopole RF Vpp = 450 V. Online mass calibration was performed using a second ionization source and a constant flow of reference solution (121.0508 and 922.0097 *m/z*). Tandem mass spectrometry analysis (LC-MS/MS) was performed for fructoselysine using the chromatographic separation and source parameters described above and the targeted-MS/MS mode of the instrument with a preferred inclusion list for parent ion with 20 ppm tolerance, Iso width set to ‘narrow width’ and collision energy set to either 10, 20, or 40 eV. The MassHunter Qualitative Analysis Software version 10.2 (Agilent) and Mass Profiler Professional version 15.1 (Agilent) were used for untargeted feature extraction and peak alignment, respectively. Software was used with standard parameters allowing tolerances for mass of 0.002 amu or 20 ppm and for a retention time of 0.3 min or 2%. The resulting table contains a list of features (i.e. *m/z* and RT) and their relative intensity per sample. Before statistical analysis, these features were submitted for blank removal (removal of features from blank injections), imputation (replacement of missing and zero values), normalization by tissue weight, quantile normalization, scaling, and centering. The final processed table was submitted to Principal Component Analysis (PCA) using the prcomp function in R 4.4.1 and RStudio version 2023.06.1. The MassHunter Quantitative Analysis Software version 7.0 (Agilent) was used for retention time-based peak integration and accurate mass measurement of fructoselysine for semi-targeted feature extraction.

#### Lipocalin-2 quantification

Lipocalin-2 was quantified in mouse feces via enzyme-linked immunosorbent assay as described previously (*20*). Briefly, 96-well plates (Brand, Germany) were coated at 4 °C overnight with 50 µl/well anti-lipocalin-2 capture antibody (R&D Systems, U.S.A.) diluted 1:200 in PBS. Excess capture antibodies were removed by washing three times with 0.05% (v/v) Tween-20 in PBS and the plate was blocked for 1 h using 100 µl/well 2% (w/v) BSA in PBS. The plate was washed three times and 50 µl of sample were added in duplicates followed by 1 h of incubation. The plate was washed six times and 50 µl/well anti-lipocalin-2 detection antibody (R&D Systems) diluted 1:200 in 2% (w/v) BSA PBS were added followed by 1 h of incubation. Excess detection antibodies were removed by washing 6 times and 100 µl/well HRP-streptavidin diluted 1:1,000 in PBS were added followed by 1 h of incubation and six additional wash steps. 100 µl substrate (1 mg ABTS in 10 ml 0.1 M NaH_2_PO_4_ pH 4, 5 µl H_2_O_2_) were added and the plate was incubated for 30 min followed by measuring absorbance at λ_405_. Lipocalin-2 concentrations were calculated using a lipocalin-2 standard curve included on each plate.

#### Metagenomics analysis

Metagenomic reads were sequenced using the Illumina NovaSeq 6000 platform with the Shotgun NovaSeq pyrosequencing method. Library preparation was performed using the NEBNext Ultra II FS DNA Kit, and sequencing was conducted on a NovaSeq S1 flow cell with 150n paired-end (PE) reads, yielding approximately 6.8 Mbases of data.

Raw paired-end sequencing reads underwent quality control and adapter removal using fastp. High-quality reads were then aligned to the reference genome with BWA-MEM, utilizing the -x intractg parameter optimized for bacterial genomes to enhance sensitivity for intracellular organisms. PCR and optical duplicates were subsequently identified and removed using Picard MarkDuplicates.

#### Variant calling in metagenomes

Genomic variants, including single nucleotide polymorphisms and insertions/deletions (indels), were identified through a custom Snakemake pipeline centered on the LoFreq variant caller. The aligned reads were preprocessed with LoFreq-specific steps, including error correction via the Viterbi algorithm and indel quality score recalibration using the DINDEL model (--dindel flag). Variants were called in parallel mode with stringent filtering parameters: a significance threshold of 1E-4, minimum coverage of 25x, minimum base quality of 25, minimum alternative base quality of 25, and minimum mapping quality of 60. This methodology excels at detecting low-frequency variants in bacterial populations, enabling sensitive and specific identification of single nucleotide polymorphisms and indels that arise during experimental evolution. To analyze the functional distribution of variants, we mapped single nucleotide polymorphisms to Clusters of Orthologous Groups (COGs) using genome annotations from GFF files. COG functional categories were obtained from the COG-20 database definitions. For visualization, we plotted allele frequencies (AF) across all samples, with variants grouped by their corresponding COG functional categories. Only variants with AF > 0.5 in at least one sample were included.

#### Coverage Analysis

Sequencing coverage data were generated from aligned reads and converted into BigWig format using bamCoverage, which normalizes and condenses alignment data into a coverage representation at 50-bp resolution. This facilitated efficient storage and visualization of genome-wide coverage patterns. Coverage data were processed in R using the rtracklayer package for BigWig file imports and the dplyr package for data manipulation. A specific region of interest in *Enterococcus faecalis* KB1 (coordinates 407,337–412,093), flanked by 5,000 bp on either side, was selected for detailed analysis. Coverage values were extracted from .bam.bw files at 50-bp intervals, annotated with their midpoint positions, and smoothed using a 3-point rolling mean (via the zoo package) to reduce noise while preserving signal fidelity. Coverage data were visualized using ggplot2, with samples grouped by timepoint (days 20, 40, 80, and offspring) and experimental condition (with or without *E. coli*). For samples without *E. coli*, mean coverage inside and outside the region of interest was computed, and the fold change at the final timepoint was determined to assess enrichment. This approach identified genomic regions with significant copy number variations or structural changes across experimental conditions and timepoints of *E. faecalis* KB1 colonization.

#### Statistical analysis

Statistical analyses were performed with GraphPad Prism (v. 10.4.1) or R (v. 4.4.1). Continuous data from non-normally distributed independent samples were analyzed using exact Mann-Whitney *U* tests or Kruskal-Wallis tests followed by Dunn’s post-hoc tests. Continuous data from normally distributed independent samples were analyzed using ANOVA with Dunnet’s multiple comparison test. Bivariate correlations were analyzed using Spearman’s rank correlation. For microbial community analyses, Bray-Curtis distances were calculated to assess the difference in microbial population among samples using Principal Coordinates Analysis. PERMANOVA tests were conducted on Bray-Curtis distances and adjusted using Bonferroni multiple testing correction. For untargeted metabolome analyses, differences in metabolite feature intensity were assessed using ANOVA, and *P*-values were FDR-corrected for multiple hypotheses testing using the Benjamini-Hochberg procedure. Proteomics-based differential protein expressions were analyzed using Student t-tests and adjusted using the Benjamini-Hochberg algorithm. To identify fructoselysine utilization operon amplifications, differences in coverage between the region of interest and the surrounding genomic context were compared using a two-sample Wilcoxon signed-rank test. This non-parametric test compares coverage values inside versus outside the region, producing one *P*-value per sample. *P*-values were adjusted for multiple comparisons using the Benjamini-Hochberg method to control the false discovery rate (FDR). The fold-change in average coverage (inside/outside ratio) was calculated to quantify enrichment. Samples were ranked by FDR-adjusted *P*-values and fold-changes to highlight those with the most significant and pronounced coverage differences. Sample sizes and *P*-values are indicated in figures or figure legends. *P*-values below 0.05 were considered significant.

#### Figure schemes

Figure and experimental schematics in Fig. 1A; 2A-B; 3A; 4E; 4A; and Fig. S1C; S2A; S3D; S4A; S4C; S6A; S7A were created using BioRender.com. Fructoselysine degradation scheme was created using ChemSketch version 2024.

#### Data availability

All metagenome sequencing data generated in this study were deposited in the NCBI Sequence Read Archive repository and is accessible via the BioProject accession number PRJNA1196824. All raw genome data will also be deposited in the NCBI Sequence Read Archive repository. Proteomic raw data and MaxQuant output files have been deposited to the ProteomeXchange Consortium via the PRIDE partner repository and can be accessed using the identifier PXD062369 (https://proteomecentral.proteomexchange.org/cgi/GetDataset?ID=PXD062369, reviewer account username, password: NPK91a6b0iZe). Metabolomic data generated in this study were deposited in the MetaboLights repository (https://www.ebi.ac.uk/metabolights/and) and accessible via the accession number MTBLS10327. Source data are provided with this paper.

---

**Fig. S1:**
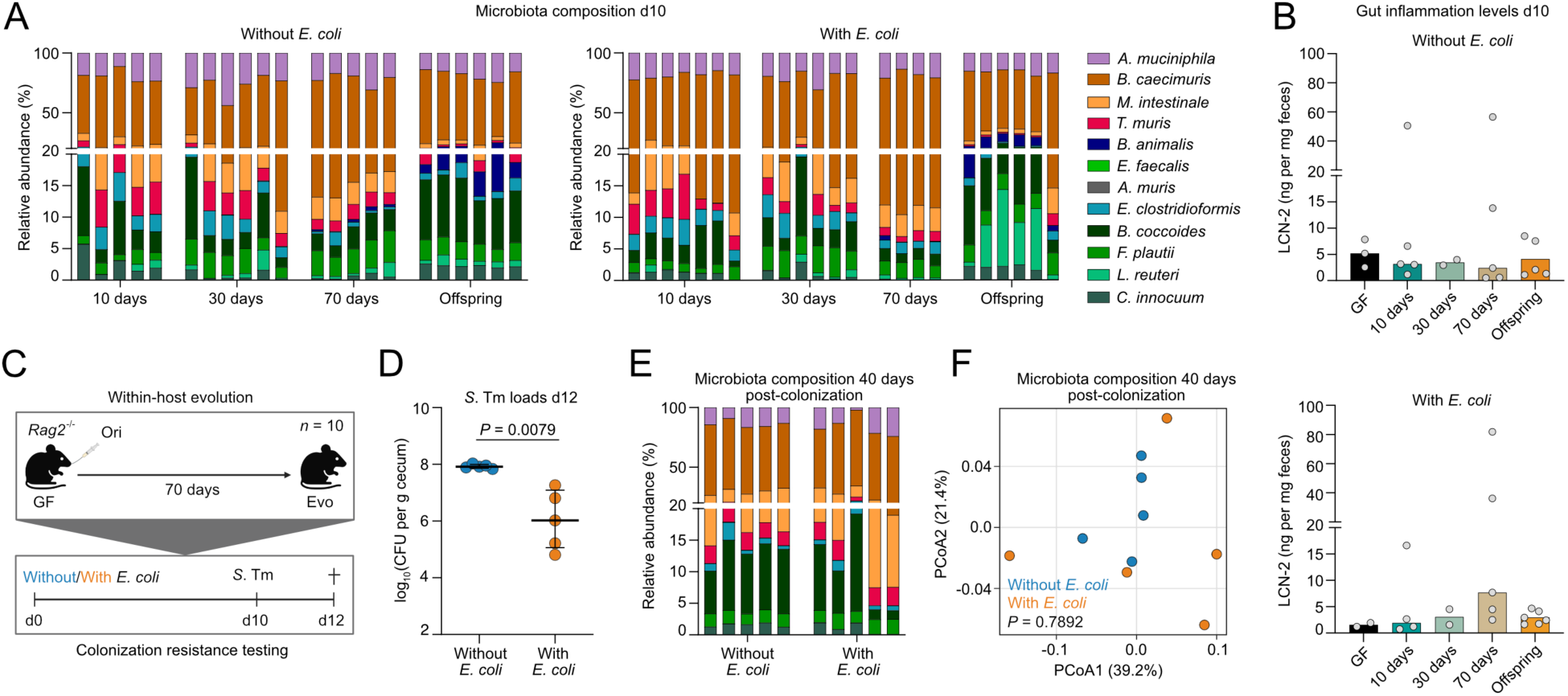
*E. coli*-mediated colonization resistance in long-term colonized OMM^12^ mice is independent of gut inflammation and adaptive immunity. **(A)** Fecal microbiota composition in mice shown in Fig. 1B on the day of *S*. Typhimurium (*S*. Tm) infection (d10). Relative abundances were determined via qPCR, quantifying species-specific 16S rRNA gene copies. Bars correspond to individual mice. **(B)** Feces lipocalin-2 concentrations in mice shown in Fig. 1B on the day of *S*. Tm infection (d10). Germ-free mice serve as controls. **(C)** Experimental scheme for assessing colonization resistance against *S*. Tm in mice lacking mature T- and B-cells. Germ-free *Rag2^-/-^* mice were colonized with the original OMM^12^ community(*24*) for 70 days. Subsequently, mice were inoculated with either *E. coli* (*n* = 5) or PBS (*n* = 5). Colonization resistance was assessed as described in Fig. 1B. **(D)** Cecum *S*. Tm loads on day two post infection (d12). Data represent medians with Interquartile ranges, and statistical analysis was performed using a Mann-Whitney *U* test. The detection limit is 10 CFU per g. (**E** to **F)** Fecal microbiota composition 40 days after the colonization of *Rag2^-/-^*mice with the OMM^12^ community. **(E)** Relative abundances were determined via qPCR, quantifying species-specific 16S rRNA gene copies. Bars correspond to individual mice. **(F)** Principal coordinate analysis was based on the Bray-Curtis dissimilarity distance matrix of relative OMM^12^ abundance profiles. Statistical analysis was performed using a PERMANOVA with Bonferroni correction, comparing mice colonized with or without *E. coli*.

**Fig. S2:**
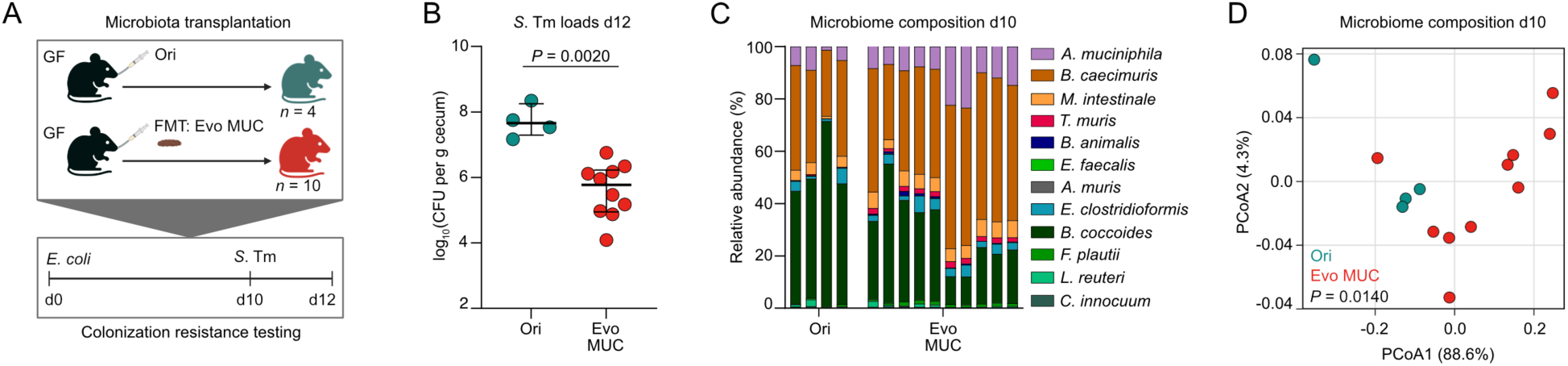
Transplantation of an independently evolved OMM^12^ community provides colonization resistance against *S. Typhimurium*. **(A)** Assessment of colonization resistance against *S*. Typhimurium (*S*. Tm) in germ-free mice colonized with the original OMM^12^ community or with an evolved OMM^12^ community (Evo MUC) harvested from stably colonized mice (Evo MUC; equals “offspring” mice shown in Fig. 1). Original OMM^12^ was freshly assembled from single-species cryo-stocks. Colonization resistance was assessed as described in Fig. 2B. **(B)** Cecum *S*. Tm loads on day two post-infection (d12). Data represent medians with interquartile ranges, and statistical analysis was performed using a Mann-Whitney *U* test. The detection limit is 10 CFU per g. **(C** and **D)** Fecal microbiota composition on the day of *S*. Tm infection (d10). **(C)** Relative abundances were determined via qPCR, quantifying species-specific 16S rRNA gene copies. Bars correspond to individual mice. **(D)** Principal coordinate analysis was based on the Bray-Curtis dissimilarity distance matrix of relative OMM^12^ abundance profiles. Statistical analysis was performed using a PERMANOVA with Bonferroni correction.

**Fig. S3:**
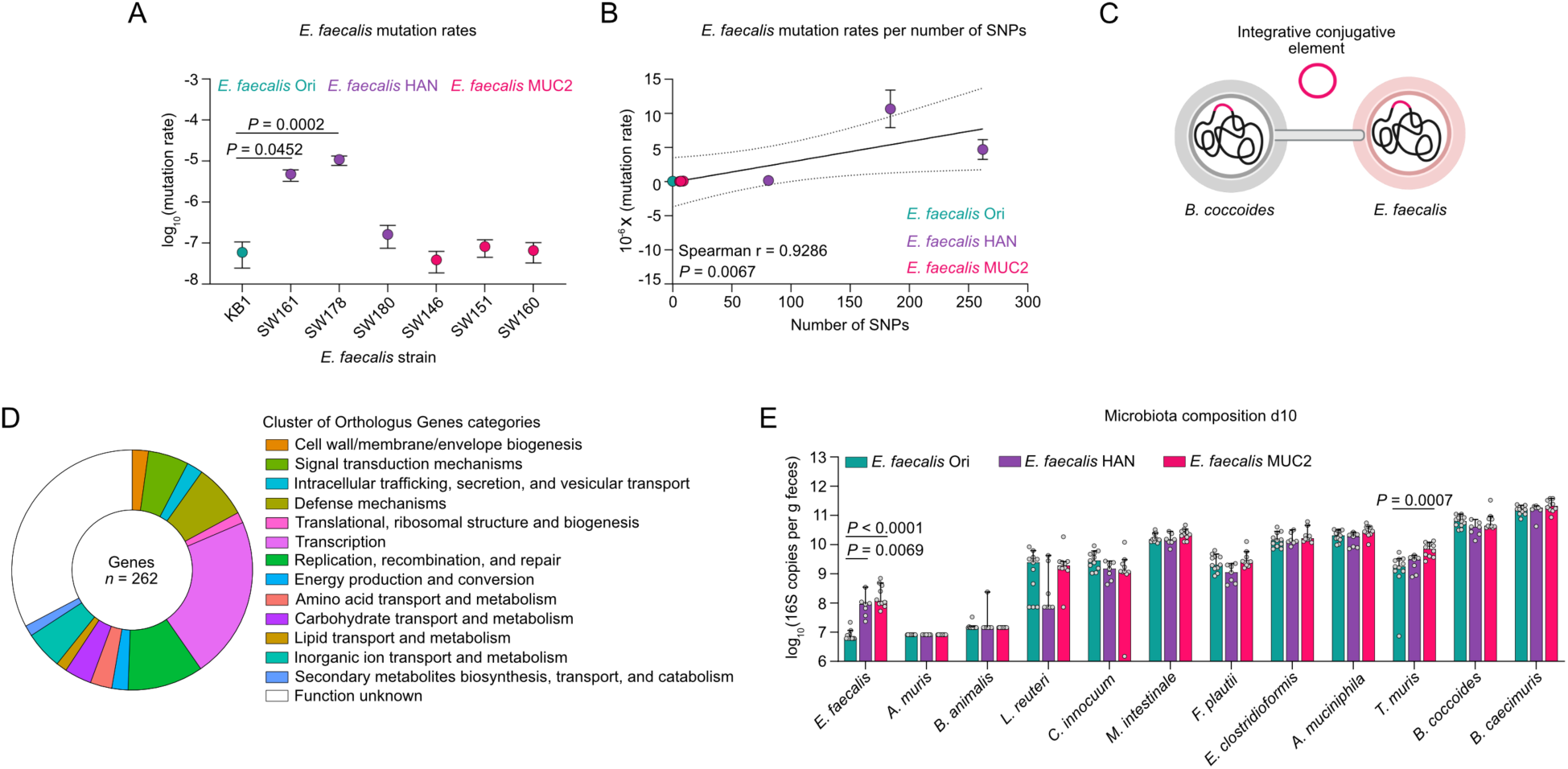
Characterization of evolved *E. faecalis* isolates from OMM^12^ mice housed in different animal facilities. **(A)** Mutation rates of *E. faecalis* KB1 and select evolved isolates from two OMM^12^ mice housed in different animal facilities (*E. faecalis* HAN, *E. faecalis* MUC2) were estimated using the Luria-Delbrück fluctuation assay, which quantifies the spontaneous development of rifampicin resistance. The isolates represent the phylogenetic clusters depicted in Fig. 3b. Statistical analysis was performed using a Dunnett’s multiple comparison test, comparing each evolved isolate to *E. faecalis* KB1. Only *P*-values below 0.05 shown. **(B)** Correlation between mutation rates and the number of single nucleotide polymorphisms identified in each *E. faecalis* strain. Spearman’s rank correlation and corresponding *P*-values are displayed, and error bars represent 95% confidence intervals. **(C)** Integrative conjugative element scheme. Several evolved *E. faecalis* isolates carry a 156 kb integrative conjugative element that originates from *B. coccoides* and is horizontally transferred during within-host evolution. **(D)** Proportions of genes per Clusters of Orthologous Genes (COG) category encoded on the integrative conjugative element. **(E)** Fecal microbiota composition on the day of *S.* Typhimurium (*S*. Tm) infection (d10) in mice shown in Fig. 3 c, d, e. Absolute abundances were determined via qPCR, quantifying species-specific 16S rRNA gene copies per g feces. Data represent medians with 95% confidence intervals, and statistical analyses were performed using Dunn’s post-hoc tests, comparing mice colonized with the original *E. faecalis* (*E. faecalis* Ori) to those colonized with evolved *E. faecalis* isolates (*E. faecalis* HAN, *E. faecalis* MUC2). Only *P*-values below 0.05 shown.

**Fig. S4:**
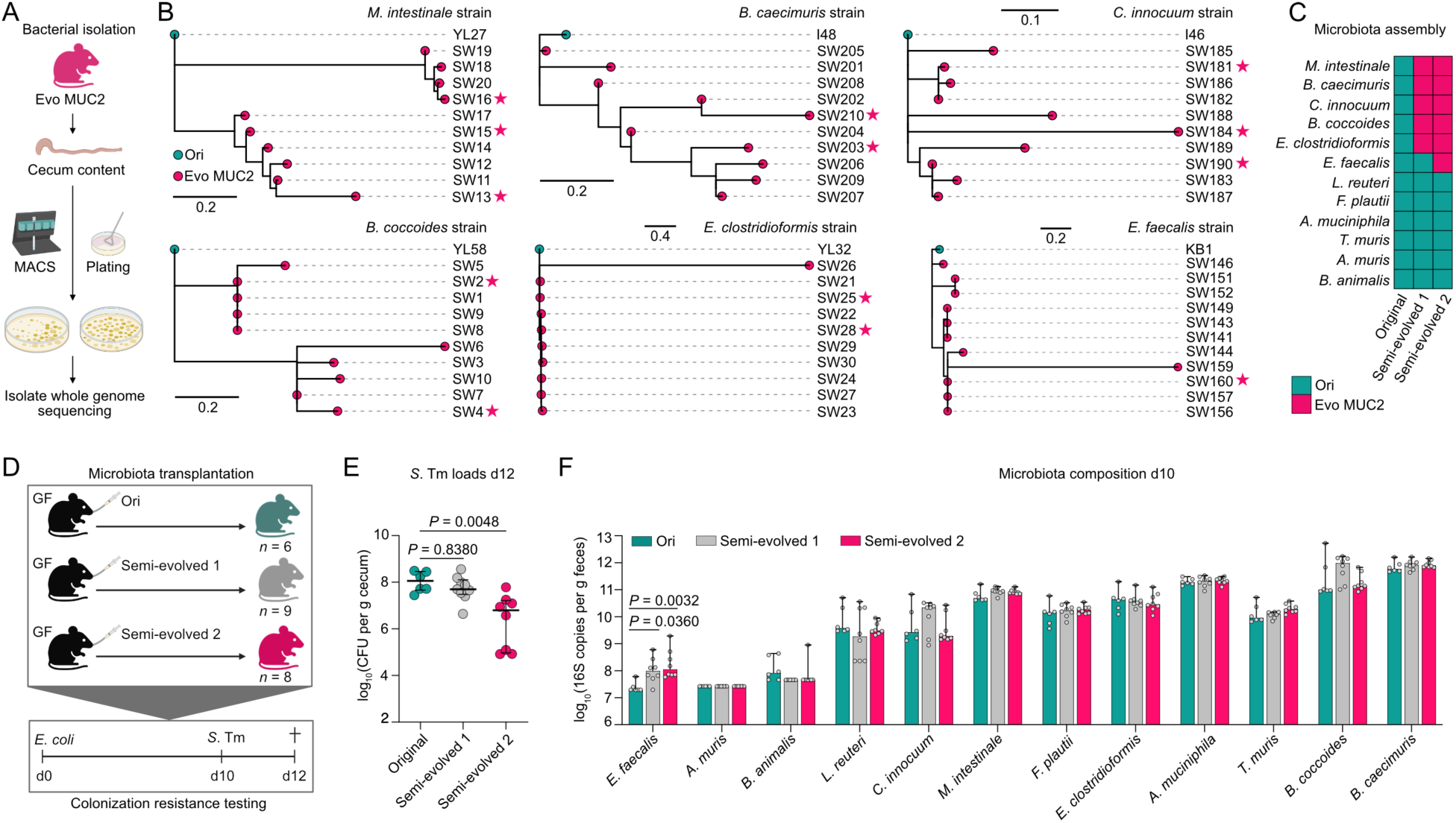
Evolved isolates of OMM^12^ species other than *E. faecalis* fail to confer colonization resistance against *S. Typhimurium*. **(A)** Bacterial isolation scheme. Evolved isolates of six different OMM^12^ species were obtained from the cecum content of a single long-term colonized OMM^12^ mouse (Evo MUC2). Isolates were subjected to short-read whole genome sequencing. *E. faecalis* isolates are also shown in Fig. 3b. MACS, magnetic-activated cell sorting. **(B)** Single nucleotide polymorphism-based phylogenetic trees of evolved isolates (*n* = 10-11 per species) and original strains. A polymorphism allele frequency cut-off of 0.8 was applied. The tree was created using the Random Accelerated Maximum Likelihood method, with branch lengths representing the substitutions per variable site. Stars highlight strains used in (D). **(C)** Semi-evolved microbiota assembly scheme. The original OMM^12^ community(*26*) was freshly assembled from single-species cryo-stocks, while semi-evolved communities were assembled using evolved isolates from several species (magenta) along with original strains of other species (turquoise). **(D)** Assessment of colonization resistance against *S*. Typhimurium (*S*. Tm) in germ-free mice colonized with original or semi-evolved OMM^12^ communities. Colonization resistance was assessed as described in Fig. 3B. **(E)** Cecum *S*. Tm loads on day two post-infection (d12). Data represent medians with interquartile ranges, and statistical analysis was performed using a Dunn’s post-hoc test. The detection limit is 10 CFU per g. **(F)** Fecal microbiota composition on the day of *S.* Tm infection (d10). Absolute abundances were determined via qPCR, quantifying species-specific 16S rRNA gene copies per g feces. Data represent medians with 95% confidence intervals, and statistical analyses were performed using Dunn’s post-hoc tests, comparing the original with each semi-evolved OMM^12^ community. Only *P*-values below 0.05 are shown.

**Fig. S5:**
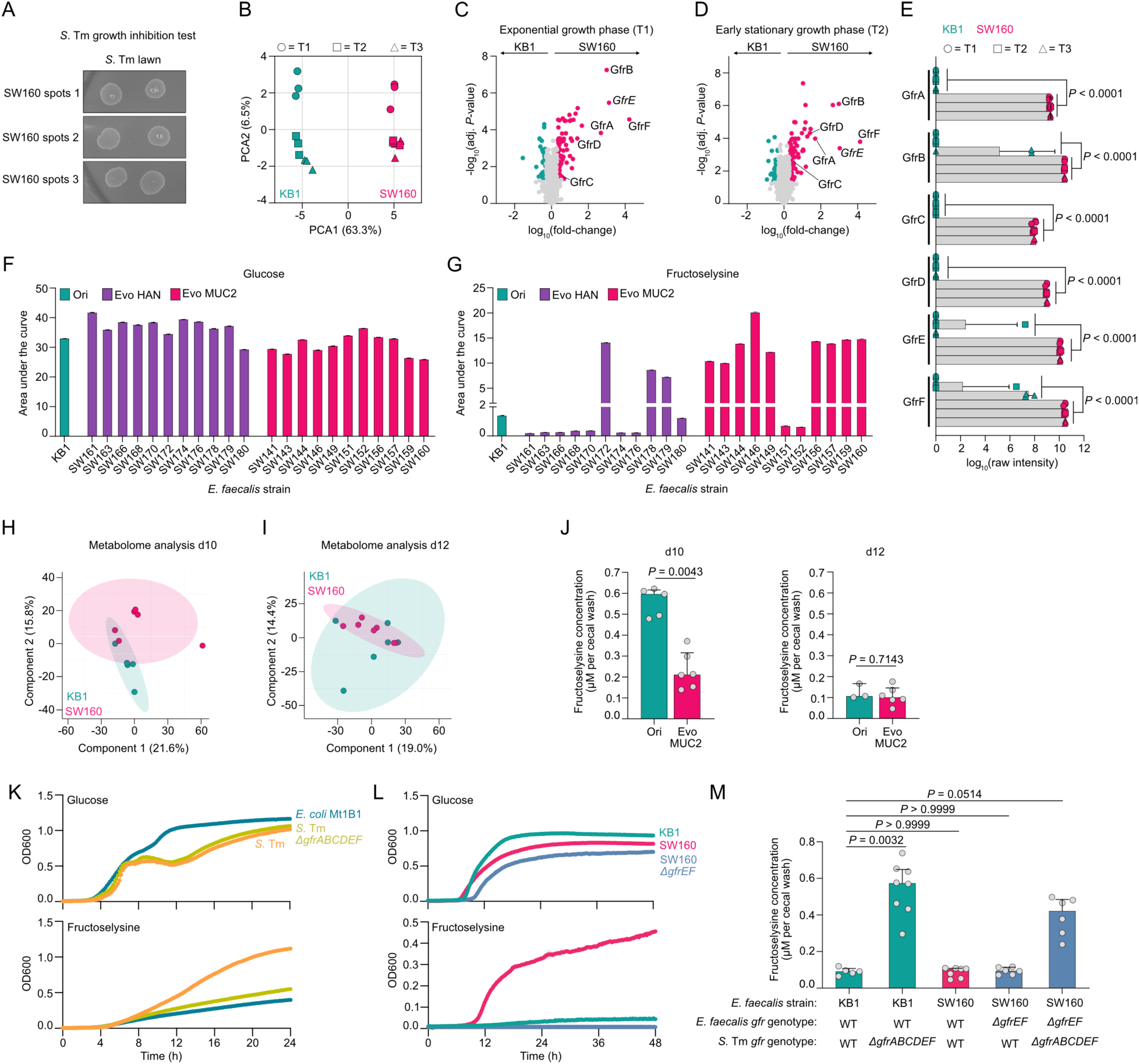
*S. Typhimurium* and evolved *E. faecalis* compete for fructoselysine in the gut. **(A)** Growth inhibition assay. *E. faecalis* SW160 was spotted on a lawn of *S*. Typhimurium (*S*. Tm). **B**-**E)** Data from the experiment described in Fig. 4E. *E. faecalis* strains were grown in AF liquid medium, and proteomes were analyzed during the exponential (T1), early stationary (T2), or late stationary (T3) growth phases. **(B)** Principal component analysis of proteomes. **(C** to **D)** Volcano plots showing proteins overrepresented in *E. faecalis* KB1 (turquoise) or SW160 (magenta). Significance cut-offs were log_10_(2) and -log_10_(0.05) for fold-changes and Benjamini-Hochberg adjusted t-test *P*-values, respectively. Fructoselysine operon-encoded proteins are highlighted. **(E)** Raw mass spectrometric intensities of fructoselysine operon-encoded proteins. Bars represent means with standard deviation, and statistical analyses were performed using Mann-Whitney *U* tests. **(F** and **G)** Area under the curve based on the growth of *E. faecalis* strains in minimal medium with glucose **(F)** or fructoselysine **(G)** as the sole carbon source. Data represent means with 95% confidence intervals. **(H** and **I)** Same experiment as shown in Fig. 4e. Metabolomes of cecal washes were analyzed in mice colonized with original *E. faecalis* KB1 or evolved *E. faecalis* SW160 on the day of *S*. Tm infection (d10; KB1: *n* = 5; SW160: *n* = 6; **H**) or on day two post-infection (d12; KB1: *n* = 6; SW160: *n* = 7; **I**). Principal coordinates analysis was performed on Bray-Curtis distance matrices. **(J)** Same experimental setup as in Fig. 4e, but comparing the metabolomes of mice colonized with an original OMM^12^ community(*24*) or an evolved community (Evo MUC2). Fructoselysine levels in cecal washes on the day of *S*. Tm infection (d10; Ori: *n* = 5; Evo MUC2: *n* = 6) and two days post-infection (d12; Ori: *n* = 6; Evo MUC2: *n* = 6). Data represent medians with Interquartile ranges, and statistical analyses were performed using Mann-Whitney *U* tests. **(K** and **L)** Growth curves of *S*. Tm, *S*. Tm *ΔgfrABCDEF*, and *E. coli* **(K)** or of *E. faecalis* KB1, *E. faecalis* SW160, and *E. faecalis* SW160 *ΔgfrEF* **(L)** in minimal medium with glucose or fructoselysine as the sole carbon source. **(M)** Same experiment as shown in Fig. 4I. Fructoselysine levels in cecal washes on day two post-infection (d12). Data represent medians with interquartile ranges, and statistical analysis was performed using a Dunn’s post-hoc test.

**Fig. S6:**
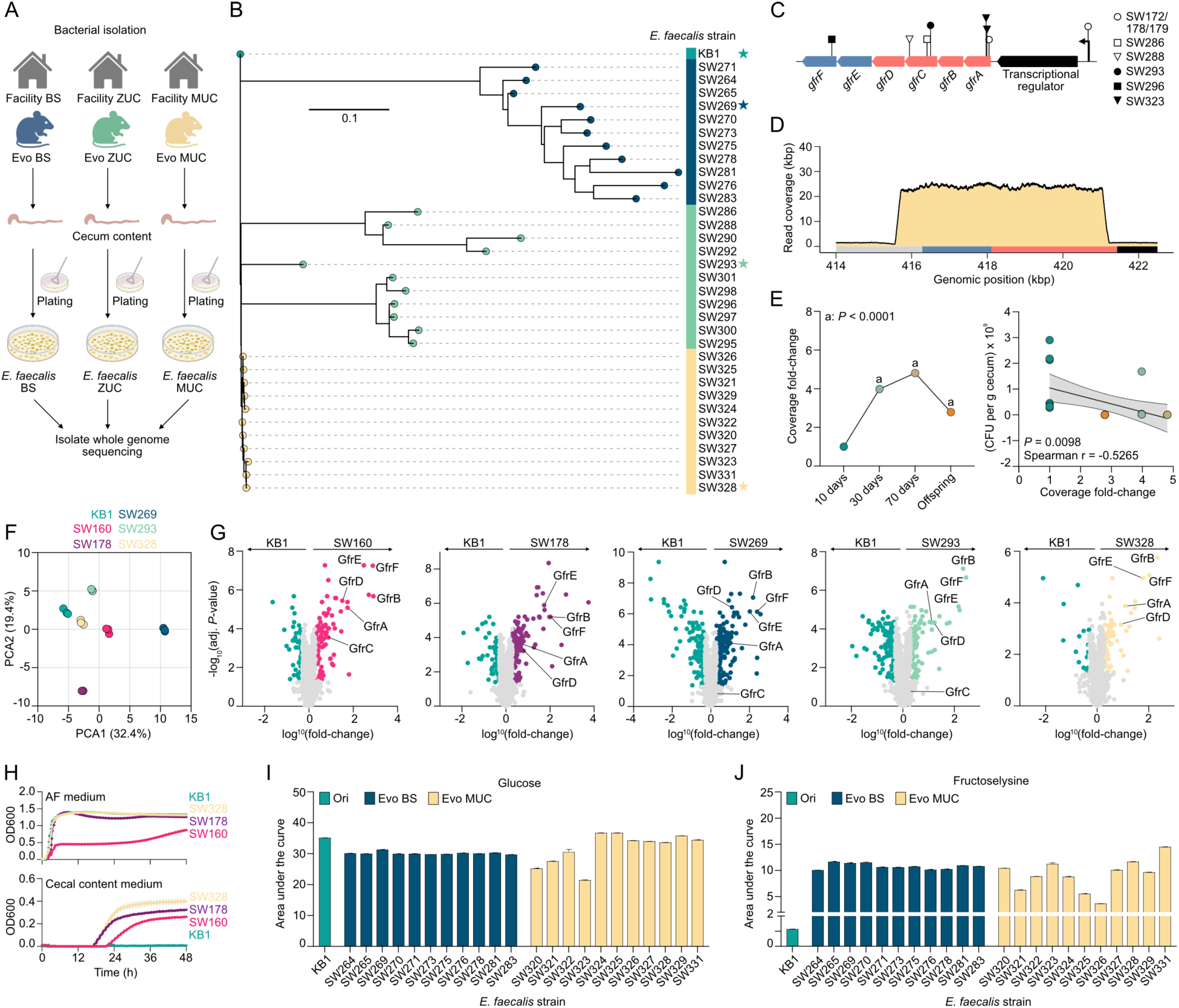
Different within-host evolutionary trajectories converge to enable *E. faecalis* fructoselysine utilization. **(A)** *E. faecalis* isolation scheme. Evolved *E. faecalis* were isolated from the cecum contents of long-term colonized OMM^12^ mice (Evo BS, Evo ZUC, Evo MUC) from three different animal facilities. The Evo MUC mouse is shown in Fig. 1 as “offspring” and in Fig. S2. Isolates were subjected to short-read whole genome sequencing (*n* = 1 mouse per facility). **(B)** Single nucleotide polymorphism-based phylogenetic tree of evolved *E. faecalis* isolates (*n* = 11 per mouse) and original *E. faecalis* KB1. A polymorphism allele frequency cut-off of 0.8 was applied. The tree was created using the Random Accelerated Maximum Likelihood method, and branch lengths represent substitutions per variable site (*n* = 2,703). Stars highlight strains used in (F) and (G). **(C)** Fructoselysine operon single nucleotide polymorphisms in *E. faecalis* isolates from this study. Phosphotransferase, deglycase, and transcriptional regulator genes are indicated in red, blue, and black, respectively. **(D)** Genome coverage plot of *E. faecalis* SW328. Sequencing reads were mapped against the *E. faecalis* KB1 reference genome. Colours correspond to the fructoselysine operon scheme in (C). The downstream region of the operon is indicated in grey. **(E)** Correlation of the fructoselysine operon amplification and colonization resistance against *S*. Tm in mice shown in Fig. 1. Coverage fold-changes of the *E. faecalis* fructoselysine operon relative to a 10 kb region outside of the operon were assessed in mouse metagenomes (left). Coverage fold-changes were correlated with *S*. Typhimurium loads in all mice from the respective groups shown in Fig. 1C (right). Metagenomes of one (10 days, 70 days, Offspring) or two mice (30 days) were analyzed per group. All mice were colonized with *E. coli*. For the 30 days timepoint, coverage fold-changes from two mice were pooled and the mean is shown. Statistical analyses were performed using Wilcoxon signed-rank tests (left) and Spearman’s rank correlation with corresponding *P*-values (right). Error bars represent 95% confidence intervals. **(F** and **G)** Proteomes of evolved *E. faecalis* isolates from five independent long-term colonized OMM^12^ mouse colonies. *E. faecalis* strains were grown in AF liquid medium, and proteomes were analyzed during the late stationary growth phase. **(F)** Principal component analysis of proteomes; **(G)** Volcano plots showing proteins overrepresented in *E. faecalis* SW160 (magenta), SW178 (purple), SW269 (blue), SW293 (green), and SW328 (beige), compared to their ancestor *E. faecalis* KB1 (turquoise). Significance cut-offs were log_10_(2) and -log_10_(0.05) for fold-changes and Benjamini-Hochberg adjusted t-test *P*-values, respectively. Fructoselysine operon-encoded proteins are highlighted. **(H)** Growth curves of *E. faecalis* KB1, SW160, SW178, and SW328 in AF or cecal content liquid medium. Cecal content originated from a mouse long-term colonized with the OMM^11^-*E. faecalis* community. (**I** and **J)** Area under the curve based on the growth of *E. faecalis* strains in minimal medium with glucose **(I)** or fructoselysine **(J)** as the sole carbon source. Data represent means with 95% confidence intervals.

**Fig. S7:**
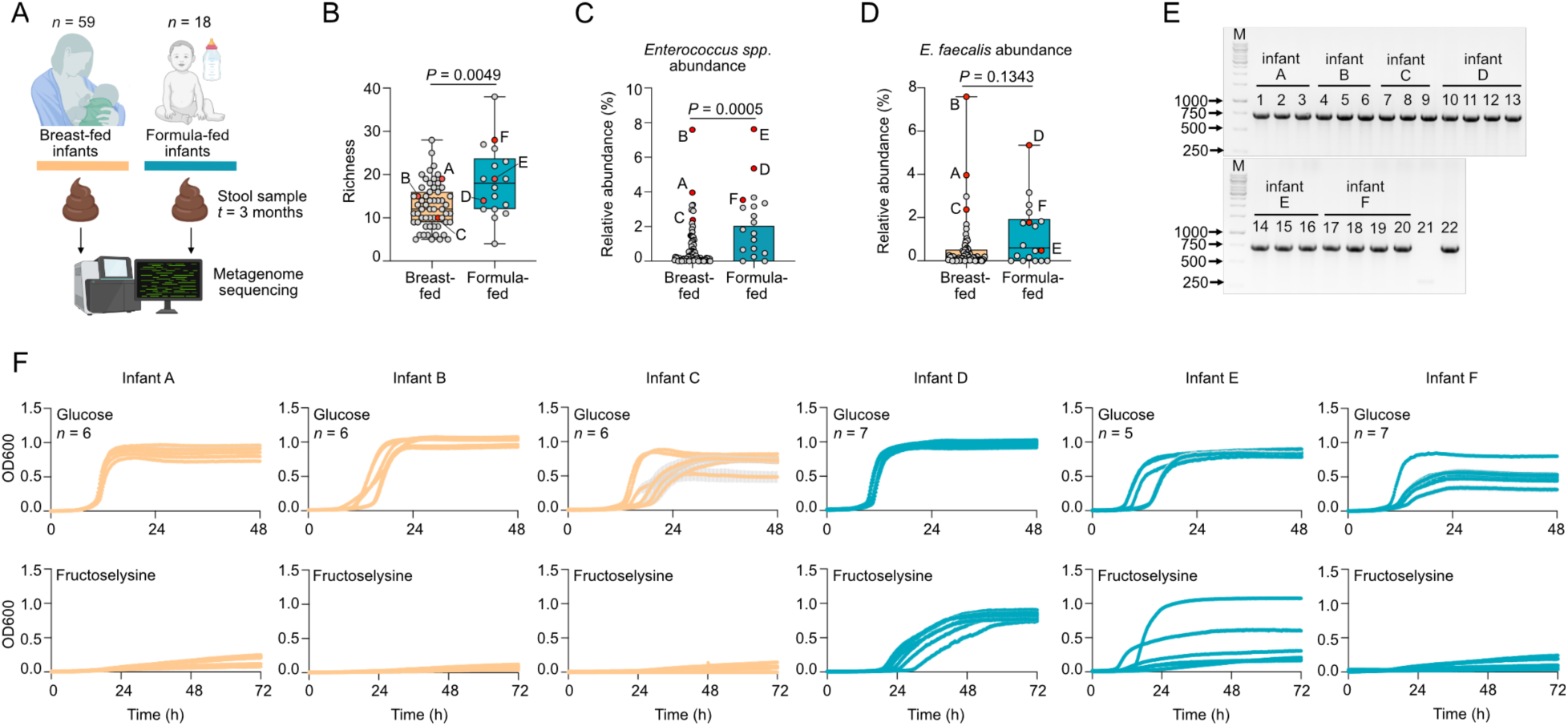
Characterization of *E. faecalis* isolates from breast- and formula-fed infants. **(A)** Study scheme. Stool samples from three-month-old breast- or formula-fed infants were collected and subjected to metagenome sequencing. **(B)** Gut microbiota richness among all breast- and formula-fed infants included in the study. Infants used for *E. faecalis* isolation are highlighted (breast-fed: A, B, C; formula-fed: D, E, F). **(C** and **D)** Metagenome-based relative abundance of *Enterococcus* species (C) and *E. faecalis* (D) across all breast- and formula-fed infants included in the study. Statistical analyses were performed using Mann-Whitney *U* tests. **E)** PCR-generated *gfrEF* gene amplicons in *E. faecalis* isolates from breast- (A, B, C) and formula-fed (D, E, F) infants (*n* = 3-4 isolates per infant). Numbers correspond to different samples, and results are representative of all obtained *E. faecalis* isolates. Genomic DNA from *E. faecalis* SW160 11*gfrEF* and *E. faecalis* SW160 was used as templates for PCRs 21 and 22, respectively. The expected amplicon size is 746 bp. M, DNA marker. **(F)** Growth curves of all *E. faecalis* isolates per infant in minimal medium with glucose or fructoselysine as the sole carbon source. Corresponding areas under the curve are shown in Fig. 5B.

**Table S1:**
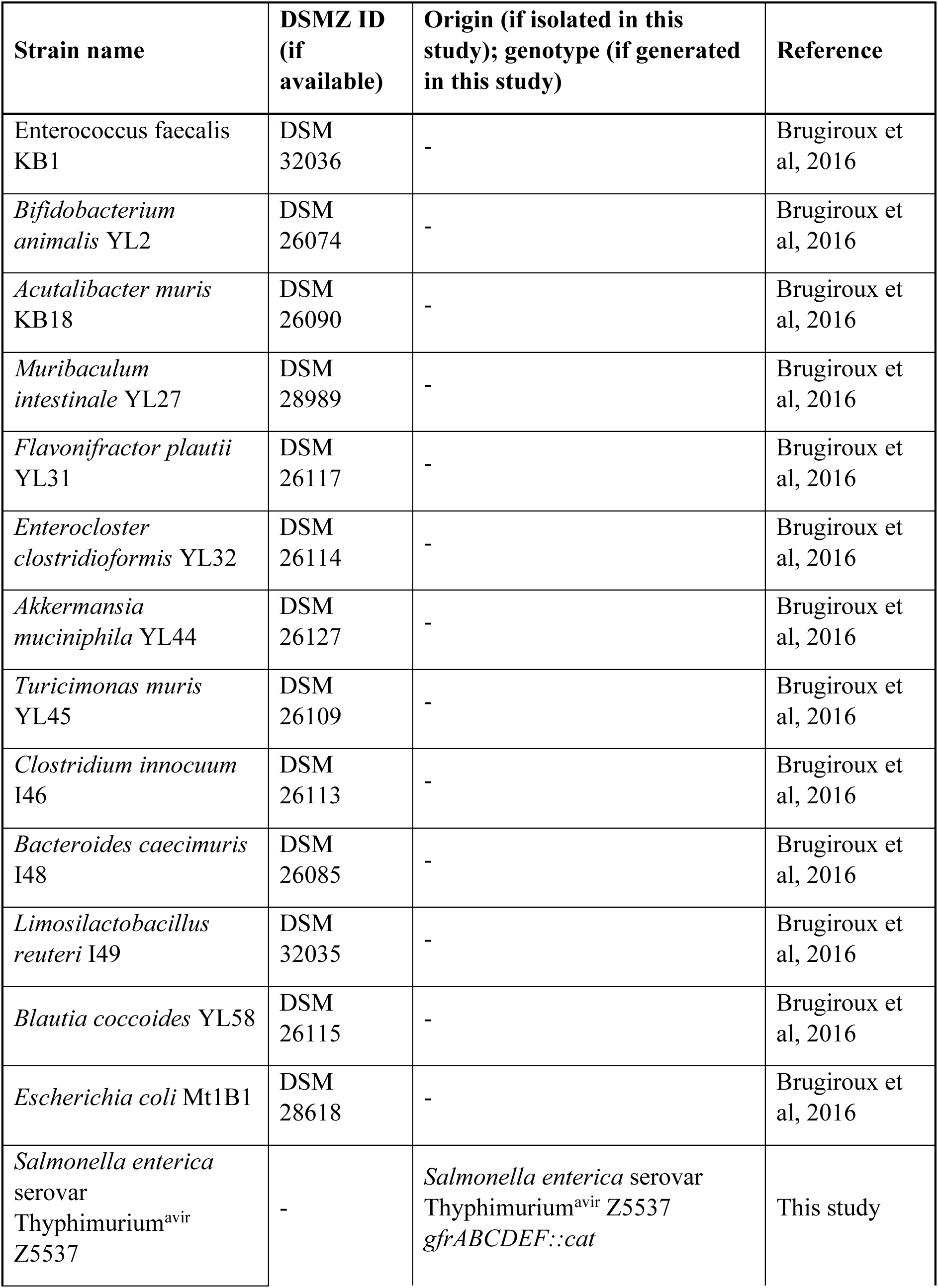
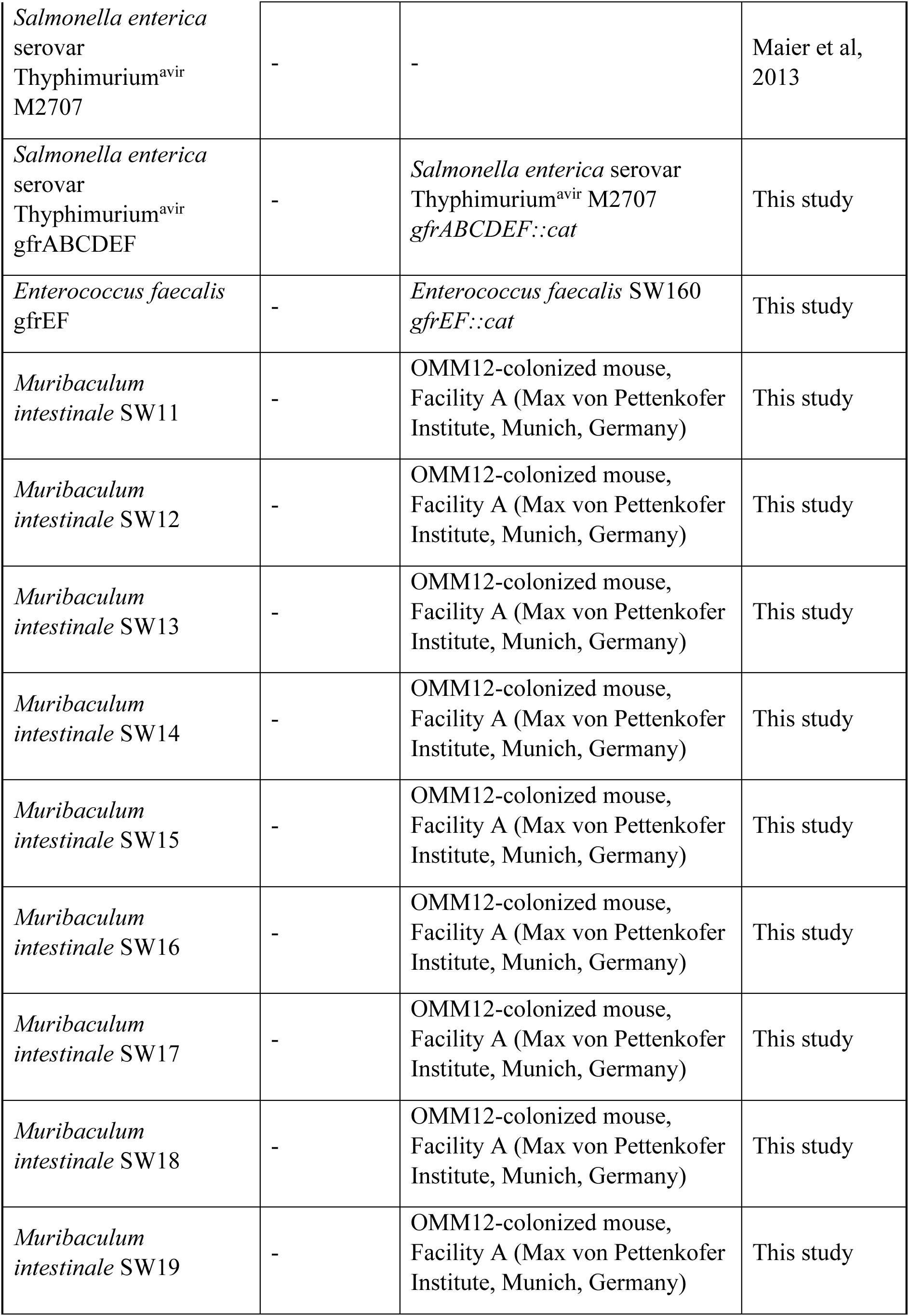
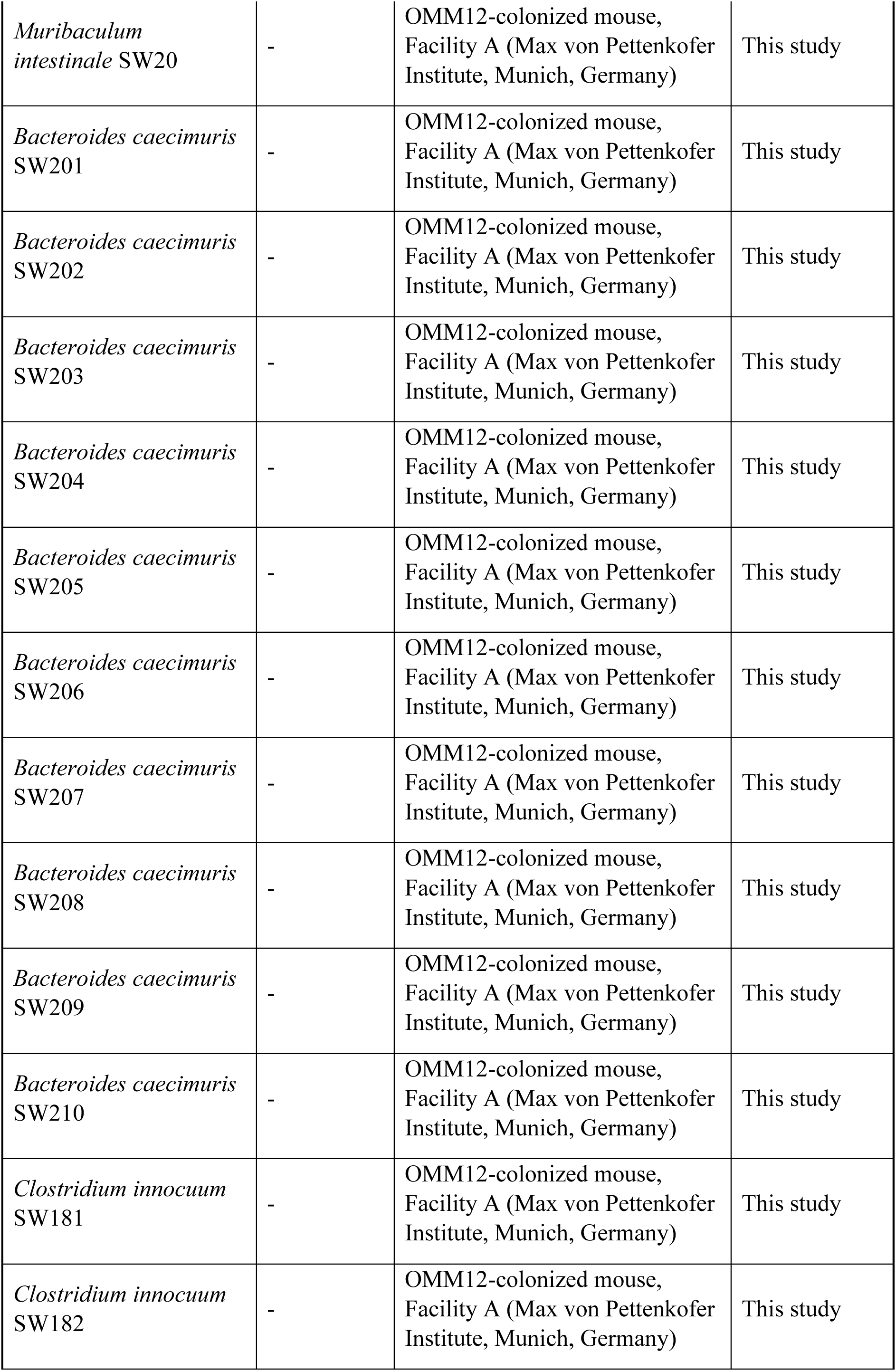
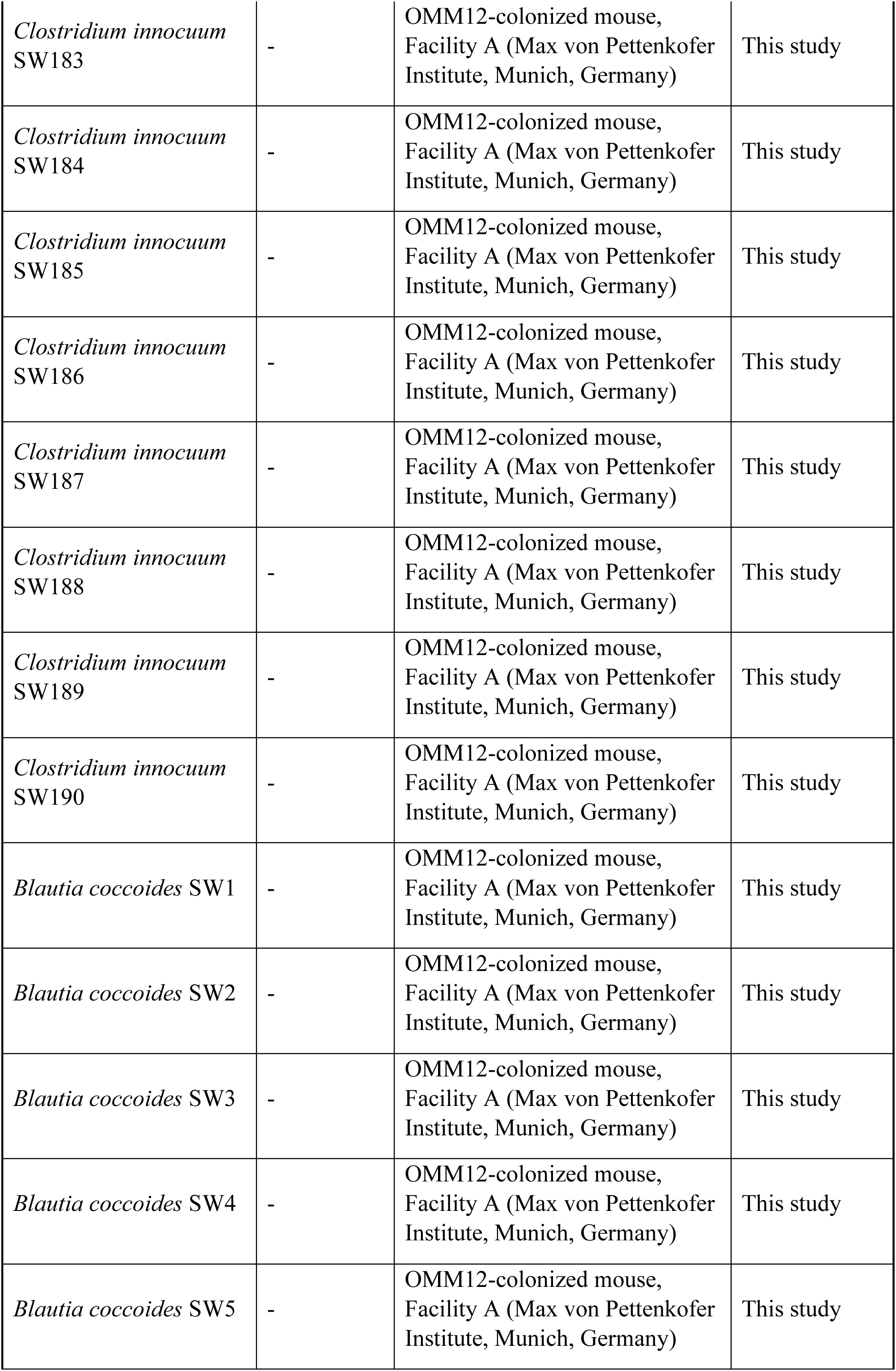
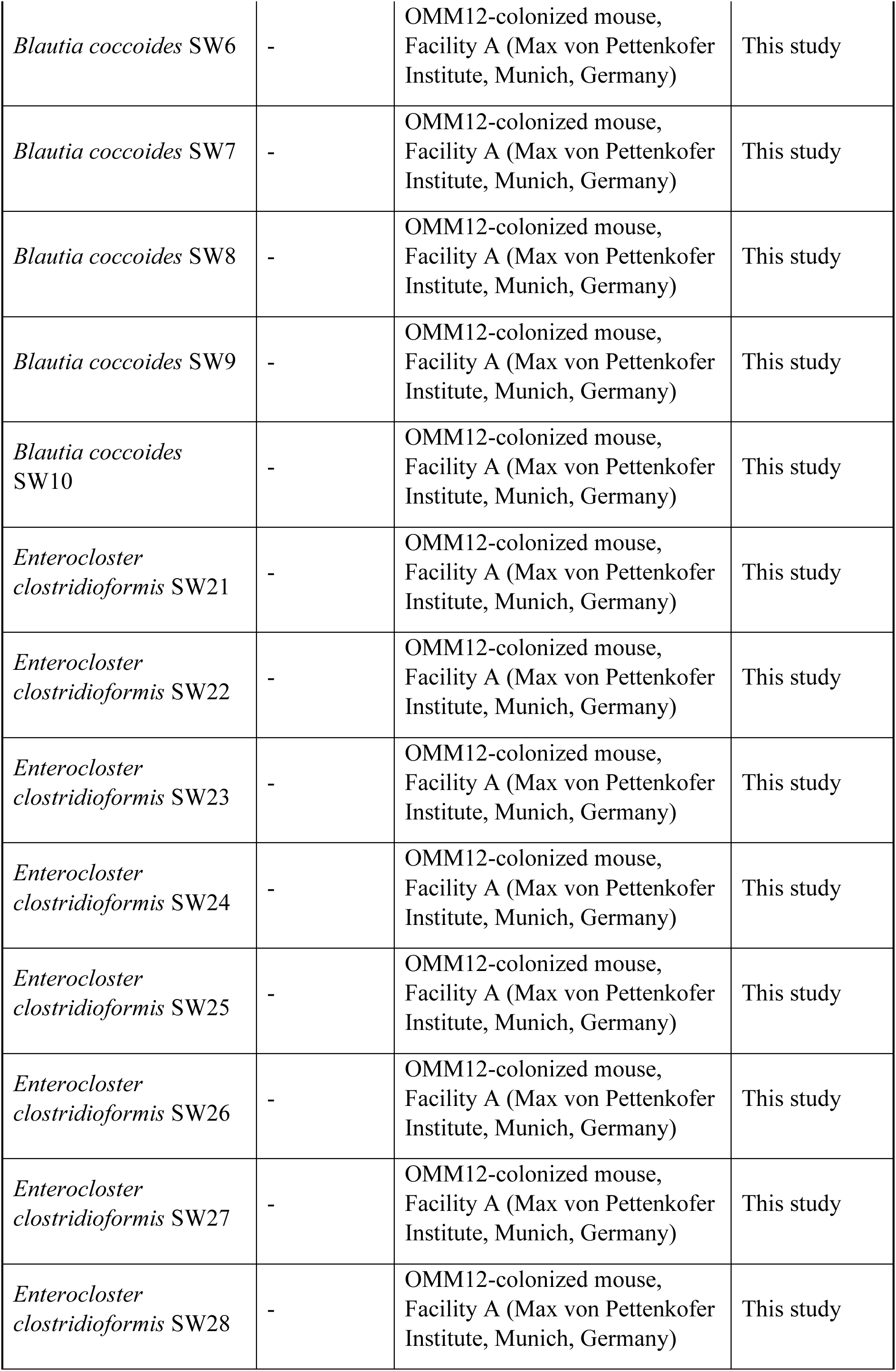
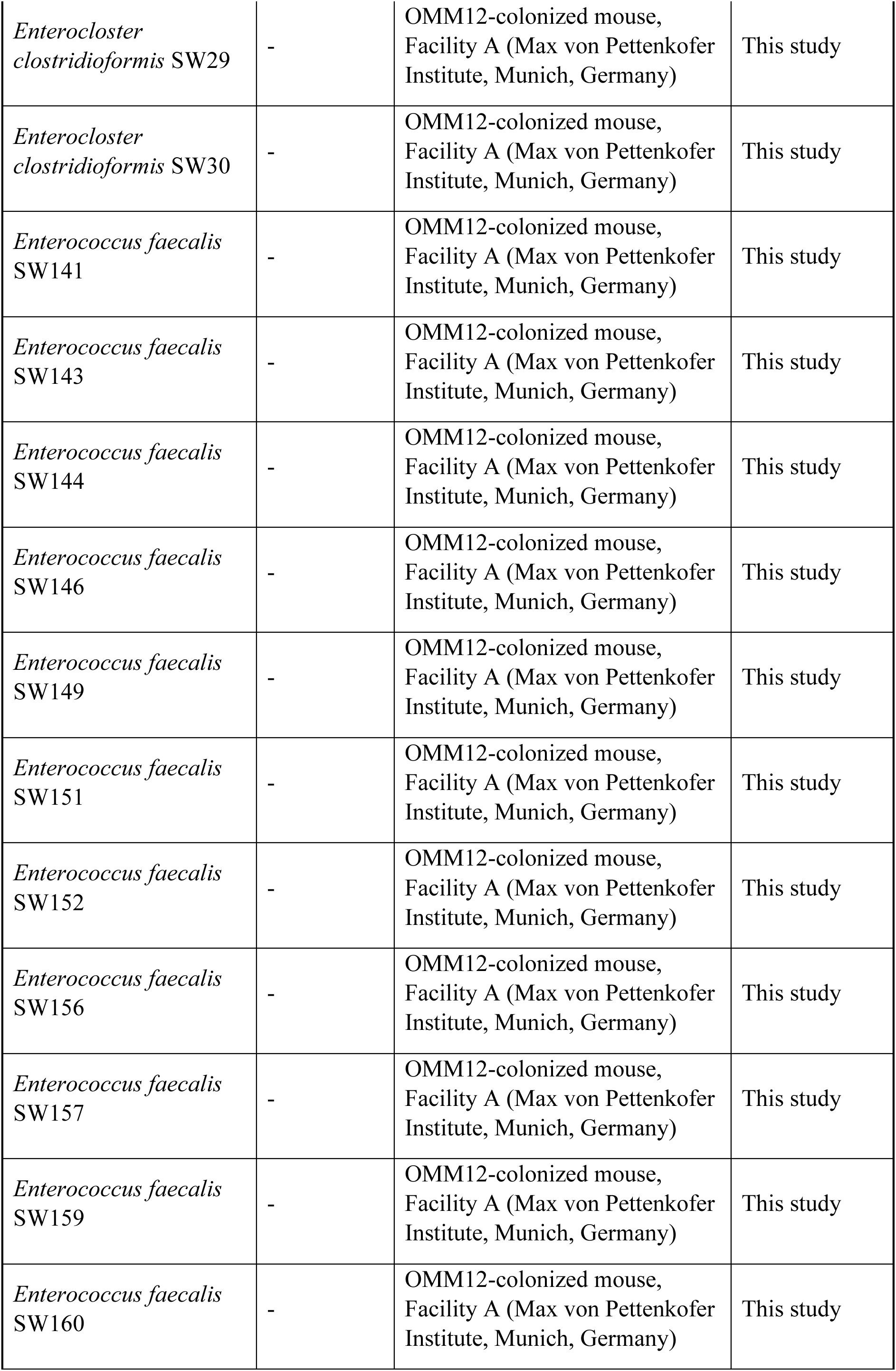
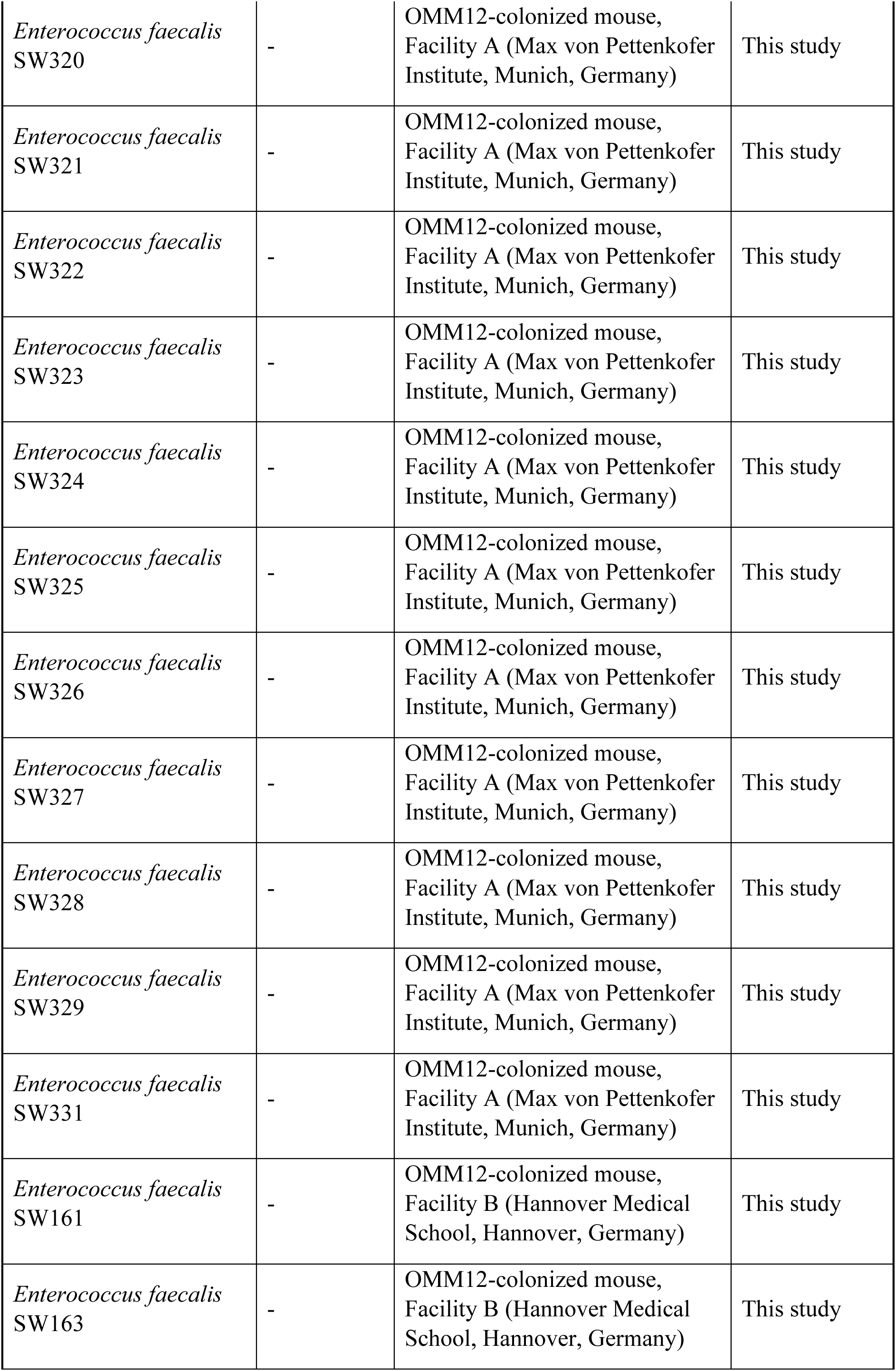
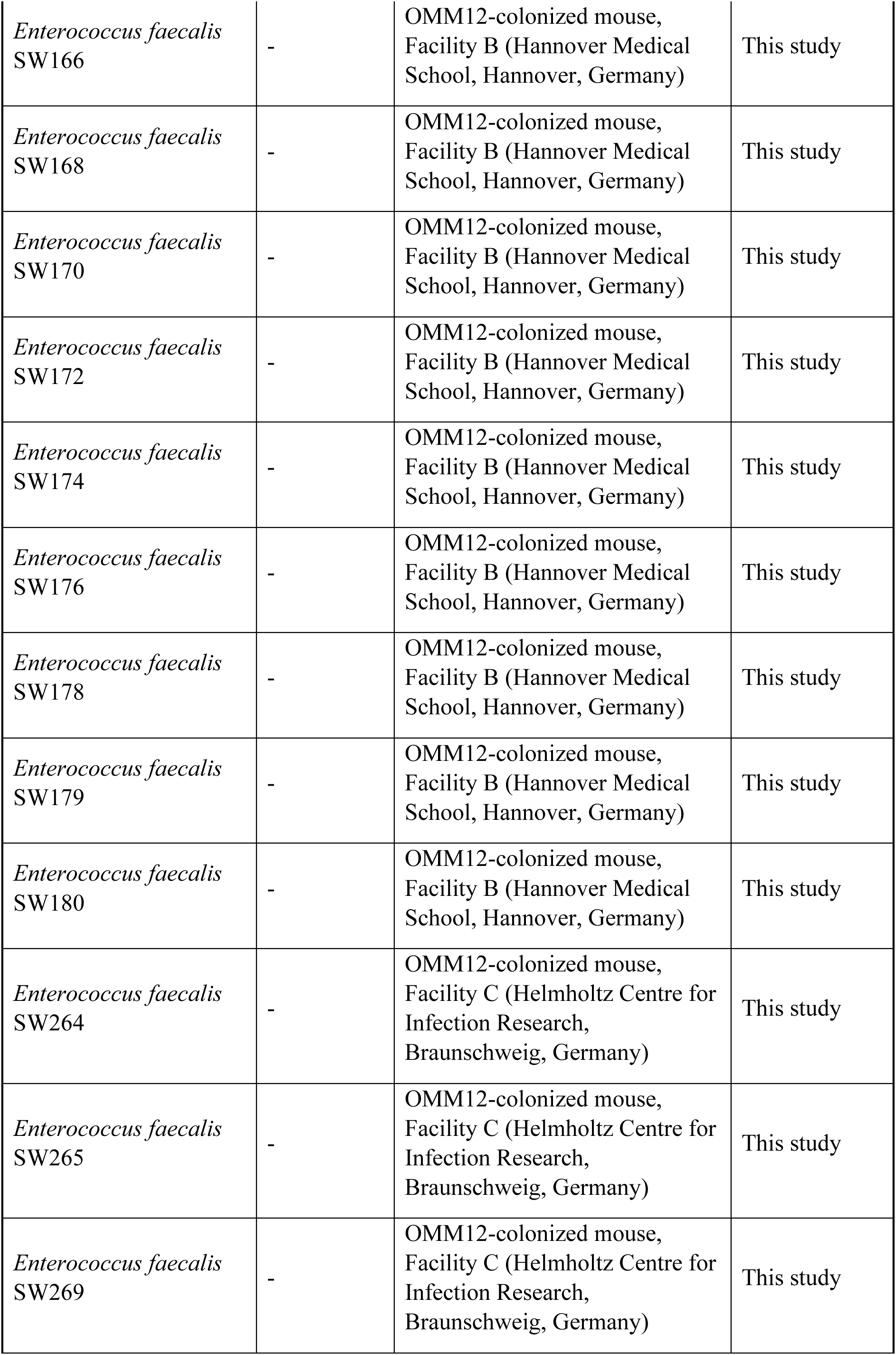
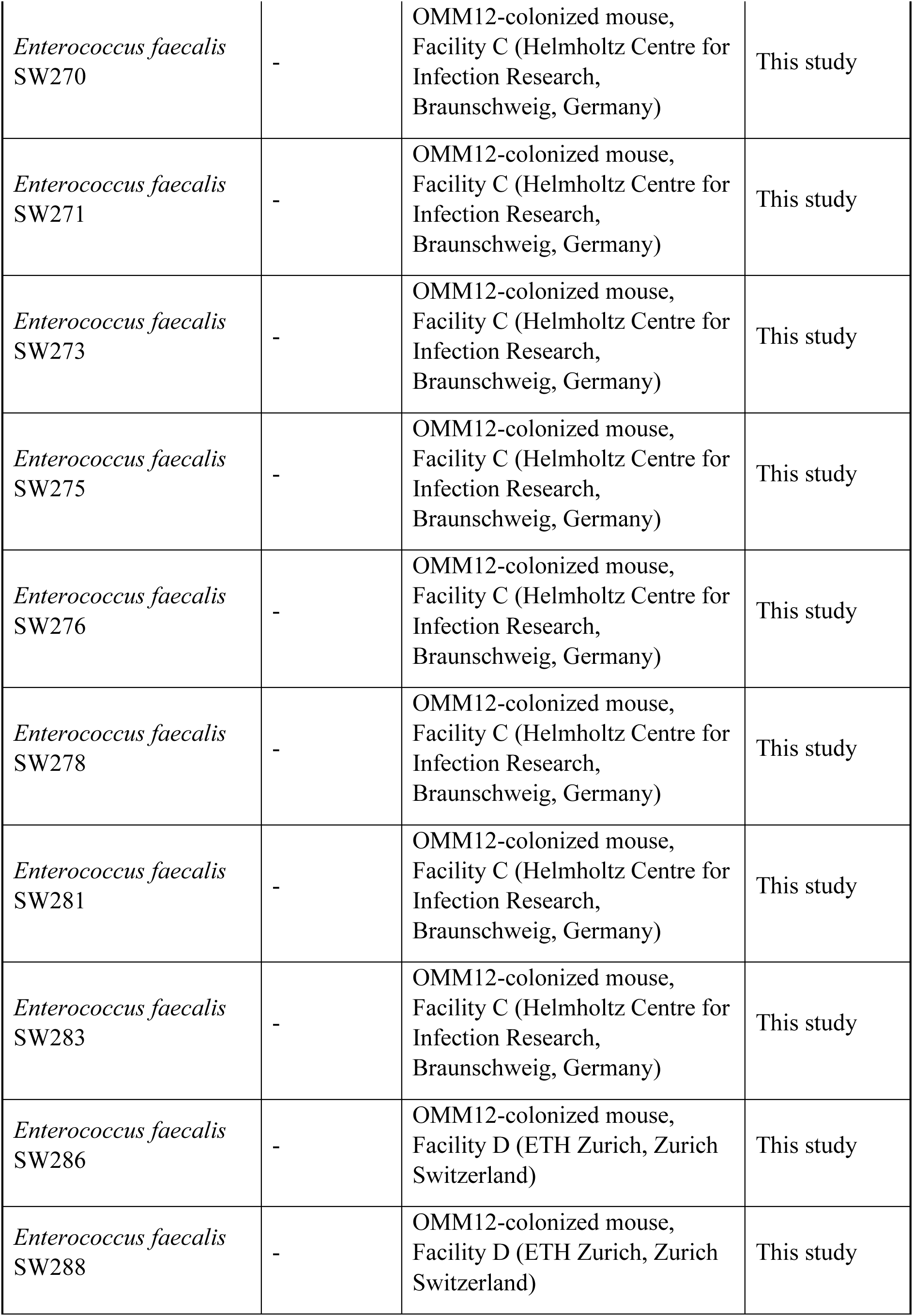
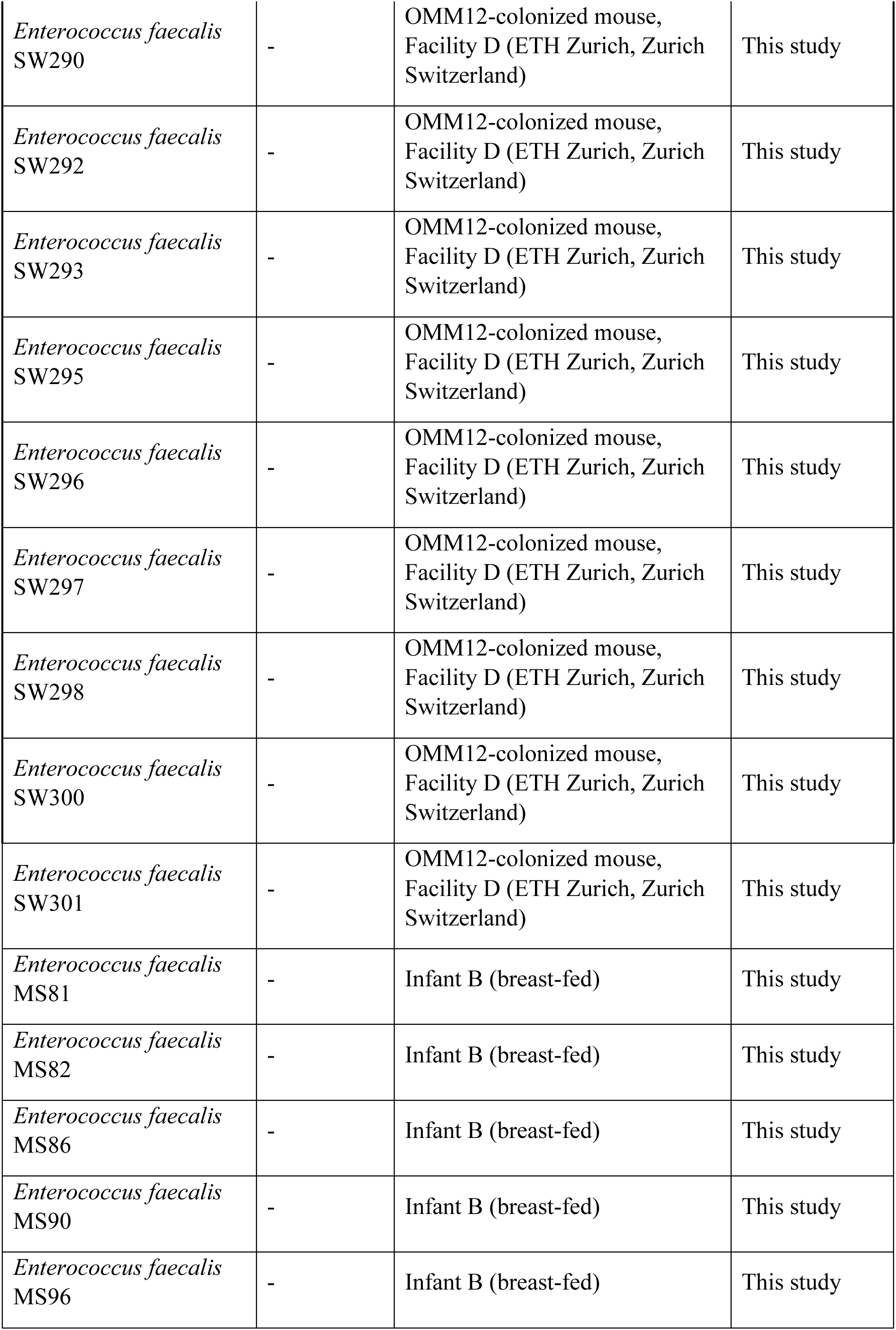
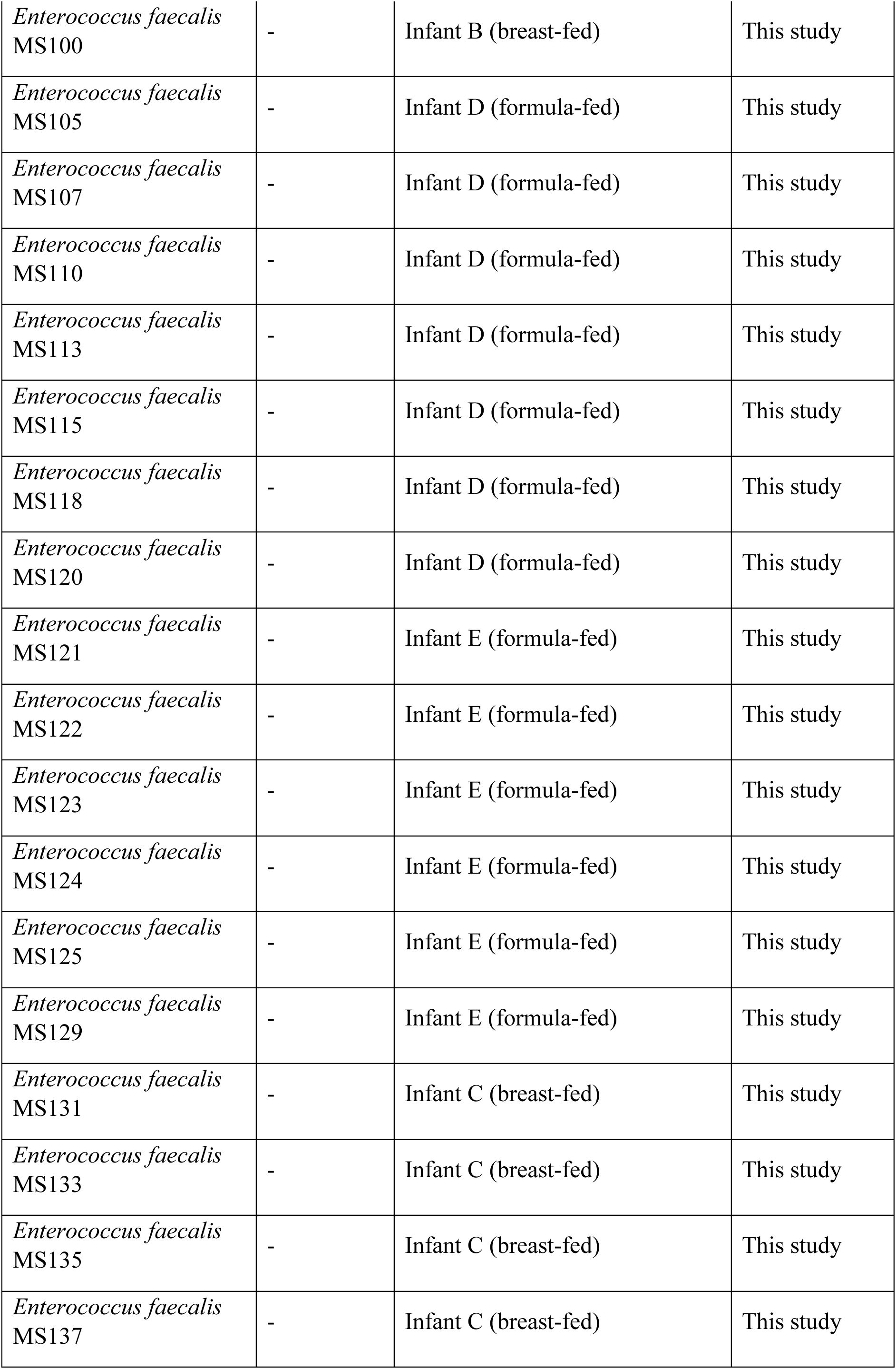
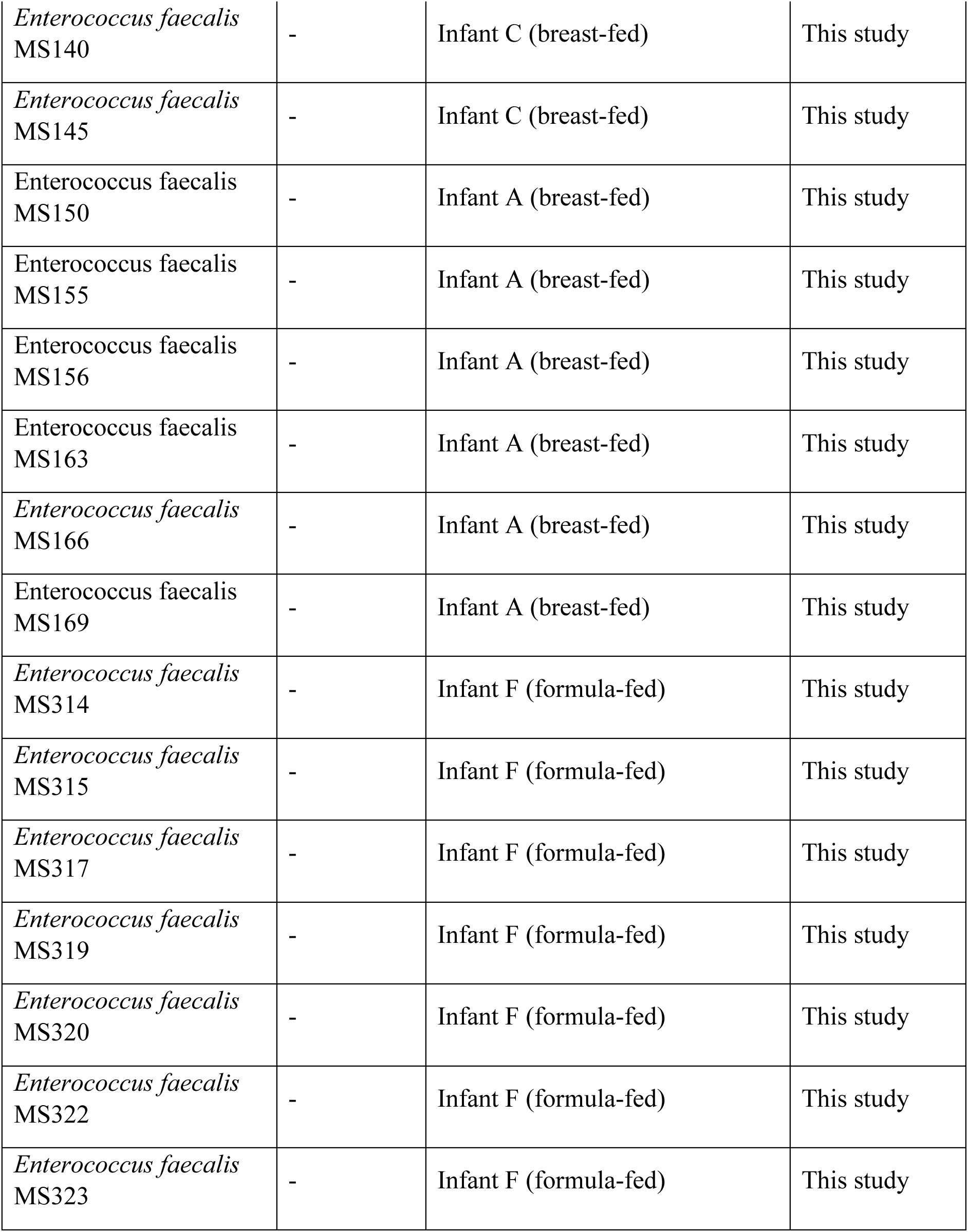
Bacterial strains used in this study.

**Table S2:**
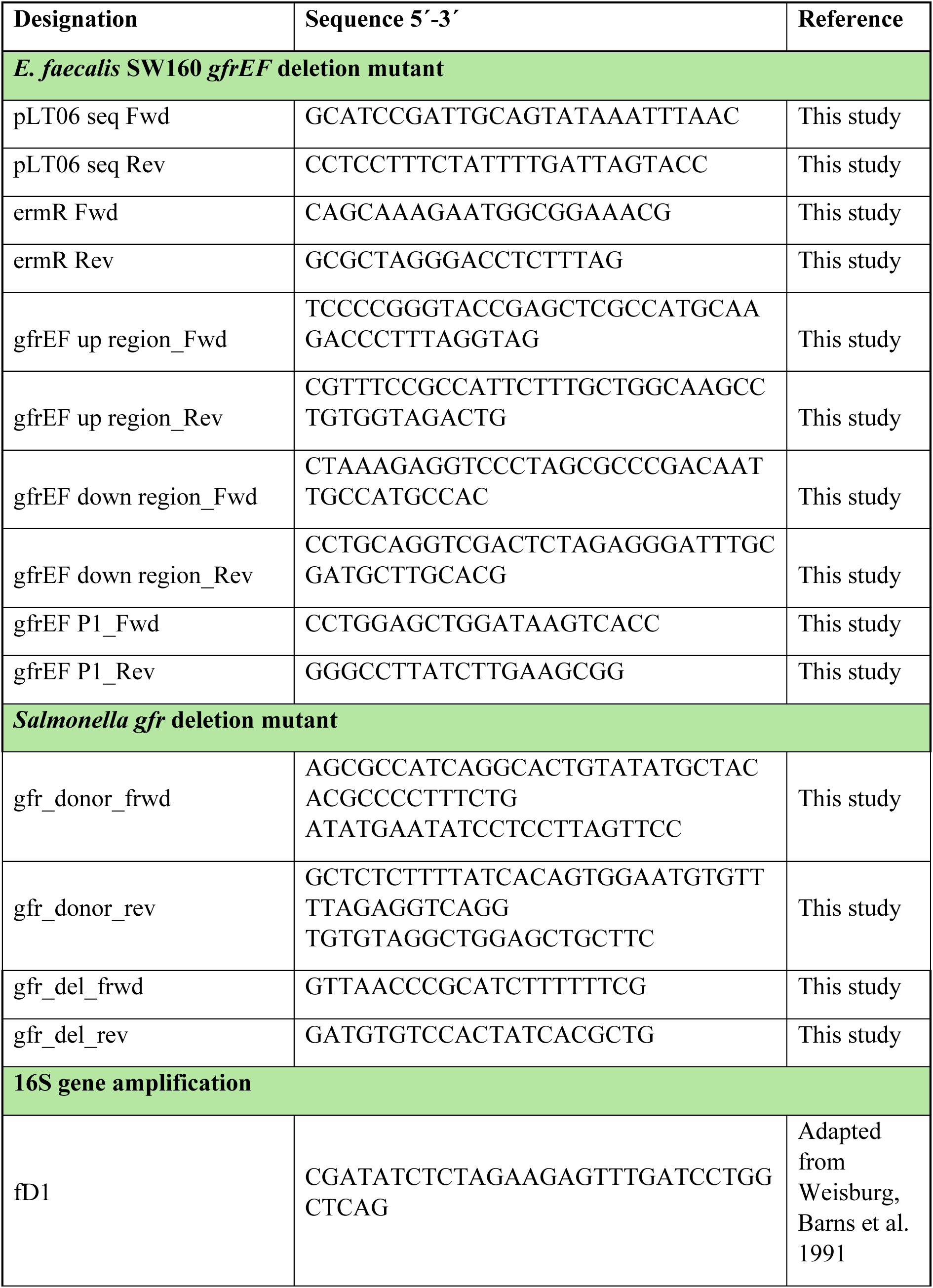
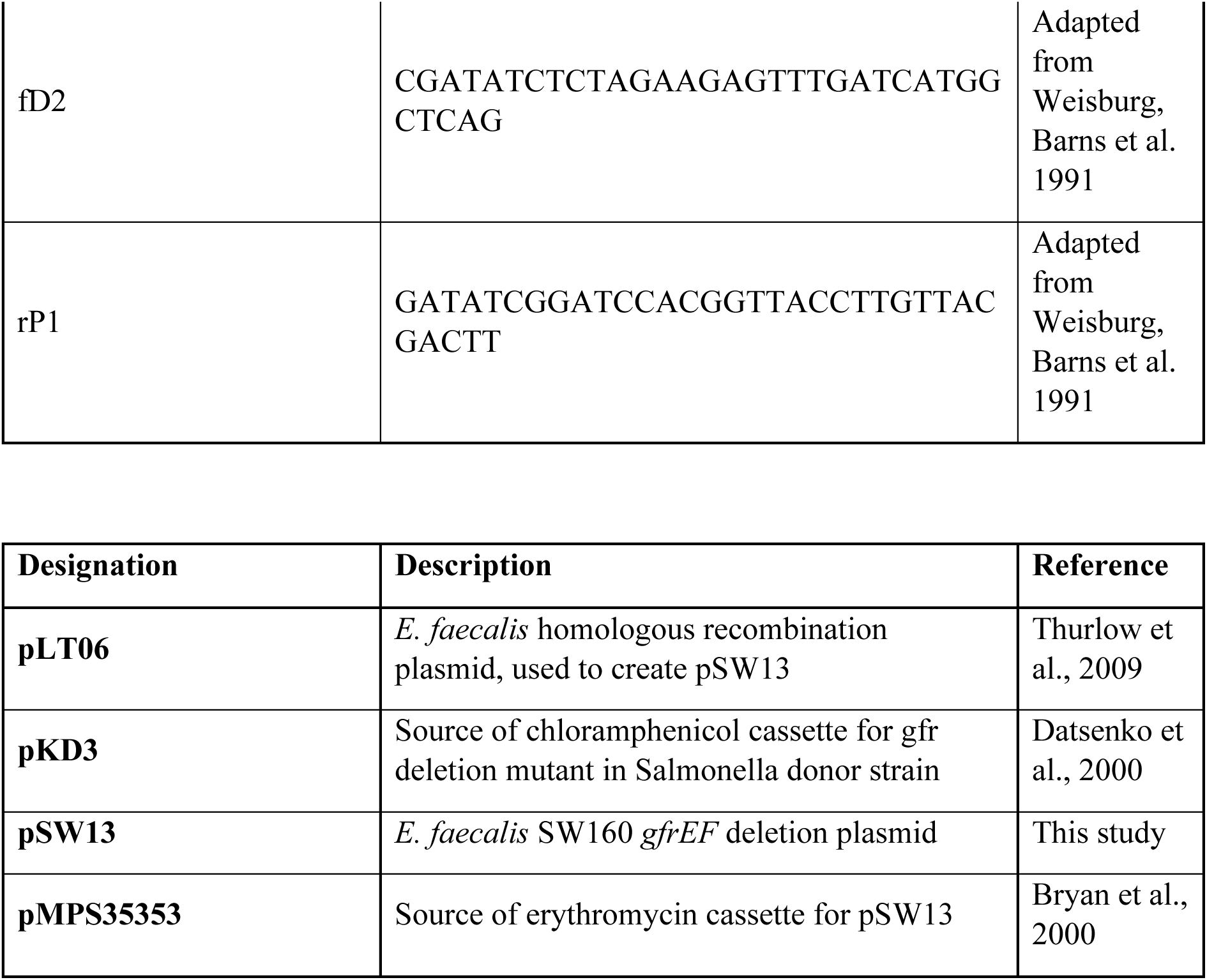
Primers and plasmids used for construction of bacterial mutants.

**Table S3:**
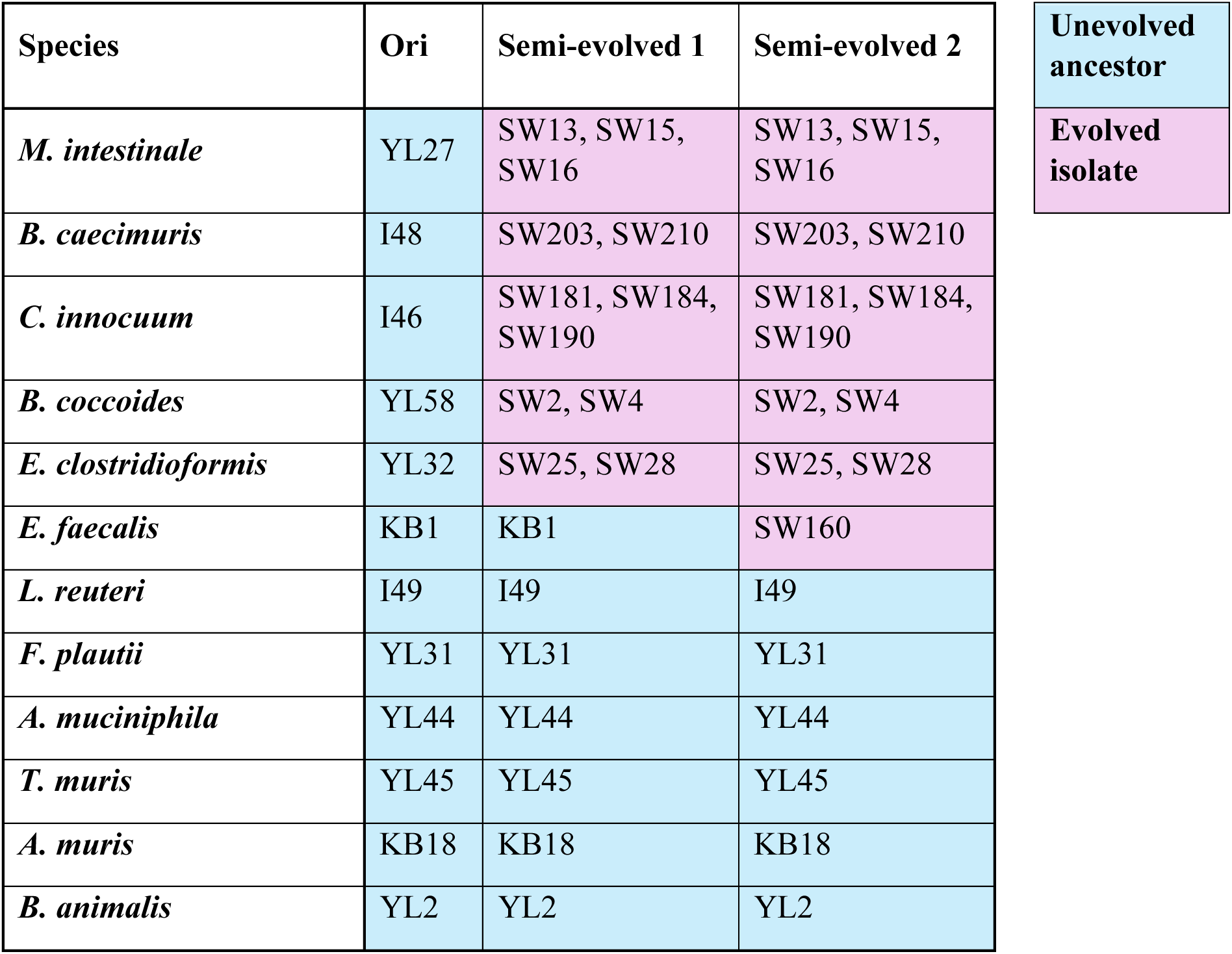
Composition of semi-evolved OMM^12^ communities.

**Table S4:**
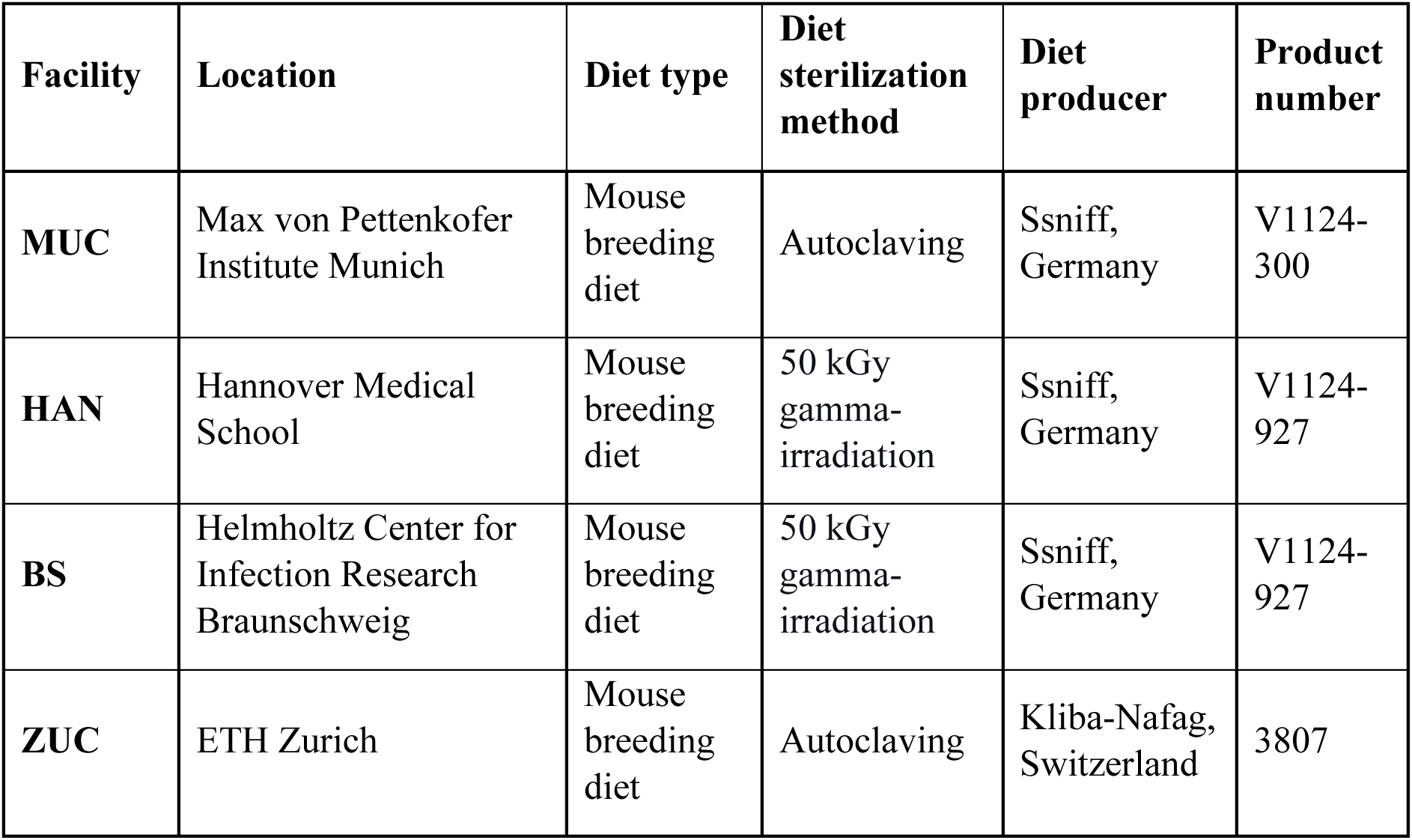
Mouse diet and sterilization method used in the different animal facilities.

## Notes

### Competing Interest Statement

The authors have declared no competing interest.

