## Supplementary material for "Gut microbiota within-host evolution enforces colonization resistance against enteric infection"

[14]Institute of Microbiology, Department of Biology, ETH Zurich, Zurich, Switzerland
[15]Institute for Laboratory Animal Science and Central Animal Facility, Hannover Medical School, Hannover, Germany
[16]Bavarian Center for Biomolecular Mass Spectrometry (BayBioMS), School of Life Sciences, Technical University of Munich, Freising, Germany
[17]Faculty of Biology, Ludwig Maximilian University, Planegg-Martinsried, Germany
[18]Laboratory for Mucosal Immunology, Institute of Food, Nutrition and Health, Department of Health Sciences and Technology, ETH Zurich, Zurich, Switzerland
[19]Sir William Dunn School of Pathology, University of Oxford, Oxford, U.K.
[20]Center for Individualized Infection Medicine, Hannover, Germany
[21]German Center for Infection Research (DZIF), Partner site Ludwig Maximilians University, Munich, Germany
[22]German Center for Infection Research (DZIF), Partner site Technical University of Munich, Munich, Germany
†These authors contributed equally to this work

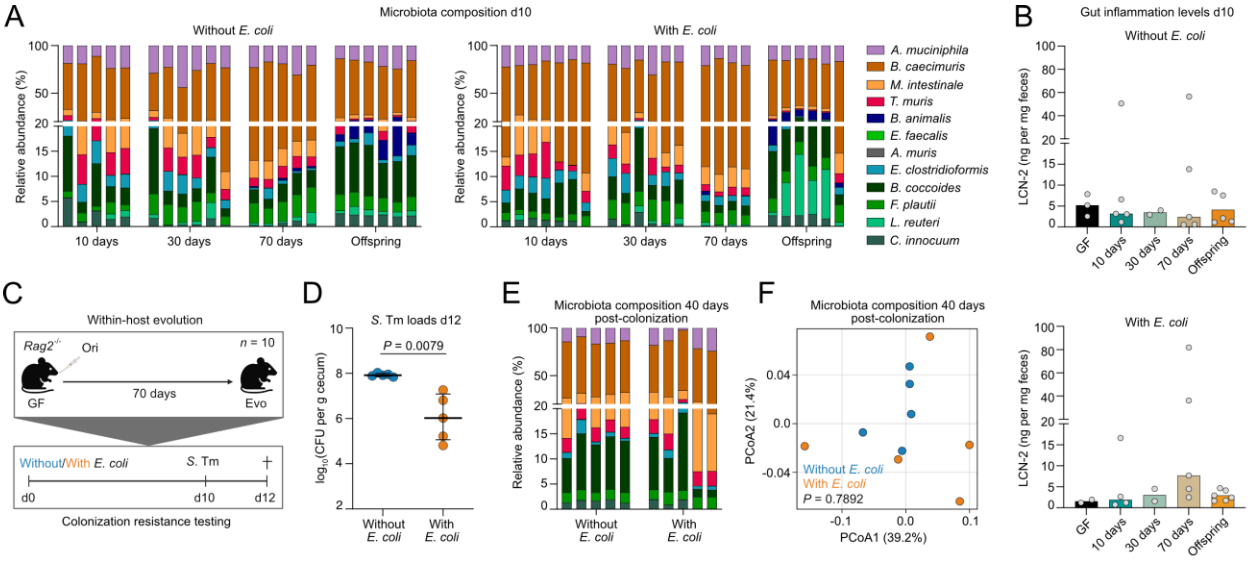


**Fig. S1: *E. coli*-mediated colonization resistance in long-term colonized OMM[12] mice is independent of gut inflammation and adaptive immunity**

**(A)** Fecal microbiota composition in mice shown in Fig. 1B on the day of *S*. Typhimurium (*S*. Tm) infection (d10). Relative abundances were determined via qPCR, quantifying species-specific 16S rRNA gene copies. Bars correspond to individual mice. **(B)** Feces lipocalin-2 concentrations in mice shown in Fig. 1B on the day of *S*. Tm infection (d10). Germ-free mice serve as controls. **(C)** Experimental scheme for assessing colonization resistance against *S*. Tm in mice lacking mature T- and B-cells. Germ-free *Rag2*$^{-/-}$ mice were colonized with the original OMM[12] community(*24*) for 70 days. Subsequently, mice were inoculated with either *E. coli* ($n$ = 5) or PBS ($n$ = 5). Colonization resistance was assessed as described in Fig. 1B. **(D)** Cecum *S*. Tm loads on day two post infection (d12). Data represent medians with Interquartile ranges, and statistical analysis was performed using a Mann-Whitney *U* test. The detection limit is 10 CFU per g. (**E** to **F)** Fecal microbiota composition 40 days after the colonization of *Rag2*$^{-/-}$ mice with the OMM[12] community. **(E)** Relative abundances were determined via qPCR, quantifying species-specific 16S rRNA gene copies. Bars correspond to individual mice. **(F)** Principal coordinate analysis was based on the Bray-Curtis dissimilarity distance matrix of relative OMM[12] abundance profiles. Statistical analysis was performed using a PERMANOVA with Bonferroni correction, comparing mice colonized with or without *E. coli*.

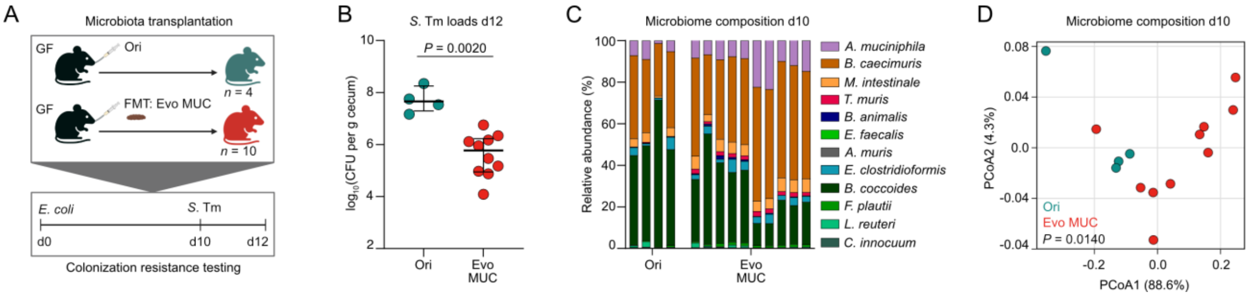


**Fig. S2: Transplantation of an independently evolved OMM$^{12}$ community provides colonization resistance against *S. Typhimurium*.**

**(A)** Assessment of colonization resistance against *S*. Typhimurium (*S*. Tm) in germ-free mice colonized with the original OMM$^{12}$ community or with an evolved OMM$^{12}$ community (Evo MUC) harvested from stably colonized mice (Evo MUC; equals "offspring" mice shown in Fig. 1). Original OMM$^{12}$ was freshly assembled from single-species cryo-stocks. Colonization resistance was assessed as described in Fig. 2B. **(B)** Cecum *S*. Tm loads on day two post-infection (d12). Data represent medians with interquartile ranges, and statistical analysis was performed using a Mann-Whitney *U* test. The detection limit is 10 CFU per g. **(C** and **D)** Fecal microbiota composition on the day of *S*. Tm infection (d10). **(C)** Relative abundances were determined via qPCR, quantifying species-specific 16S rRNA gene copies. Bars correspond to individual mice. **(D)** Principal coordinate analysis was based on the Bray-Curtis dissimilarity distance matrix of relative OMM$^{12}$ abundance profiles. Statistical analysis was performed using a PERMANOVA with Bonferroni correction.

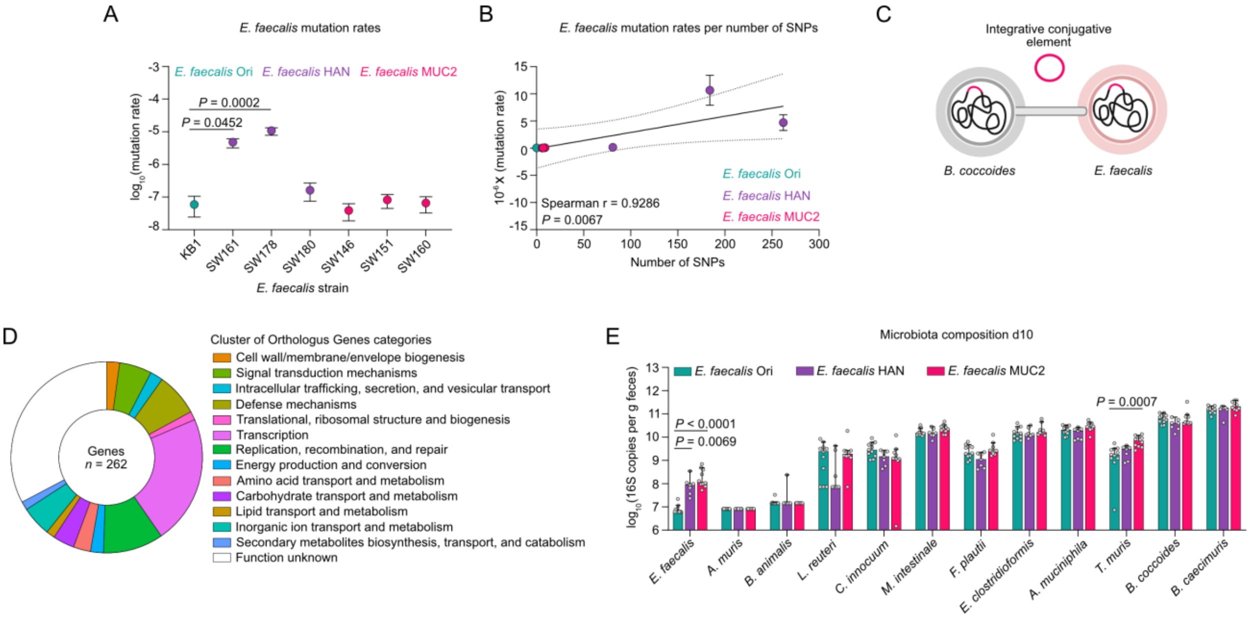


**Fig. S3: Characterization of evolved *E. faecalis* isolates from OMM$^{12}$ mice housed in different animal facilities.**

**(A)** Mutation rates of *E. faecalis* KB1 and select evolved isolates from two OMM$^{12}$ mice housed in different animal facilities (*E. faecalis* HAN, *E. faecalis* MUC2) were estimated using the Luria-Delbrück fluctuation assay, which quantifies the spontaneous development of rifampicin resistance. The isolates represent the phylogenetic clusters depicted in Fig. 3b. Statistical analysis was performed using a Dunnett's multiple comparison test, comparing each evolved isolate to *E. faecalis* KB1. Only *P*-values below 0.05 shown. **(B)** Correlation between mutation rates and the number of single nucleotide polymorphisms identified in each *E. faecalis* strain. Spearman's rank correlation and corresponding *P*-values are displayed, and error bars represent 95% confidence intervals. **(C)** Integrative conjugative element scheme. Several evolved *E. faecalis* isolates carry a 156 kb integrative conjugative element that originates from *B. coccoides* and is horizontally transferred during within-host evolution. **(D)** Proportions of genes per Clusters of Orthologous Genes (COG) category encoded on the integrative conjugative element. **(E)** Fecal microbiota composition on the day of *S.* Typhimurium (*S.* Tm) infection (d10) in mice shown in Fig. 3 c**,** d, e. Absolute abundances were determined via qPCR, quantifying species-specific 16S rRNA gene copies per g feces. Data represent medians with 95% confidence intervals, and statistical analyses were performed using Dunn's post-hoc tests, comparing mice colonized with the original *E.*

*faecalis* (*E. faecalis* Ori) to those colonized with evolved *E. faecalis* isolates (*E. faecalis* HAN, *E. faecalis* MUC2). Only *P*-values below 0.05 shown.

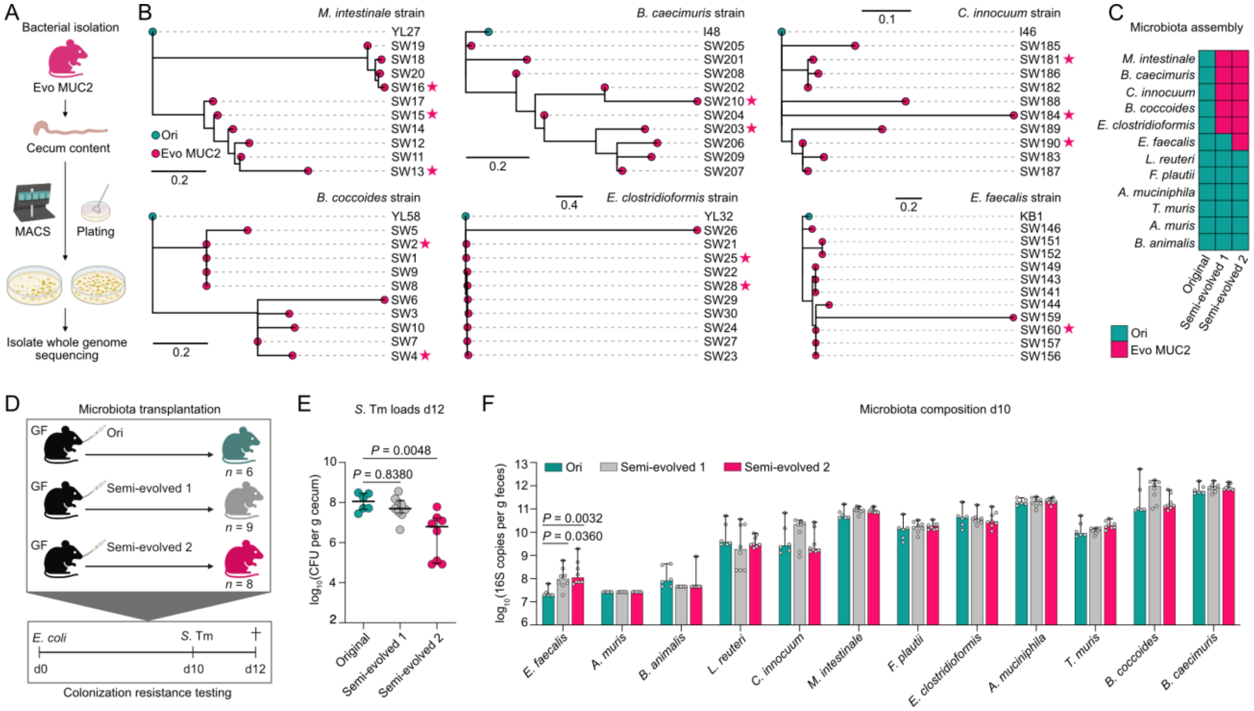


**Fig. S4: Evolved isolates of OMM$^{12}$ species other than *E. faecalis* fail to confer colonization resistance against *S. Typhimurium*.**

**(A)** Bacterial isolation scheme. Evolved isolates of six different OMM$^{12}$ species were obtained from the cecum content of a single long-term colonized OMM$^{12}$ mouse (Evo MUC2). Isolates were subjected to short-read whole genome sequencing. *E. faecalis* isolates are also shown in Fig. 3b. MACS, magnetic-activated cell sorting. **(B)** Single nucleotide polymorphism-based phylogenetic trees of evolved isolates (*n* = 10-11 per species) and original strains. A polymorphism allele frequency cut-off of 0.8 was applied. The tree was created using the Random Accelerated Maximum Likelihood method, with branch lengths representing the substitutions per variable site. Stars highlight strains used in (D). **(C)** Semi-evolved microbiota assembly scheme. The original OMM$^{12}$ community(*26*) was freshly assembled from single-species cryo-stocks, while semi-evolved communities were assembled using evolved isolates from several species (magenta) along with original strains of other species (turquoise). **(D)** Assessment of colonization resistance against *S*. Typhimurium (*S*. Tm) in germ-free mice colonized with original or semi-evolved OMM$^{12}$ communities. Colonization resistance was assessed as described in Fig. 3B. **(E)** Cecum *S*. Tm loads on day two post-infection (d12). Data represent medians with interquartile ranges, and statistical analysis was performed using a Dunn's post-hoc test. The detection limit is 10 CFU per g. **(F)** Fecal microbiota composition on the day of *S*. Tm infection (d10). Absolute abundances were determined via qPCR, quantifying species-specific 16S rRNA gene copies per g feces. Data represent medians with 95% confidence intervals, and statistical analyses were performed using Dunn's post-hoc tests, comparing the original with each semi-evolved $OMM^{12}$ community. Only *P*-values below 0.05 are shown.

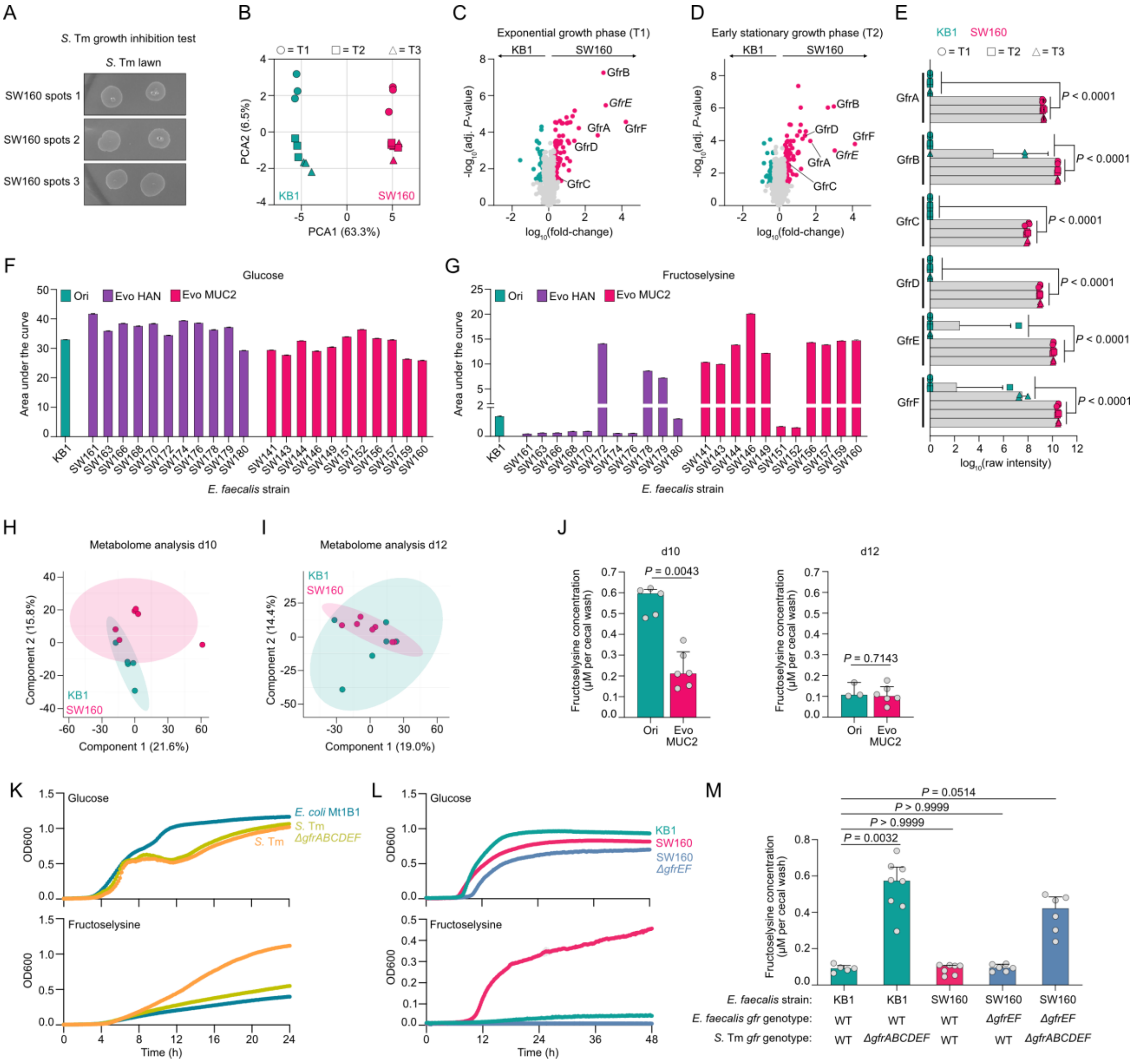


**Fig. S5: *S. Typhimurium* and evolved *E. faecalis* compete for fructoselysine in the gut.**
**(A)** Growth inhibition assay. *E. faecalis* SW160 was spotted on a lawn of *S.* Typhimurium (*S.* Tm). **B-E)** Data from the experiment described in Fig. 4E. *E. faecalis* strains were grown in AF liquid medium, and proteomes were analyzed during the exponential (T1), early stationary (T2), or late stationary (T3) growth phases. **(B)** Principal component analysis of proteomes. **(C** to **D)** Volcano plots showing proteins overrepresented in *E. faecalis* KB1 (turquoise) or SW160 (magenta). Significance cut-offs were $\log_{10}(2)$ and $-\log_{10}(0.05)$ for fold-changes and Benjamini-Hochberg adjusted t-test *P*-values, respectively. Fructoselysine operon-encoded proteins are highlighted. **(E)** Raw mass spectrometric intensities of fructoselysine operon-encoded proteins. Bars represent

means with standard deviation, and statistical analyses were performed using Mann-Whitney *U* tests. **(F** and **G)** Area under the curve based on the growth of *E. faecalis* strains in minimal medium with glucose **(F)** or fructoselysine **(G)** as the sole carbon source. Data represent means with 95% confidence intervals. **(H** and **I)** Same experiment as shown in Fig. 4e. Metabolomes of cecal washes were analyzed in mice colonized with original *E. faecalis* KB1 or evolved *E. faecalis* SW160 on the day of *S*. Tm infection (d10; KB1: $n$ = 5; SW160: $n$ = 6; **H**) or on day two post-infection (d12; KB1: $n$ = 6; SW160: $n$ = 7; **I**). Principal coordinates analysis was performed on Bray-Curtis distance matrices. **(J)** Same experimental setup as in Fig. 4e, but comparing the metabolomes of mice colonized with an original OMM$^{12}$ community(*24*) or an evolved community (Evo MUC2). Fructoselysine levels in cecal washes on the day of *S*. Tm infection (d10; Ori: $n$ = 5; Evo MUC2: $n$ = 6) and two days post-infection (d12; Ori: $n$ = 6; Evo MUC2: $n$ = 6). Data represent medians with Interquartile ranges, and statistical analyses were performed using Mann-Whitney *U* tests. **(K** and **L)** Growth curves of *S*. Tm, *S*. Tm *ΔgfrABCDEF*, and *E. coli* **(K)** or of *E. faecalis* KB1, *E. faecalis* SW160, and *E. faecalis* SW160 *ΔgfrEF* **(L)** in minimal medium with glucose or fructoselysine as the sole carbon source. **(M)** Same experiment as shown in Fig. 4I. Fructoselysine levels in cecal washes on day two post-infection (d12). Data represent medians with interquartile ranges, and statistical analysis was performed using a Dunn's post-hoc test.

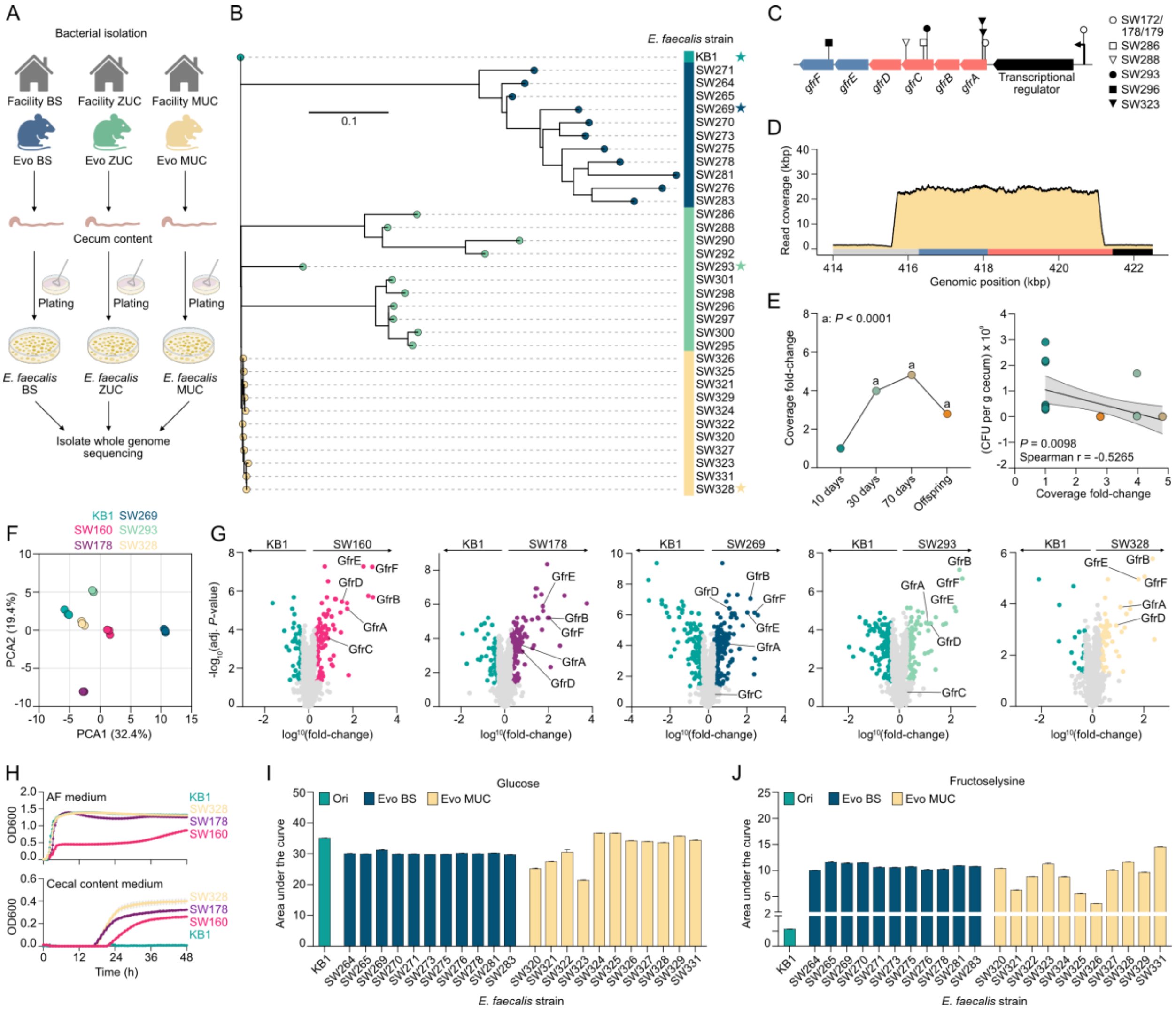


**Fig. S6: Different within-host evolutionary trajectories converge to enable *E. faecalis* fructoselysine utilization**

**(A)** *E. faecalis* isolation scheme. Evolved *E. faecalis* were isolated from the cecum contents of long-term colonized OMM[12] mice (Evo BS, Evo ZUC, Evo MUC) from three different animal facilities. The Evo MUC mouse is shown in Fig. 1 as "offspring" and in Fig. S2. Isolates were subjected to short-read whole genome sequencing ($n = 1$ mouse per facility). **(B)** Single nucleotide polymorphism-based phylogenetic tree of evolved *E. faecalis* isolates ($n = 11$ per mouse) and original *E. faecalis* KB1. A polymorphism allele frequency cut-off of 0.8 was applied. The tree was created using the Random Accelerated Maximum Likelihood method, and branch lengths represent substitutions per variable site ($n = 2{,}703$). Stars highlight strains used in (F) and (G). **(C)** Fructoselysine operon single nucleotide polymorphisms in *E. faecalis* isolates from this study.

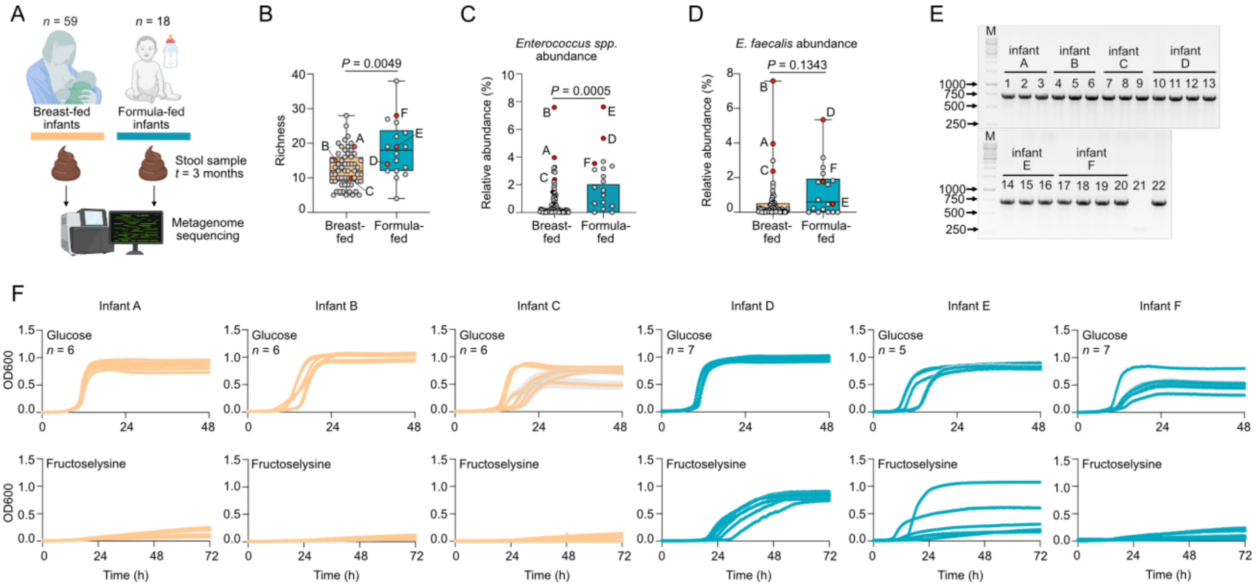


**Fig. S7: Characterization of *E. faecalis* isolates from breast- and formula-fed infants.** **(A)** Study scheme. Stool samples from three-month-old breast- or formula-fed infants were collected and subjected to metagenome sequencing. **(B)** Gut microbiota richness among all breast- and formula-fed infants included in the study. Infants used for *E. faecalis* isolation are highlighted (breast-fed: A, B, C; formula-fed: D, E, F). **(C and D)** Metagenome-based relative abundance of *Enterococcus* species (C) and *E. faecalis* (D) across all breast- and formula-fed infants included in the study. Statistical analyses were performed using Mann-Whitney *U* tests. **E)** PCR-generated *gfrEF* gene amplicons in *E. faecalis* isolates from breast- (A, B, C) and formula-fed (D, E, F) infants ($n$ = 3-4 isolates per infant). Numbers correspond to different samples, and results are representative of all obtained *E. faecalis* isolates. Genomic DNA from *E. faecalis* SW160 Δ*gfrEF* and *E. faecalis* SW160 was used as templates for PCRs 21 and 22, respectively. The expected amplicon size is 746 bp. M, DNA marker. **(F)** Growth curves of all *E. faecalis* isolates per infant in minimal medium with glucose or fructoselysine as the sole carbon source. Corresponding areas under the curve are shown in Fig. 5B.

**Table S1: Bacterial strains used in this study.**

| Strain name | DSMZ ID (if available) | Origin (if isolated in this study); genotype (if generated in this study) | Reference |
|---|---|---|---|
| Enterococcus faecalis KB1 | DSM 32036 | - | Brugiroux et al, 2016 |
| *Bifidobacterium animalis* YL2 | DSM 26074 | - | Brugiroux et al, 2016 |
| *Acutalibacter muris* KB18 | DSM 26090 | - | Brugiroux et al, 2016 |
| *Muribaculum intestinale* YL27 | DSM 28989 | - | Brugiroux et al, 2016 |
| *Flavonifractor plautii* YL31 | DSM 26117 | - | Brugiroux et al, 2016 |
| *Enterocloster clostridioformis* YL32 | DSM 26114 | - | Brugiroux et al, 2016 |
| *Akkermansia muciniphila* YL44 | DSM 26127 | - | Brugiroux et al, 2016 |
| *Turicimonas muris* YL45 | DSM 26109 | - | Brugiroux et al, 2016 |
| *Clostridium innocuum* I46 | DSM 26113 | - | Brugiroux et al, 2016 |
| *Bacteroides caecimuris* I48 | DSM 26085 | - | Brugiroux et al, 2016 |
| *Limosilactobacillus reuteri* I49 | DSM 32035 | - | Brugiroux et al, 2016 |
| *Blautia coccoides* YL58 | DSM 26115 | - | Brugiroux et al, 2016 |
| *Escherichia coli* Mt1B1 | DSM 28618 | - | Brugiroux et al, 2016 |
| *Salmonella enterica* serovar Thyphimurium$^{avir}$ Z5537 | - | *Salmonella enterica* serovar Thyphimurium$^{avir}$ Z5537 *gfrABCDEF::cat* | This study |
| *Salmonella enterica* serovar Thyphimurium$^{avir}$ M2707 | - | - | Maier et al, 2013 |
| *Salmonella enterica* serovar Thyphimurium$^{avir}$ gfrABCDEF | - | *Salmonella enterica* serovar Thyphimurium$^{avir}$ M2707 *gfrABCDEF::cat* | This study |
| *Enterococcus faecalis* gfrEF | - | *Enterococcus faecalis* SW160 *gfrEF::cat* | This study |

| | | | |
|---|---|---|---|
| *Muribaculum intestinale* SW11 | - | OMM12-colonized mouse, Facility A (Max von Pettenkofer Institute, Munich, Germany) | This study |
| *Muribaculum intestinale* SW12 | - | OMM12-colonized mouse, Facility A (Max von Pettenkofer Institute, Munich, Germany) | This study |
| *Muribaculum intestinale* SW13 | - | OMM12-colonized mouse, Facility A (Max von Pettenkofer Institute, Munich, Germany) | This study |
| *Muribaculum intestinale* SW14 | - | OMM12-colonized mouse, Facility A (Max von Pettenkofer Institute, Munich, Germany) | This study |
| *Muribaculum intestinale* SW15 | - | OMM12-colonized mouse, Facility A (Max von Pettenkofer Institute, Munich, Germany) | This study |
| *Muribaculum intestinale* SW16 | - | OMM12-colonized mouse, Facility A (Max von Pettenkofer Institute, Munich, Germany) | This study |
| *Muribaculum intestinale* SW17 | - | OMM12-colonized mouse, Facility A (Max von Pettenkofer Institute, Munich, Germany) | This study |
| *Muribaculum intestinale* SW18 | - | OMM12-colonized mouse, Facility A (Max von Pettenkofer Institute, Munich, Germany) | This study |
| *Muribaculum intestinale* SW19 | - | OMM12-colonized mouse, Facility A (Max von Pettenkofer Institute, Munich, Germany) | This study |
| *Muribaculum intestinale* SW20 | - | OMM12-colonized mouse, Facility A (Max von Pettenkofer Institute, Munich, Germany) | This study |
| *Bacteroides caecimuris* SW201 | - | OMM12-colonized mouse, Facility A (Max von Pettenkofer Institute, Munich, Germany) | This study |
| *Bacteroides caecimuris* SW202 | - | OMM12-colonized mouse, Facility A (Max von Pettenkofer Institute, Munich, Germany) | This study |
| *Bacteroides caecimuris* SW203 | - | OMM12-colonized mouse, Facility A (Max von Pettenkofer Institute, Munich, Germany) | This study |
| *Bacteroides caecimuris* SW204 | - | OMM12-colonized mouse, Facility A (Max von Pettenkofer Institute, Munich, Germany) | This study |
| *Bacteroides caecimuris* SW205 | - | OMM12-colonized mouse, Facility A (Max von Pettenkofer Institute, Munich, Germany) | This study |

| | | | |
|---|---|---|---|
| *Bacteroides caecimuris* SW206 | - | OMM12-colonized mouse, Facility A (Max von Pettenkofer Institute, Munich, Germany) | This study |
| *Bacteroides caecimuris* SW207 | - | OMM12-colonized mouse, Facility A (Max von Pettenkofer Institute, Munich, Germany) | This study |
| *Bacteroides caecimuris* SW208 | - | OMM12-colonized mouse, Facility A (Max von Pettenkofer Institute, Munich, Germany) | This study |
| *Bacteroides caecimuris* SW209 | - | OMM12-colonized mouse, Facility A (Max von Pettenkofer Institute, Munich, Germany) | This study |
| *Bacteroides caecimuris* SW210 | - | OMM12-colonized mouse, Facility A (Max von Pettenkofer Institute, Munich, Germany) | This study |
| *Clostridium innocuum* SW181 | - | OMM12-colonized mouse, Facility A (Max von Pettenkofer Institute, Munich, Germany) | This study |
| *Clostridium innocuum* SW182 | - | OMM12-colonized mouse, Facility A (Max von Pettenkofer Institute, Munich, Germany) | This study |
| *Clostridium innocuum* SW183 | - | OMM12-colonized mouse, Facility A (Max von Pettenkofer Institute, Munich, Germany) | This study |
| *Clostridium innocuum* SW184 | - | OMM12-colonized mouse, Facility A (Max von Pettenkofer Institute, Munich, Germany) | This study |
| *Clostridium innocuum* SW185 | - | OMM12-colonized mouse, Facility A (Max von Pettenkofer Institute, Munich, Germany) | This study |
| *Clostridium innocuum* SW186 | - | OMM12-colonized mouse, Facility A (Max von Pettenkofer Institute, Munich, Germany) | This study |
| *Clostridium innocuum* SW187 | - | OMM12-colonized mouse, Facility A (Max von Pettenkofer Institute, Munich, Germany) | This study |
| *Clostridium innocuum* SW188 | - | OMM12-colonized mouse, Facility A (Max von Pettenkofer Institute, Munich, Germany) | This study |
| *Clostridium innocuum* SW189 | - | OMM12-colonized mouse, Facility A (Max von Pettenkofer Institute, Munich, Germany) | This study |
| *Clostridium innocuum* SW190 | - | OMM12-colonized mouse, Facility A (Max von Pettenkofer Institute, Munich, Germany) | This study |

| | | | |
|---|---|---|---|
| *Blautia coccoides* SW1 | - | OMM12-colonized mouse, Facility A (Max von Pettenkofer Institute, Munich, Germany) | This study |
| *Blautia coccoides* SW2 | - | OMM12-colonized mouse, Facility A (Max von Pettenkofer Institute, Munich, Germany) | This study |
| *Blautia coccoides* SW3 | - | OMM12-colonized mouse, Facility A (Max von Pettenkofer Institute, Munich, Germany) | This study |
| *Blautia coccoides* SW4 | - | OMM12-colonized mouse, Facility A (Max von Pettenkofer Institute, Munich, Germany) | This study |
| *Blautia coccoides* SW5 | - | OMM12-colonized mouse, Facility A (Max von Pettenkofer Institute, Munich, Germany) | This study |
| *Blautia coccoides* SW6 | - | OMM12-colonized mouse, Facility A (Max von Pettenkofer Institute, Munich, Germany) | This study |
| *Blautia coccoides* SW7 | - | OMM12-colonized mouse, Facility A (Max von Pettenkofer Institute, Munich, Germany) | This study |
| *Blautia coccoides* SW8 | - | OMM12-colonized mouse, Facility A (Max von Pettenkofer Institute, Munich, Germany) | This study |
| *Blautia coccoides* SW9 | - | OMM12-colonized mouse, Facility A (Max von Pettenkofer Institute, Munich, Germany) | This study |
| *Blautia coccoides* SW10 | - | OMM12-colonized mouse, Facility A (Max von Pettenkofer Institute, Munich, Germany) | This study |
| *Enterocloster clostridioformis* SW21 | - | OMM12-colonized mouse, Facility A (Max von Pettenkofer Institute, Munich, Germany) | This study |
| *Enterocloster clostridioformis* SW22 | - | OMM12-colonized mouse, Facility A (Max von Pettenkofer Institute, Munich, Germany) | This study |
| *Enterocloster clostridioformis* SW23 | - | OMM12-colonized mouse, Facility A (Max von Pettenkofer Institute, Munich, Germany) | This study |
| *Enterocloster clostridioformis* SW24 | - | OMM12-colonized mouse, Facility A (Max von Pettenkofer Institute, Munich, Germany) | This study |
| *Enterocloster clostridioformis* SW25 | - | OMM12-colonized mouse, Facility A (Max von Pettenkofer Institute, Munich, Germany) | This study |

| | | | |
|---|---|---|---|
| *Enterocloster clostridioformis* SW26 | - | OMM12-colonized mouse, Facility A (Max von Pettenkofer Institute, Munich, Germany) | This study |
| *Enterocloster clostridioformis* SW27 | - | OMM12-colonized mouse, Facility A (Max von Pettenkofer Institute, Munich, Germany) | This study |
| *Enterocloster clostridioformis* SW28 | - | OMM12-colonized mouse, Facility A (Max von Pettenkofer Institute, Munich, Germany) | This study |
| *Enterocloster clostridioformis* SW29 | - | OMM12-colonized mouse, Facility A (Max von Pettenkofer Institute, Munich, Germany) | This study |
| *Enterocloster clostridioformis* SW30 | - | OMM12-colonized mouse, Facility A (Max von Pettenkofer Institute, Munich, Germany) | This study |
| *Enterococcus faecalis* SW141 | - | OMM12-colonized mouse, Facility A (Max von Pettenkofer Institute, Munich, Germany) | This study |
| *Enterococcus faecalis* SW143 | - | OMM12-colonized mouse, Facility A (Max von Pettenkofer Institute, Munich, Germany) | This study |
| *Enterococcus faecalis* SW144 | - | OMM12-colonized mouse, Facility A (Max von Pettenkofer Institute, Munich, Germany) | This study |
| *Enterococcus faecalis* SW146 | - | OMM12-colonized mouse, Facility A (Max von Pettenkofer Institute, Munich, Germany) | This study |
| *Enterococcus faecalis* SW149 | - | OMM12-colonized mouse, Facility A (Max von Pettenkofer Institute, Munich, Germany) | This study |
| *Enterococcus faecalis* SW151 | - | OMM12-colonized mouse, Facility A (Max von Pettenkofer Institute, Munich, Germany) | This study |
| *Enterococcus faecalis* SW152 | - | OMM12-colonized mouse, Facility A (Max von Pettenkofer Institute, Munich, Germany) | This study |
| *Enterococcus faecalis* SW156 | - | OMM12-colonized mouse, Facility A (Max von Pettenkofer Institute, Munich, Germany) | This study |
| *Enterococcus faecalis* SW157 | - | OMM12-colonized mouse, Facility A (Max von Pettenkofer Institute, Munich, Germany) | This study |
| *Enterococcus faecalis* SW159 | - | OMM12-colonized mouse, Facility A (Max von Pettenkofer Institute, Munich, Germany) | This study |

| | | | |
|---|---|---|---|
| *Enterococcus faecalis* SW160 | - | OMM12-colonized mouse, Facility A (Max von Pettenkofer Institute, Munich, Germany) | This study |
| *Enterococcus faecalis* SW320 | - | OMM12-colonized mouse, Facility A (Max von Pettenkofer Institute, Munich, Germany) | This study |
| *Enterococcus faecalis* SW321 | - | OMM12-colonized mouse, Facility A (Max von Pettenkofer Institute, Munich, Germany) | This study |
| *Enterococcus faecalis* SW322 | - | OMM12-colonized mouse, Facility A (Max von Pettenkofer Institute, Munich, Germany) | This study |
| *Enterococcus faecalis* SW323 | - | OMM12-colonized mouse, Facility A (Max von Pettenkofer Institute, Munich, Germany) | This study |
| *Enterococcus faecalis* SW324 | - | OMM12-colonized mouse, Facility A (Max von Pettenkofer Institute, Munich, Germany) | This study |
| *Enterococcus faecalis* SW325 | - | OMM12-colonized mouse, Facility A (Max von Pettenkofer Institute, Munich, Germany) | This study |
| *Enterococcus faecalis* SW326 | - | OMM12-colonized mouse, Facility A (Max von Pettenkofer Institute, Munich, Germany) | This study |
| *Enterococcus faecalis* SW327 | - | OMM12-colonized mouse, Facility A (Max von Pettenkofer Institute, Munich, Germany) | This study |
| *Enterococcus faecalis* SW328 | - | OMM12-colonized mouse, Facility A (Max von Pettenkofer Institute, Munich, Germany) | This study |
| *Enterococcus faecalis* SW329 | - | OMM12-colonized mouse, Facility A (Max von Pettenkofer Institute, Munich, Germany) | This study |
| *Enterococcus faecalis* SW331 | - | OMM12-colonized mouse, Facility A (Max von Pettenkofer Institute, Munich, Germany) | This study |
| *Enterococcus faecalis* SW161 | - | OMM12-colonized mouse, Facility B (Hannover Medical School, Hannover, Germany) | This study |
| *Enterococcus faecalis* SW163 | - | OMM12-colonized mouse, Facility B (Hannover Medical School, Hannover, Germany) | This study |
| *Enterococcus faecalis* SW166 | - | OMM12-colonized mouse, Facility B (Hannover Medical School, Hannover, Germany) | This study |

| | | | |
|---|---|---|---|
| *Enterococcus faecalis* SW168 | - | OMM12-colonized mouse, Facility B (Hannover Medical School, Hannover, Germany) | This study |
| *Enterococcus faecalis* SW170 | - | OMM12-colonized mouse, Facility B (Hannover Medical School, Hannover, Germany) | This study |
| *Enterococcus faecalis* SW172 | - | OMM12-colonized mouse, Facility B (Hannover Medical School, Hannover, Germany) | This study |
| *Enterococcus faecalis* SW174 | - | OMM12-colonized mouse, Facility B (Hannover Medical School, Hannover, Germany) | This study |
| *Enterococcus faecalis* SW176 | - | OMM12-colonized mouse, Facility B (Hannover Medical School, Hannover, Germany) | This study |
| *Enterococcus faecalis* SW178 | - | OMM12-colonized mouse, Facility B (Hannover Medical School, Hannover, Germany) | This study |
| *Enterococcus faecalis* SW179 | - | OMM12-colonized mouse, Facility B (Hannover Medical School, Hannover, Germany) | This study |
| *Enterococcus faecalis* SW180 | - | OMM12-colonized mouse, Facility B (Hannover Medical School, Hannover, Germany) | This study |
| *Enterococcus faecalis* SW264 | - | OMM12-colonized mouse, Facility C (Helmholtz Centre for Infection Research, Braunschweig, Germany) | This study |
| *Enterococcus faecalis* SW265 | - | OMM12-colonized mouse, Facility C (Helmholtz Centre for Infection Research, Braunschweig, Germany) | This study |
| *Enterococcus faecalis* SW269 | - | OMM12-colonized mouse, Facility C (Helmholtz Centre for Infection Research, Braunschweig, Germany) | This study |
| *Enterococcus faecalis* SW270 | - | OMM12-colonized mouse, Facility C (Helmholtz Centre for Infection Research, Braunschweig, Germany) | This study |
| *Enterococcus faecalis* SW271 | - | OMM12-colonized mouse, Facility C (Helmholtz Centre for Infection Research, Braunschweig, Germany) | This study |
| *Enterococcus faecalis* SW273 | - | OMM12-colonized mouse, Facility C (Helmholtz Centre for | This study |

| | | | |
|---|---|---|---|
| | | Infection Research, Braunschweig, Germany) | |
| *Enterococcus faecalis* SW275 | - | OMM12-colonized mouse, Facility C (Helmholtz Centre for Infection Research, Braunschweig, Germany) | This study |
| *Enterococcus faecalis* SW276 | - | OMM12-colonized mouse, Facility C (Helmholtz Centre for Infection Research, Braunschweig, Germany) | This study |
| *Enterococcus faecalis* SW278 | - | OMM12-colonized mouse, Facility C (Helmholtz Centre for Infection Research, Braunschweig, Germany) | This study |
| *Enterococcus faecalis* SW281 | - | OMM12-colonized mouse, Facility C (Helmholtz Centre for Infection Research, Braunschweig, Germany) | This study |
| *Enterococcus faecalis* SW283 | - | OMM12-colonized mouse, Facility C (Helmholtz Centre for Infection Research, Braunschweig, Germany) | This study |
| *Enterococcus faecalis* SW286 | - | OMM12-colonized mouse, Facility D (ETH Zurich, Zurich Switzerland) | This study |
| *Enterococcus faecalis* SW288 | - | OMM12-colonized mouse, Facility D (ETH Zurich, Zurich Switzerland) | This study |
| *Enterococcus faecalis* SW290 | - | OMM12-colonized mouse, Facility D (ETH Zurich, Zurich Switzerland) | This study |
| *Enterococcus faecalis* SW292 | - | OMM12-colonized mouse, Facility D (ETH Zurich, Zurich Switzerland) | This study |
| *Enterococcus faecalis* SW293 | - | OMM12-colonized mouse, Facility D (ETH Zurich, Zurich Switzerland) | This study |
| *Enterococcus faecalis* SW295 | - | OMM12-colonized mouse, Facility D (ETH Zurich, Zurich Switzerland) | This study |
| *Enterococcus faecalis* SW296 | - | OMM12-colonized mouse, Facility D (ETH Zurich, Zurich Switzerland) | This study |
| *Enterococcus faecalis* SW297 | - | OMM12-colonized mouse, Facility D (ETH Zurich, Zurich Switzerland) | This study |

| | | | |
|---|---|---|---|
| *Enterococcus faecalis* SW298 | - | OMM12-colonized mouse, Facility D (ETH Zurich, Zurich Switzerland) | This study |
| *Enterococcus faecalis* SW300 | - | OMM12-colonized mouse, Facility D (ETH Zurich, Zurich Switzerland) | This study |
| *Enterococcus faecalis* SW301 | - | OMM12-colonized mouse, Facility D (ETH Zurich, Zurich Switzerland) | This study |
| *Enterococcus faecalis* MS81 | - | Infant B (breast-fed) | This study |
| *Enterococcus faecalis* MS82 | - | Infant B (breast-fed) | This study |
| *Enterococcus faecalis* MS86 | - | Infant B (breast-fed) | This study |
| *Enterococcus faecalis* MS90 | - | Infant B (breast-fed) | This study |
| *Enterococcus faecalis* MS96 | - | Infant B (breast-fed) | This study |
| *Enterococcus faecalis* MS100 | - | Infant B (breast-fed) | This study |
| *Enterococcus faecalis* MS105 | - | Infant D (formula-fed) | This study |
| *Enterococcus faecalis* MS107 | - | Infant D (formula-fed) | This study |
| *Enterococcus faecalis* MS110 | - | Infant D (formula-fed) | This study |
| *Enterococcus faecalis* MS113 | - | Infant D (formula-fed) | This study |
| *Enterococcus faecalis* MS115 | - | Infant D (formula-fed) | This study |
| *Enterococcus faecalis* MS118 | - | Infant D (formula-fed) | This study |
| *Enterococcus faecalis* MS120 | - | Infant D (formula-fed) | This study |
| *Enterococcus faecalis* MS121 | - | Infant E (formula-fed) | This study |
| *Enterococcus faecalis* MS122 | - | Infant E (formula-fed) | This study |
| *Enterococcus faecalis* MS123 | - | Infant E (formula-fed) | This study |
| *Enterococcus faecalis* MS124 | - | Infant E (formula-fed) | This study |
| *Enterococcus faecalis* MS125 | - | Infant E (formula-fed) | This study |

| | | | |
|---|---|---|---|
| *Enterococcus faecalis* MS129 | - | Infant E (formula-fed) | This study |
| *Enterococcus faecalis* MS131 | - | Infant C (breast-fed) | This study |
| *Enterococcus faecalis* MS133 | - | Infant C (breast-fed) | This study |
| *Enterococcus faecalis* MS135 | - | Infant C (breast-fed) | This study |
| *Enterococcus faecalis* MS137 | - | Infant C (breast-fed) | This study |
| *Enterococcus faecalis* MS140 | - | Infant C (breast-fed) | This study |
| *Enterococcus faecalis* MS145 | - | Infant C (breast-fed) | This study |
| Enterococcus faecalis MS150 | - | Infant A (breast-fed) | This study |
| Enterococcus faecalis MS155 | - | Infant A (breast-fed) | This study |
| Enterococcus faecalis MS156 | - | Infant A (breast-fed) | This study |
| Enterococcus faecalis MS163 | - | Infant A (breast-fed) | This study |
| *Enterococcus faecalis* MS166 | - | Infant A (breast-fed) | This study |
| Enterococcus faecalis MS169 | - | Infant A (breast-fed) | This study |
| *Enterococcus faecalis* MS314 | - | Infant F (formula-fed) | This study |
| *Enterococcus faecalis* MS315 | - | Infant F (formula-fed) | This study |
| *Enterococcus faecalis* MS317 | - | Infant F (formula-fed) | This study |
| *Enterococcus faecalis* MS319 | - | Infant F (formula-fed) | This study |
| *Enterococcus faecalis* MS320 | - | Infant F (formula-fed) | This study |
| *Enterococcus faecalis* MS322 | - | Infant F (formula-fed) | This study |
| *Enterococcus faecalis* MS323 | - | Infant F (formula-fed) | This study |

**Table S2: Primers and plasmids used for construction of bacterial mutants.**

| Designation | Sequence 5´-3´ | Reference |
|---|---|---|
| ***E. faecalis* SW160 *gfrEF* deletion mutant** | | |
| pLT06 seq Fwd | GCATCCGATTGCAGTATAAATTTAAC | This study |
| pLT06 seq Rev | CCTCCTTTCTATTTGATTAGTACC | This study |
| ermR Fwd | CAGCAAAGAATGGCGGAAACG | This study |
| ermR Rev | GCGCTAGGGACCTCTTTAG | This study |
| gfrEF up region_Fwd | TCCCCGGGTACCGAGCTCGCCATGCAAGACCCTTTAGGTAG | This study |
| gfrEF up region_Rev | CGTTTCCGCCATTCTTTGCTGGCAAGCCTGTGGTAGACTG | This study |
| gfrEF down region_Fwd | CTAAAGAGGTCCCTAGCGCCCGACAATTGCCATGCCAC | This study |
| gfrEF down region_Rev | CCTGCAGGTCGACTCTAGAGGGATTTGCGATGCTTGCACG | This study |
| gfrEF P1_Fwd | CCTGGAGCTGGATAAGTCACC | This study |
| gfrEF P1_Rev | GGGCCTTATCTTGAAGCGG | This study |
| ***Salmonella gfr* deletion mutant** | | |
| gfr_donor_frwd | AGCGCCATCAGGCACTGTATATGCTAC<br>ACGCCCCTTTCTG<br>ATATGAATATCCTCCTTAGTTCC | This study |
| gfr_donor_rev | GCTCTCTTTTATCACAGTGGAATGTGTT<br>TTAGAGGTCAGG<br>TGTGTAGGCTGGAGCTGCTTC | This study |
| gfr_del_frwd | GTTAACCCGCATCTTTTTTCG | This study |
| gfr_del_rev | GATGTGTCCACTATCACGCTG | This study |
| **16S gene amplification** | | |
| fD1 | CGATATCTCTAGAAGAGTTTGATCCTGGCTCAG | Adapted from Weisburg, Barns et al. 1991 |
| fD2 | CGATATCTCTAGAAGAGTTTGATCATGGCTCAG | Adapted from Weisburg, Barns et al. 1991 |
| rP1 | GATATCGGATCCACGGTTACCTTGTTACGACTT | Adapted from Weisburg, Barns et al. 1991 |

| Designation | Description | Reference |
|---|---|---|
| **pLT06** | *E. faecalis* homologous recombination plasmid, used to create pSW13 | Thurlow et al., 2009 |
| **pKD3** | Source of chloramphenicol cassette for gfr deletion mutant in Salmonella donor strain | Datsenko et al., 2000 |
| **pSW13** | *E. faecalis* SW160 *gfrEF* deletion plasmid | This study |
| **pMPS35353** | Source of erythromycin cassette for pSW13 | Bryan et al., 2000 |

**Table S3: Composition of semi-evolved OMM$^{12}$ communities.**

| Species | Ori | Semi-evolved 1 | Semi-evolved 2 |
|---|---|---|---|
| ***M. intestinale*** | YL27 | SW13, SW15, SW16 | SW13, SW15, SW16 |
| ***B. caecimuris*** | I48 | SW203, SW210 | SW203, SW210 |
| ***C. innocuum*** | I46 | SW181, SW184, SW190 | SW181, SW184, SW190 |
| ***B. coccoides*** | YL58 | SW2, SW4 | SW2, SW4 |
| ***E. clostridioformis*** | YL32 | SW25, SW28 | SW25, SW28 |
| ***E. faecalis*** | KB1 | KB1 | SW160 |
| ***L. reuteri*** | I49 | I49 | I49 |
| ***F. plautii*** | YL31 | YL31 | YL31 |
| ***A. muciniphila*** | YL44 | YL44 | YL44 |
| ***T. muris*** | YL45 | YL45 | YL45 |
| ***A. muris*** | KB18 | KB18 | KB18 |
| ***B. animalis*** | YL2 | YL2 | YL2 |

| |
|---|
| **Unevolved ancestor** |
| **Evolved isolate** |

**Table S4: Mouse diet and sterilization method used in the different animal facilities.**

| Facility | Location | Diet type | Diet sterilization method | Diet producer | Product number |
|---|---|---|---|---|---|
| **MUC** | Max von Pettenkofer Institute Munich | Mouse breeding diet | Autoclaving | Ssniff, Germany | V1124-300 |
| **HAN** | Hannover Medical School | Mouse breeding diet | 50 kGy gamma-irradiation | Ssniff, Germany | V1124-927 |
| **BS** | Helmholtz Center for Infection Research Braunschweig | Mouse breeding diet | 50 kGy gamma-irradiation | Ssniff, Germany | V1124-927 |
| **ZUC** | ETH Zurich | Mouse breeding diet | Autoclaving | Kliba-Nafag, Switzerland | 3807 |
